# A novel dual enrichment strategy provides soil- and digestate-competent N_2_O-respiring bacteria for mitigating climate forcing in agriculture

**DOI:** 10.1101/2021.05.11.443593

**Authors:** Kjell Rune Jonassen, Ida Ormaasen, Clara Duffner, Torgeir R Hvidsten, Åsa Frostegård, Lars R Bakken, Silas HW Vick

## Abstract

Manipulating soil metabolism by heavy inoculation with microbes is deemed realistic if waste from anaerobic digestion (digestate) is utilized as substrate and vector, but requires organisms that can grow both in digestate and soil (=generalist). We designed a strategy to enrich and isolate such generalist N_2_O-respiring bacteria (NRB) in soil and digestate, to provide inoculum for reducing N_2_O-emissions from agricultural soil. Sequential anaerobic enrichment cultures were provided with a small dose of O_2_ and unlimited N_2_O, alternating between sterilized digestate and soil as substrates. The cultures were monitored for gas kinetics and community composition (16SrDNA), and cluster-analysis identified generalist-OTUs which became dominant, digestate/soil-specialists which did not, and a majority that were diluted out. Several NRBs circumscribed by generalist-OTU’s were isolated, genome sequenced to screen for catabolic capacity, and phenotyped, to assess their capacity as N_2_O-sinks in soil. The two isolates *Cloacibacterium* sp., carrying only N_2_O-reductase (Clade-II) and *Pseudomonas sp*., with full-fledged denitrification-pathway, were both very effective N_2_O-sinks in soil, with *Pseudomonas* sp., showing a long-lasting sink effect, suggesting better survival in soil. This avenue for utilizing waste to bioengineer the soil microbiota holds promise to effectively combat N_2_O-emissions but could also be utilized for enhancing other metabolic functions in soil.

**Graphical abstract:** 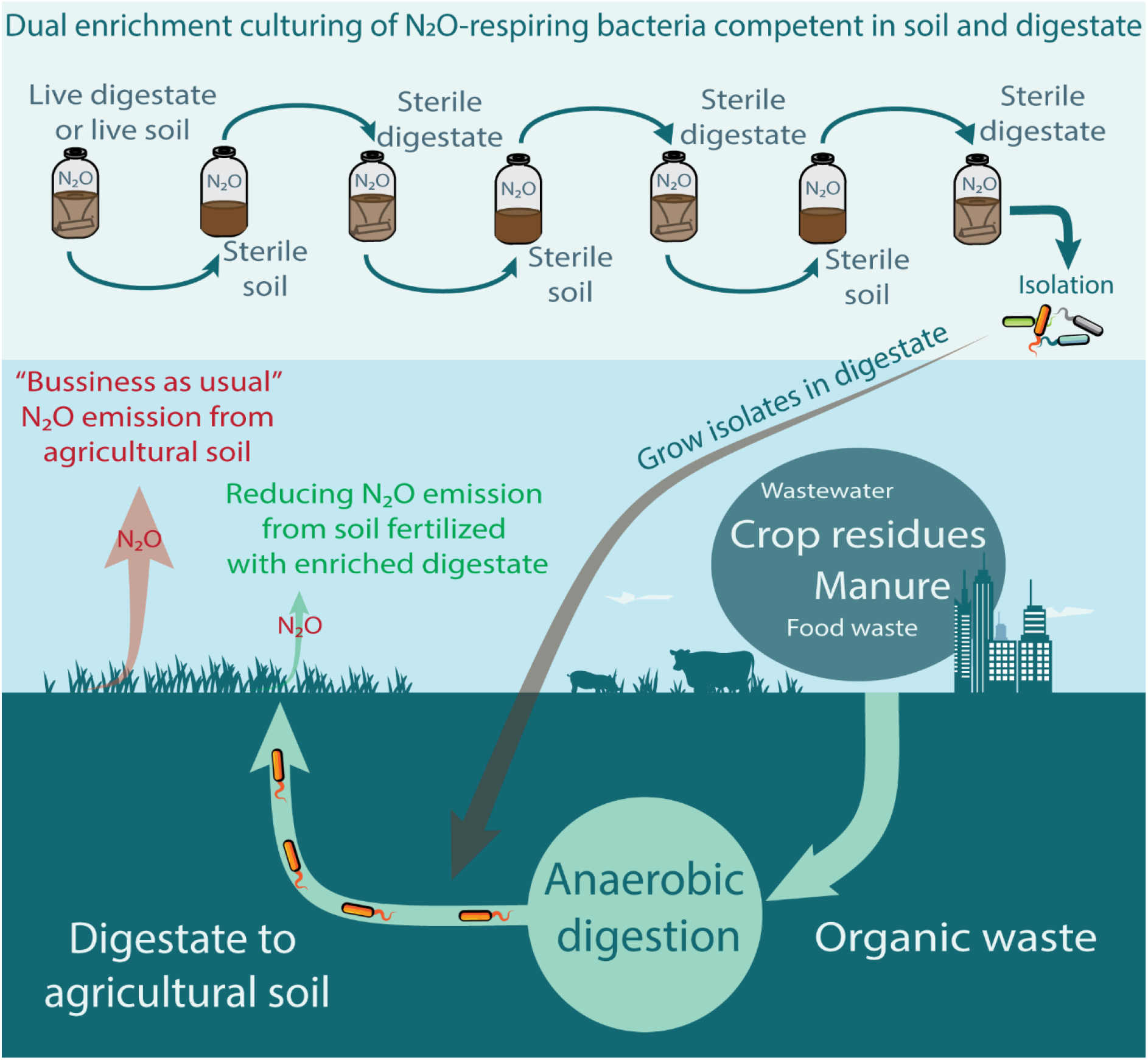

## Introduction

The N_2_O-concentration in the atmosphere is increasing, largely driven by the input of reactive nitrogen species in agriculture (Davidson 2009, Thompson et al 2019). N_2_O-emissions from farmed soils account for 52 % of the total anthropogenic emissions of N_2_O (Tian et al 2020) and approximately 1/3 of the climate forcing from food production (Robertson 2014). Limiting the input of reactive nitrogen to soils would be an effective mitigation measure but at the expense of lowering crop yields. This dichotomy has proven difficult to bypass, and estimates indicate only modest N_2_O mitigation potentials if currently available N_2_O abatement options were to be implemented at large scale (Winiwarter et al 2018).

In agricultural soils nitrification and denitrification are the main sources of N_2_O (Butterbach-Bahl et al 2013). Nitrous oxide reductase (Nos) is the only known enzyme catalyzing the reduction of N_2_O. Nos is expressed as part of the denitrification pathway sustaining anaerobic respiration by stepwise reduction of NO_3_^−^ → NO_2_^−^ → NO → N_2_O → N_2_, catalyzed by the enzymes nitrate reductase (Nar), nitrite reductase (Nir), nitric oxide reductase (Nor) and nitrous oxide reductase (Nos) encoded by the genes *nar/nap, nir, nor* and *nosZ*, respectively (Zumft 2007). A significant share of denitrifying prokaryotes, however, are truncated, i.e. lacking genes encoding for 1 to 3 of the four enzymes (Shapleigh 2013; Graf et al 2014), and truncated denitrifying pathways may significantly affect the N_2_O-emissions in soils under denitrifying conditions. Organisms that lack all denitrification genes other than *nosZ* are particularly interesting as they can act as net sinks for N_2_O. The propensity of the soil community to emit N_2_O can be reduced by increasing the relative abundance of such N_2_O-respiring bacteria (NRB) (Philippot et al 2011, Domeignoz-Horta et al 2016). However, as a stand-alone operation, such modification of soil microflora by inoculation would be prohibitively expensive.

We have previously demonstrated that anaerobic digestion (AD) provides a promising industrial platform for low-cost large-scale introduction of N_2_O-reducing bacteria to soil (Jonassen et al 2021): denitrifying bacteria with a strong preference for N_2_O over NO_3_ were isolated from an AD-digestate, which could be grown aerobically to high cell densities in a sterilized digestate, providing a cheap inoculum for reducing N_2_O-emission from soil. The isolated organisms did not include NRB (bacteria with only *nosZ*), however, and it was evident that the organisms were not well adapted for activity and survival in soil. Here we present a new approach to obtain more ideal isolates by a deliberate attempt to enrich (and isolate) organisms that can grow both in digestate and soil. Conceptually the N_2_O-reducing organisms within a community can be divided into three categories according to their ability to grow/survive in digestates and soil: *digestate specialists* (***D***) with a competitive advantage in digestate, *soil specialists* (***S***) with a competitive advantage in soil, and *generalists* (***G***) organisms capable of growth in both environments, but plausibly at lower growth rates in both substrates relative to the two specialists. We hypothesized that we could enrich ***G*** by sequential enrichment culturing, alternating between soil and digestate as substrate (coined *dual enrichment*), and explored this with a logistic growth model for the competition between three organisms, assigning hypothetical growth and death rates. The model revealed that the selective pressure could be modulated by the duration of each enrichment and the fraction of enriched material transferred from one enrichment to the next, and predicted that a reasonably competitive *generalist* would reach dominance after a limited number of repeated passages.

Using this theoretical framework, we designed an enrichment strategy whereby a microbial community, originating from digestate or soil, was passaged through a series of enrichment cultures alternating between gamma sterilized soil (γ-soil) and autoclave sterilized digestate (AC-digestate) (**Fig. 1**). We anticipated that *generalists* would gradually increase in abundance throughout the enrichment series and that organisms that are non-competitive in either substrates would be washed out due to the repeated dilution each transfer represented. Strong specialists would likely reappear when reintroduced in their preferred environment, and thus be easily identifiable. By means of this novel enrichment strategy along with targeted isolation of N_2_O respiring isolates, genome sequencing and physiological experiments designed to unravel the isolates’ denitrifying regulatory phenotypes, we provide insight into the targeted enrichment of generalist type organisms and their performance as N_2_O mitigating inoculants when vectored by digestate to agricultural soil.

**Figure 1:**
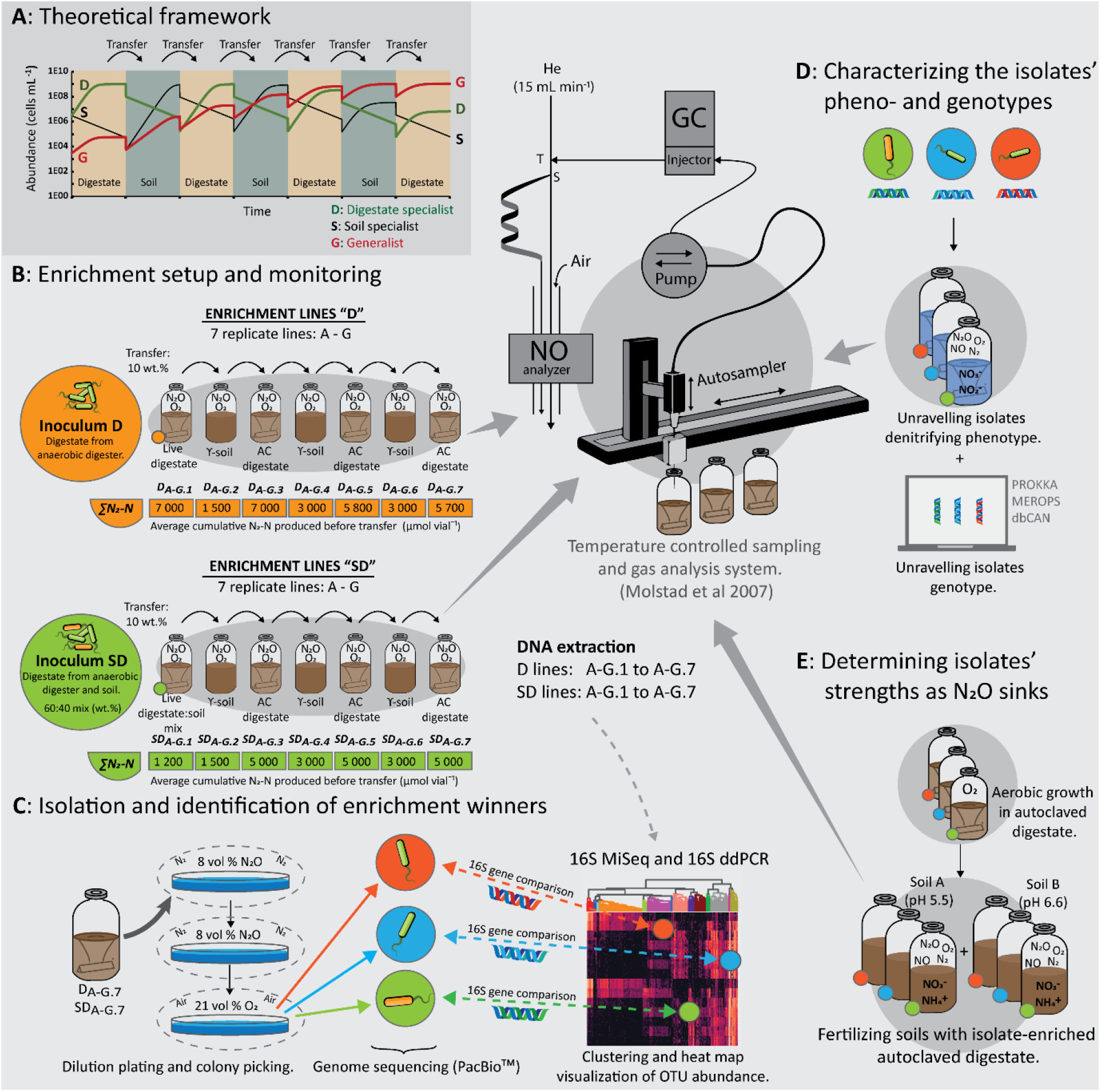
Graphical summary of materials and methods. **A:** The dual enrichment was modelled by a set of Lotka-Volterra logistic equations for three organisms; digestate specialist (D), soil specialist (S), and generalist (G), competing for a common substrate pool. The repeated transfer of enriched material from one enrichment to the next, alternating between soil and digestate, was predicted to enrich generalists by nature. The modelling is presented in detail in **Supplementary materials Section 1**: **Figs. S1** to **S6** and **Tab. S1**). **B**: Enrichment culturing experimental setup for the two enrichment lines “D” (digestate derived inoculum) and “SD” (soil and digestate derived mixed inoculum), each consisting of seven parallel replicate lines (A - G) over seven transfers. Each batch was supplemented with O_2_ and N_2_O (He background) in the headspace and monitored for O_2_, N_2_O, and N_2_ kinetics by frequent sampling of the headspace. While O_2_ was allowed to deplete by respiration, N_2_O was sustained throughout by repeated injections. Average cumulative N_2_ produced for each culture is indicated below vials (∑N_2_-N). DNA was extracted from every vial at the conclusion of each enrichment. **C**: Extracted DNA was subjected to 16S rDNA amplicon sequencing, OTU clustering, and taxonomic assignment. The abundance of organisms circumscribed by each OTU was calculated from their relative abundance and the abundance of 16S rDNA mL^−1^ as measured with digital droplet PCR (ddPCR). Relative abundance of the 500 most abundant OTUs throughout the enrichment was clustered using the Ward variance minimization algorithm. This allowed for identification of clades of OTUs with similar development throughout the dual enrichments. The OTUs 16S consensus sequences were aligned and matched with the 16S genes recovered from full genome sequencing of axenic N_2_O-reducing isolates obtained from the final enrichments. **D**: The isolates’ denitrifying phenotypes were assessed in pure culture incubations supplemented with either NO_3_^−^ or NO_2_^−^, and O_2_ or N_2_O and O_2_ and their phenotype matched against their denitrifying genotypes. Eco-physiological genome analysis by annotation of carbohydrate-active enzymes, peptidases, denitrification reductase genes, and other genes provided insight into the suitability of these isolates as N_2_O-reducing inoculants for soil inoculation. **E**: Each isolate was grown aerobically to high cell densities in autoclaved aerated digestate before amendment in two live soils (Soil A: pH 5.5 and Soil B: pH 6.6) supplemented O_2_ and NO_3_^−^ to assess performance as N_2_O-reducing inoculants in soil.

## Materials and methods

### Incubation- and gas measurement

All incubations were done in 120 mL serum vials sealed with butyl-rubber septa, using a robotized incubation system (Molstad et al 2007, 2016) which monitors gas kinetics (O_2_, N_2_, N_2_O, NO, CO_2_ and CH_4_) by repeated sampling of the headspace, returning an equal volume of He each time. Elaborated calculus routines, accounting for dilution by sampling and leakage (Bakken 2021) secures accurate estimates of production/consumption rates of each gas, electron flow rates to the various electron acceptors (O_2_, NO_3_^−^, NO_2_-, NO, N_2_O), and N mass-balance. Digestates and liquid cultures of isolated strains were stirred continuously (600 rpm) with a 23 mm Teflon-coated triangular magnet. Prior to incubation, the headspace air was replaced with He by repeated evacuation and filling with He and supplemented with pure N_2_O and/or O_2_ (Molstad et al 2007).

### Digestate and soils

The digestate was taken from the anaerobic digester of a municipal WWTP (Jonassen et al 2020), with chemical characteristics given in **Tab. S2**. Two clay loam soils were taken from a long-term liming experiment (Nadeem et al 2020), one with pH_CaCl2_=6.6 (Soil A), and one with pH_CaCl2_=5.5 (Soil B). Live digestate and live soil A were used in the initial enrichment cultures, while the substrates for subsequent enrichments (**Fig. 1B**) were autoclaved digestate (pH adjusted to 7.2 by titration with HCl), and γ-irradiated Soil A (25.9 kGy, 12 months prior to experiments). Digestate used for aerobic growth of the isolated N_2_O-reducing bacteria before soil amendments (**Fig. 1E**) was autoclaved (121 °C, 20 min), then aerated by pumping sterile filtered air through a stirred suspension of digestate for 36 hours, and then pH adjusted to ~7.50-7.75 (**Tab. S2**) by titration with 4 M HCl. Aeration of the digestate was necessary in order to oxidize Fe^2+^ in the digestate to Fe^3+^, as otherwise the abiotic reduction of O_2_ by Fe^2+^ obscured measurements of oxygen consumption (Jonassen et al 2021).

### Dual enrichment culturing

Enrichment series were started with two live materials: 50 mL digestate (**D-lines**) (pH 7.6±0.1) and 20 g Soil A + 30 mL digestate (**SD-lines**) (pH 7.2±0.1), each with 7 independent lines (A to G) (**Fig. 1A**). The nomenclature used throughout the text is: *D_A-G,j_* and *SD_A-G,j_*, where D/SD denotes the initial live materials, *A-G* denotes the 7 independent replicates and *j* the enrichment number (1–7). D_0_/SD_0_ denotes live material before enrichment with N_2_O. After replacing the headspace air with He, 3 mL N_2_O, and 3 mL O_2_ were injected into the vials, which were then incubated at 20 °C in the incubation system monitoring the O_2_, N_2_O and N_2_. Additional N_2_O was injected when needed to avoid N_2_O depletion. Subsequent enrichment cultures (j=2-7), alternating between γ-sterilized soil (45 g soil dry weight vial^−1^+ 16 mL sterile water) autoclaved digestate (45 mL), were inoculated with ~10 wt% of the previous enrichment, following the same experimental procedure and conditions as explained above for the live starting materials. At the completion of each enrichment, samples were taken for DNA extraction and analysis and for isolation in the final enrichment.

### Community analysis

DNA was extracted from technical duplicates, sampled at the conclusion of each enrichment cycle, from all D_A-G,j_ and SD_A-G,j_ vials (j = 1-7), the live materials (D_0_ and SD_0_), autoclaved digestate and γ-soil. DNA was extracted from 1 mL digestate slurry or 0.25 g soil using the PowerLyzer™ Soil DNA extraction kit (QIAGEN) following a modified kit protocol where bead beating for 30s at 4.5 ms^−1^ in a MP Biomedicals™ FastPrep^®^-24 (Thermo Fischer Scientific Inc) replaced the vortexing step in the manufacturers protocol. Quantitative digital droplet PCR (ddPCR) was performed in technical triplicates on pooled samples of DNA extracts from biological and technical replicates from each enrichment cycle (j = 1-7), and pooled samples of technical replicate DNA extractions from D_0_/SD_0_, autoclaved digestate, and γ-soil, respectively. The ddPCR reaction mix (QX200 ddPCR EvaGreen^®^ Supermix, BioRad) was prepared according to the manufacturer’s instructions using the universal primers PRK341F (5’-CCTACGGGRBGCASCAG-3’) and PRK806R (5’-GGACTACYVGGGTATCT-3’) (Eurofins Genomic) targeting the V3-V4 region of the 16S rDNA gene (Yu et al 2005). The QX200 droplet generator (Bio-Rad) was used to generate oil droplet suspensions that were subjected to PCR with parameters given in **Tab. S3**. The PCR products were measured in a QX200 droplet reader (Bio-Rad), and the data analyzed in Quantasoft™ Analysis Pro 1.0.596 software (Bio-Rad). Microbial community composition was determined through 16S rDNA amplicon sequencing (V3-V4 region) and taxonomic classification of 16S rRNA gene sequences. Library preparation and sequencing data processing were performed according to Nilsen et al (2020) except the library was quantified with the KAPA library quantification kit (universal; Roche) in a CFX96 Touch Real-Time PCR Detection System (Bio-Rad, USA). The amplicon library was diluted to 7 pM containing 20 % PhiX before sequencing on the MiSeq platform (Illumina, USA) using MiSeq reagent v3 kit to generate 300 bp paired-end reads. The sequencing produced 6139309 reads after quality filtering. The samples were rarefied at 9000 reads, resulting in the loss of 9 samples (SD_2.1_-A, SD_7.6_-B, SD_3.7_-A, SD_7.7_-A, D_1.1_-A, D_1.3_-B, D_2.3_-A, D_2.4_-A, D_3.6_-A). The seaborn.clustermap in the Seaborn software suite (Waskom 2020) was used to generate hierarchically clustered heatmaps and was based on the Ward variance minimalization linkage algorithm (Ward 1963) for the 500 most abundant OTUs (sum abundance across all samples). Statistical analysis included principal component analysis (PCA) and similarity percentage (SIMPER) (Clarke 1996) using the PAST software (Hammer et al 2011). OTU absolute abundance was calculated as the product of its relative abundance and the abundance of 16S rDNA assessed by ddPCR for all OTUs (16S rDNA copies enrichment-vial^−1^).

### Isolation and characterization of N_2_O-reducing organisms

Dilution series of the final enrichments (D_A-G,7_ and SD_A-G,7_) were spread on Sistrom’s succinate medium (SS), R2-A, tryptic soy broth (TSB) and Nutrient broth (NB) agar plates (1.5 wt. %) (media composition is given in **Supplementary Materials Section 2**), incubated in N_2_+ N_2_O atmosphere as described in Jonassen et al (2021) (**Fig. 1C**). Colonies were transferred to 120 mL vials containing 50 mL of the corresponding liquid medium and incubated aerobically with stirring (700 rpm) at 20 °C. 16S gene analysis showed that several different isolates were obtained, six of which were selected for full genome sequencing (working names in **bold**): *Aeromonas* sp. **AM**, *Ochrobactrum* sp. **OB**, *Pseudomonas* sp. **PS-02** and *Brachymonas* sp. (**BM**), which were isolated on SS medium, and *Cloacibacterium* sp. (**CB-01** and **03**) isolated on NB medium. Cultures were grown aerobically at 20 °C in SS (**AM, OB, BM, PS-02**) or NB (**CB-01**, **CB-03**) medium to OD_660_ ≈ 1.0. After centrifugation, DNA was extracted from the pellets using PowerLyzer™ Soil DNA extraction kit (QIAGEN) as described above. The genomic DNA was sheared to approximately 8-14 kb long fragments and a library was generated with the SMRTbell Express Template Prep Kit 2.0 (PacBio) without size selection. The library was sequenced on a PacBio™ SMRT cell with the PacBio™ Sequel System using 3.0 chemistry at the Helmholtz Centre, Munich. After data demultiplexing, the genomes **CB-01**, **CB-03, BM**, and **AM** were assembled with the ‘HGAP4’ pipeline (SMRT Link Software, PacBio) with a seed coverage of 30 for **CB-01**, **CB-03**, and **BM**, and a seed coverage of 22 for **AM. PS-02** and **OB** were assembled with the ‘Microbial Assembly’ pipeline (SMRT Link Software, PacBio) with a seed coverage of 20 and 15, respectively. Genome quality was assessed with CheckM v1.0.18 (Parks et al 2015). Annotation of coding genes was done with Prokka v1.14.5 (Seemann et al 2014) using default parameters. The draft genomes were functionally annotated for carbohydrate-active enzymes (dbCAN2 meta server, Zhang et al 2018) and peptidases (MEROPS database, release 12.3) (Rawlings et al 2010). Signal P 5.0 (Raut et al 2021) was used to identify genes containing putative signal peptides as defined for gram negative bacteria. The denitrification regulatory phenotypes of the isolated strains was investigated by monitoring the kinetics of O_2_, NO, N_2_O, NO_2_^−^ and NO_3_^−^ in stirred batch cultures as they depleted the oxygen and switched to anaerobic respiration as described by Jonassen et al (2021). 200 μL samples of liquid culture were taken for NH_4_^+^ measurements and immediately stored at −20 °C before colorimetric analysis in LCK303 cuvettes (Hach Lange) in a DR 3900 spectrophotometer (Hach Lange).

### Evaluation of N_2_O-reducing isolates as N_2_O sinks in soil

The soils A and B were amended with five variations of digestate; **1:** live digestate (directly from the anaerobic digester), **2:** digestate heat treated to 70 °C for two hours, **3:** autoclaved pH-adjusted (7.75) digestate, **4:** autoclaved, aerated, and pH adjusted (7.75) digestate in which the isolates AM, BM, PS-02, CB-01 or OB had been grown by aerobic respiration, **5:** as 4, with CB-01, then heated to 70 °C (2 h). Each variation was tested in duplicate 120 mL vials containing 10 g soil (Soil A or Soil B) (**Fig. 1E**) amended with 0.6 mL digestate (1-5) and 50 μmol NO_3_^−^ and 1 mL O_2_ in a He atmosphere. Sterilized water was added to adjust the soil WFPS to 62 ± 1 % (Franzluebbers 1999). The vials were incubated at 20 °C, and monitored for O_2_, NO, N_2_O and N_2_ (**Fig. 1D**). A follow-up experiment with the same experimental design was performed to test the dose dependence effect for three of the isolates. Finally, we tested the persistence of the strains in soil, by making an identical extra set of vials (**1-5** above) which were stored aerobically in moist chambers, then amended with 1 mg g^−1^ soil ground plant material (clover) to secure high metabolic activity, and incubated as described above.

To assess the effect of isolates on the potential N_2_O emission from denitrification in soil, we used the N_2_O-index, ***I_N2O_*** (Liu et al 2014), which is the integral of the N_2_O-curve divided by the integral of the total N-gas, for a given period (0-*T*):

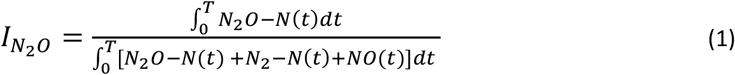

The time period (*T*) is not fixed but set as the time when a given percentage of the available nitrogen oxyanions (NO_3_^−^+NO_2_^−^) are reduced to N-gas (N_2_+N_2_O+NO). In our case, we calculated ***I_N2O_*** for 40% and 100% recovery of nitrogen oxyanions as N_2_+N_2_O+NO (coined ***I_N2O40%_*** and ***I_N2O100%_***, respectively).

### Data availability

The sequencing data for this study have been deposited in the European Nucleotide Archive (ENA) at EMBL-EBI under accession number PRJEB44171 (https://www.ebi.ac.uk/ena/browser/view/PRJEB44171).

## Results and discussion

### Dual enrichment culturing

To enrich and isolate N_2_O-respiring organisms which can grow both in digestate and soil environments, we used a dual enrichment approach, i.e. sequential batch cultures, alternating between sterile digestate and sterile soil as substrates (**Fig. 1**). Each batch was provided with a small dose of O_2_, to suppress obligate anaerobic organisms and select organisms capable of rapid transition from O_2_- to N_2_O-respiration. Subjecting the enrichments to recurrent changes (i.e. growth substrate, oxic/anoxic) selects for organisms with a capacity to adapt rapidly to changing environmental conditions (Brooks et al 2011), a desirable trait in an organism destined for soil amendment.

The kinetics of N_2_O reduction to N_2_ throughout the consecutive enrichments is shown in **Fig. 2A** (more detailed analyses of the gas kinetics are shown in **Figs. S7** and **S8**). In the line D enrichment, which started from live digestate only (D_A-G.1_), the N_2_-kinetics indicated the presence of two populations of N_2_O-respiring organisms; one whose activity was gradually declining, indicated by declining N_2_-rates (μ = −0.03 h^−1^), and a second population growing from initially extremely low numbers until their N_2_O respiration exceeded that of the declining population, increasing at a rate of 0.1 h^−1^ (modelled in **Fig. S8**, top right panel). In contrast, theline SD enrichment, which started with a mixture of live soil and live digestate (SD_A-G.1_) showed exponentially increasing rates for N_2_ production initially. Interestingly, the rates of N_2_-production in SD_A-G.1_ did not reach as high as D_A-G.1_ (~10 vs ~120 μmol N_2_-N h^−1^ vial^−1^), which could be taken to suggest that a) the N_2_O-reducing organisms originating from the soil quickly reached dominance due to the high initial numbers, b) these were less capable of scavenging electron donors in the digestate than the organisms originating from the digestate itself and/or c) the indigenous digestate bacteria were suppressed by the soil bacteria. Throughout the subsequent enrichments, the N_2_-kinetics of the SD and D line became more similar, characterized by a short exponentially increasing rate, and subsequent more or less stable plateaus, probably reflecting an early depletion of the most easily available carbon substrates. The seven-replicate series within each line (D and SD) had remarkably similar kinetics, reflected in the marginal standard deviation (**Fig. 2A**).

**Figure 2:**
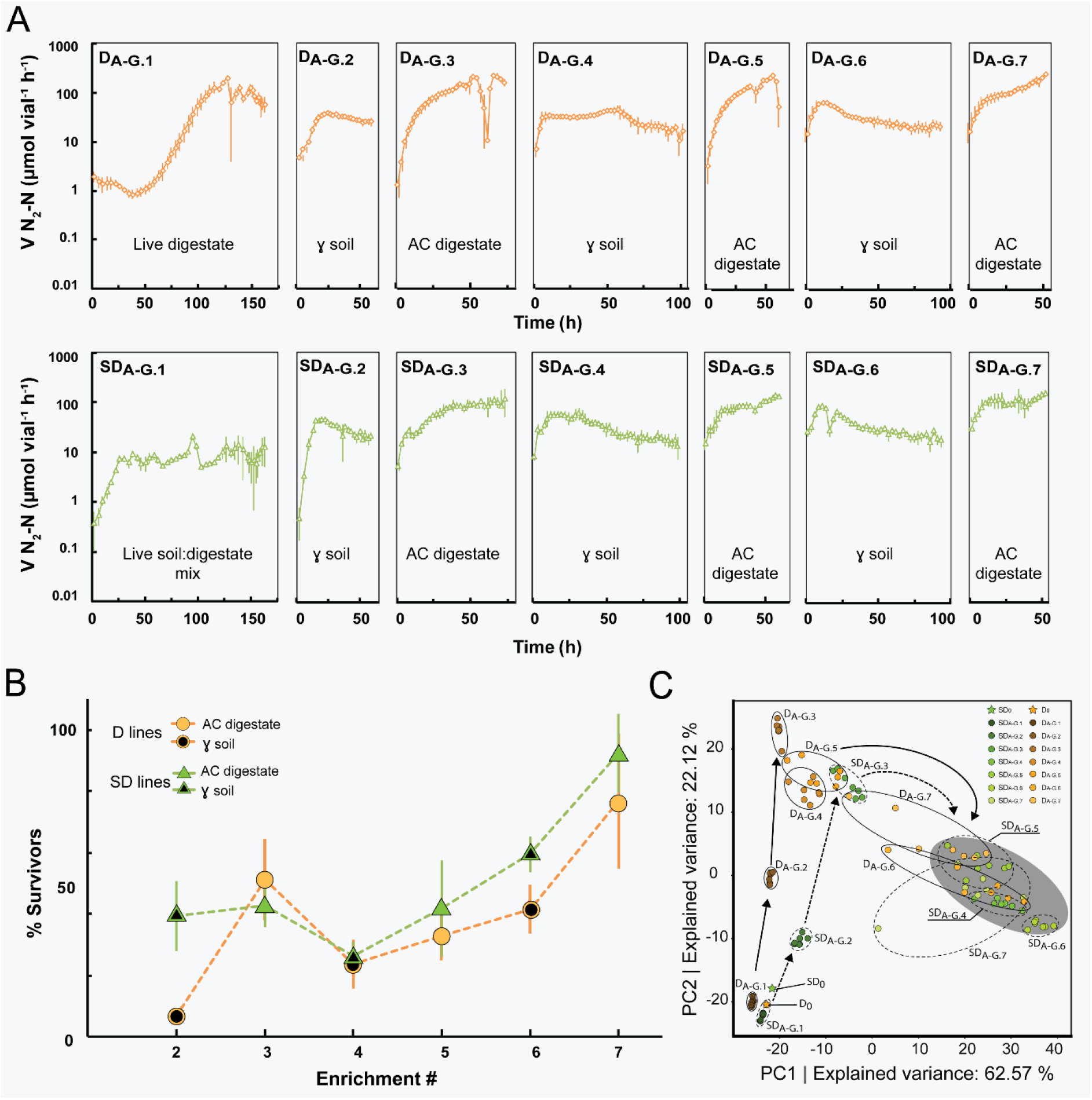
Gas kinetics and PCA of enrichment cultures. **Panel A:** Average rate of N_2_-N production for the two lines of enrichment culturing. 10 weight % of enriched material was transferred from one replicate vial to the next (D_A-G.j_ and SD_A-G.j_ to D_A-G.j+1_ and SD_A-G.j+1_). AC digestate = autoclaved. γ soil = gamma sterilized. **Panel B**: Assessment of the fraction of the community surviving transfer to the next enrichment cycle (details in **Fig. S7**). **Panel C**: PCA of OTU relative abundances. Each dot represents an individual replicate (A-G). Standard deviation (n=7) is shown as vertical bars (panels A and B).

In theory (see supplementary **Figs. S1-S6**), the dual enrichment culturing should select for organisms that are able to grow both in soil and digestate (generalists **G**) over the organisms that can only grow in soil (soil specialists, **S**) and digestate (digestate specialists **D**), leading to a gradual increase in the G/(S+D) abundance ratio, which means that the percentage of N_2_O-respiring cells that survive the transfer to a new substrate (from soil to digestate and *vice versa*) should increase. We achieved crude estimates of the % survivors for each transfer based on the cumulated N_2_ in each enrichment and the initial rates in the next (explained in detail in **Fig. S9**), and the results (**Fig. 2B**) lend support to the hypothesis.

### Microbial community development in enrichment cultures

The microbial community dynamics were analyzed based on 16S rRNA amplicon sequencing and OTU clustering. PCA of community profiles demonstrated close similarity between replicate vials (A-G) throughout the first three enrichments, and some divergence thereafter (**Fig. 2C**). SIMPER analysis revealed that 10 OTUs accounted for 94.4 and 93.5 % of the explained variance in the D and SD line respectively, of which 8 OTUs were shared between the two lines (**Tabs. S4** and **S5**). The D and SD lines followed similar trajectories and clustered in proximity to each other from enrichment SD_4_ and D_6_ forward, indicating a convergence towards a similar community structure (grey circle in **Fig 2C**). The PCA clearly verified that the community underwent continuous dynamic succession and, surprisingly, that a high fraction of dominant OTUs were shared between the two lines.

By targeted isolation of N_2_O-respiring bacteria from the final enrichment cycle in autoclaved digestate (D_A-G.7_ and S_DA-G.7_), we obtained seven axenic N_2_O-respring cultures and sequenced the genomes of six, using the PacBio sequencing platform (**Tab. S6**). The isolates were named according to genera with which they clustered in the phylogenetic tree generated with the 16S rDNA gene sequences of the isolates and related strains (**Fig. S10**) and given working names (**bold**): *Pseudomonas* sp. **PS-02**, *Aeromonas* sp. **AM**, *Brachymonas* sp. **BM**, *Ochrobactrum* sp. **OB**, *Cloacibacterium* sp. **CB-01**, *Cloacibacterium* sp. **CB-03**, and *Azonexus* sp. **AN**. **AN** was not genome sequenced, as its 16S partial sequence (obtained from Sanger sequencing of 16S PCR amplicons using 27F/1492R primer pairs) matched the 16S gene (99.2 % sequence identity) of the dominating N_2_O-reducing *Azonexus* sp. (ERR4842639) isolated and characterized in the aforementioned experiments of Jonassen et al (2021).

The 16S genes recovered from the annotated genomes were compared to 16S of the OTUs using usearch.global (Edgar 2010). This revealed high sequence identity (>97 %) in the overlapping region (404 −429 bp) of the 16S rDNA consensus sequence of some OTUs, hence these OTUs circumscribed the isolated species (**Tab. S7**). The isolates CB-01, CB-03, AN, BM and PS-02 were circumscribed by OTU1, OTU1, OTU2, OTU8 and OTU19, respectively. These OTUs represented four of the top six most abundant OTUs of the D_A-G.7_ and SD_A-G.7_ samples. Including OTU74, circumscribing the isolate **OB**, five of the top 15 OTUs circumscribed the isolates. Summed, the average abundances of these OTUs were 59.8 ± 1.2 % and 60.0 ± 1.1 % in the D_A-G.7_ and SD_A-G.7_ enrichments, of which the dominating OTU1 accounted for 33 ± 10 % and 39 ± 10 % of the total abundance.

The dynamic change in OTU abundance of the 500 most abundant OTUs (sum abundance across all samples) throughout the consecutive enrichments of the D and SD lines was hierarchically clustered based on Euclidian distance measures using the Ward’s linkage algorithm (Ward 1963) and visualized by heatmapping of OTU abundance (**Fig. 3A**). The hierarchical clustering identified six clades, denoted A to E in **Fig. 3A**, that clustered OTUs according to their abundance patterns throughout the consecutive enrichments. To achieve a more quantitative assessment of the phenomena portrayed in the heatmap we combined the total 16S rDNA gene abundance (**Tab. S8**) with the relative abundance of each clade and individual OTUs (**Fig. 3BCD**). This analysis included an assessment of the relative increase of individual OTUs from the consecutive enrichment cultures calculated as *R_i_* = ln(N_(i)_/(N_(i-1)_ · 0.1)), where N_i_ is the estimated absolute abundance at the end of enrichment *i* and N_(i-1)_ is the estimated absolute abundance at the end of the foregoing enrichment. The average *R_i_* for soil (*R_soil_*) and for digestate-enrichments (*R_Digestate_*) for each OTU was used to judge whether the OTU is a soil specialist (high *R_Soil_*, low/negative *R_Digestate_*), a generalist (high *R_Soil_* and *R_Digestate_*) or a digestate specialist (high *R_Digestate_*, low/negative *R_Soil_*).

**Figure 3:**
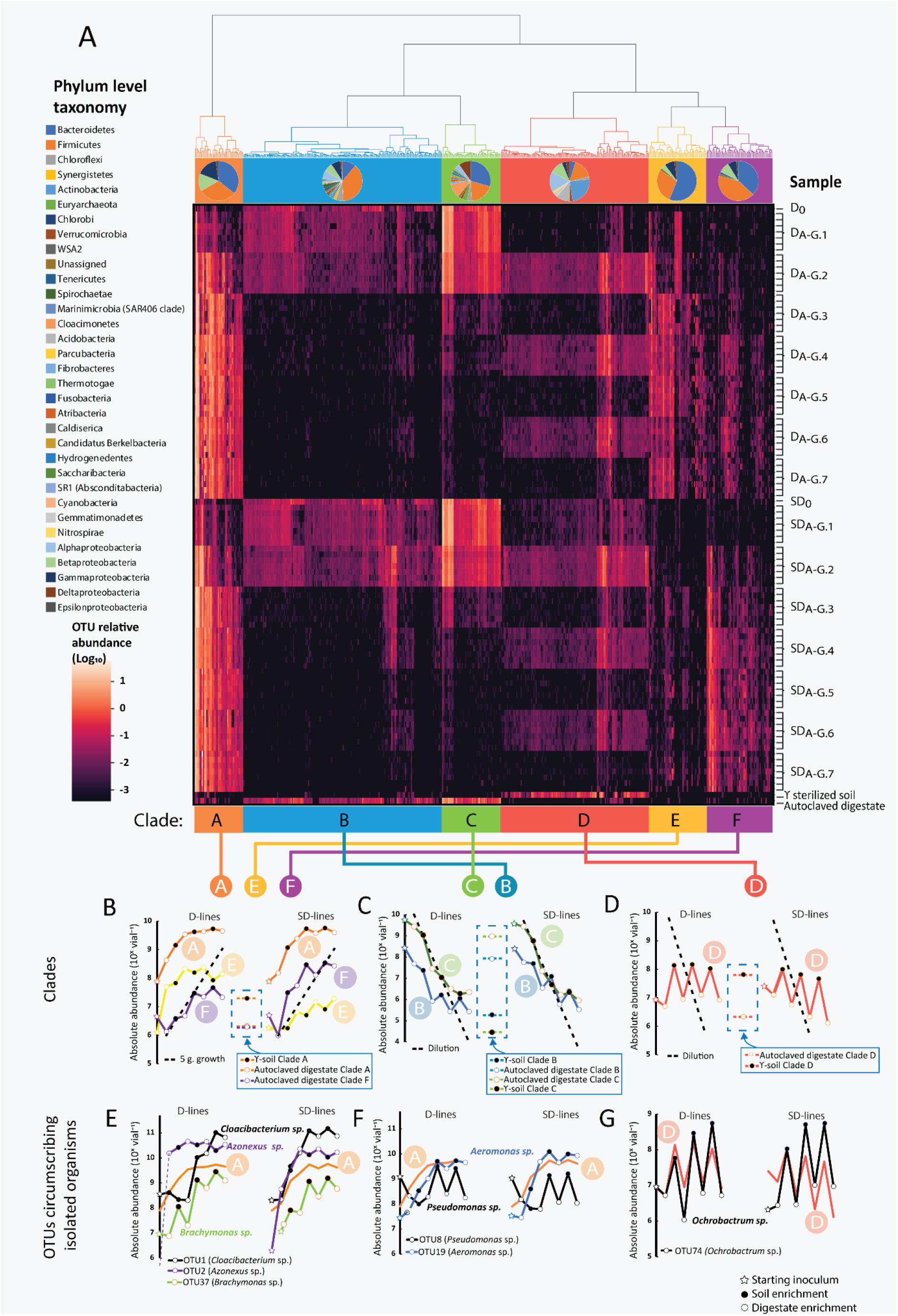
Abundance of clustered OTU’s throughout the dual enrichment culturing. **Panel A:** Heatmapping and hierarchical clustering of the 500 most abundant OTUs from all biological replicates of the D line and SD lines of the dual enrichment culturing, including starting inocula (D_0_ and SD_0_) and background samples of γ-soil and autoclaved digestate used as growth medium in the enrichments. OTUs are arranged in columns and samples in rows. The clustering has been delineated into six clades (A, B, C, D, E, F) with phylogenetic composition of OTUs in clades displayed below the cladogram. **Panels B-D** show the average absolute abundances (copies vial^−1^=) for the OTU’s within each clade throughout each enrichment; filled symbol= enrichment in soil, open symbol= enrichment in digestate, star = starting inoculum. The dashed lines in panels **C** and **D** represents the predicted decline by dilution, given a 10 % transfer rate, i.e. neither growth nor death. The dashed line in panel **B** represents a growth rate of 5 generations per enrichment. The OTU-abundancies in sterile materials are shown within dashed frames. **Panels E-G** shows the abundance of the OTU’s which circumscribe the isolated strains, together with the averages of their resident clades.

Most OTUs within Clade A were present initially in both enrichment lines (D_0_ and SD_0_), suggesting a primarily digestate origin of these OTUs, of which most were assigned to the phyla *Bacteriodetes, Cloacimonetes* and *Betaproteobacteria* (**Fig. 3A**). Clade A showed an increase in abundance throughout the enrichment in both enrichment lines (**Fig. 3B**) with an increase equivalent to ~5 cell divisions in the first 3-4 enrichment cultures (dashed line, **Fig. 3B**). Inspection of the growth of individual OTU’s (*R_i_* values) within Clade A showed that they were able to grow both in digestate and soil, but they span a range from soil specialists (*R_soil_* close to zero) to generalists (R_soil_ and R_digestate_ >2, **Fig. S11**). The OTUs circumscribing the isolated cultures **CB-01** (OTU1), **CB-03** (OTU1), **AN** (OTU2), **PS-02** (OTU8), **AM** (OTU19) and **BM** (OTU37) were all within Clade A (**Fig. 3EF**). OTU2, circumscribing *Azonexus sp*. **AN**, grew better in digestate than in soil (*R_Digestate_* 3.40 ± 0.35 and R_Soil_ 2.27 ± 0.35) and reached dominance in the first enrichment in live digestate (D_A-G.1_ culture vials), which was also observed in the enrichments of Jonassen et al (2021).

Clade B and C plausibly harbored digestate derived OTUs, which were diluted out, rather than dying out, since their abundance declined with a rate largely as predicted by the dilution rate (**Fig. 3C** and **Figs. S12-13**). In autoclaved digestate, the absolute abundance of OTUs clustered in clade B and C was ~10^8^ and 10^9^ vial^−1^, respectively, while the abundance at the end of each enrichment was much lower, suggesting that their DNA is not destroyed by autoclaving, but that this relic DNA is degraded once the digestate is inoculated with live organisms. Thus, the high degree of clustering of samples by PCA (**Fig. 2C**) in the initial enrichments is probably not influenced by relic DNA as reported by others (Lennon et al 2018).

Clade D appeared to consist of soil specialist that sustained abundance in soil only, or alternatively, partly made up of relic DNA (DNA in the γ-sterilized soil) not metabolized during the enrichments in soil as mineral or humic substances may protect free DNA from rapid degradation (Nielsen et al 2007). However, some did appear to be true soil specialists due to their absence in the γ-sterilized soil (**Fig. 3A**). Our quantitative assessment confirmed that Clade D organisms grew in soil, while declining in digestate (**Fig. 3D**, and calculated *R* values **Fig. S14**). This clade harbored the soil specialist OTU74, circumscribing the isolated *Ochrobactrum* sp. **OB** (**Fig. 3G**), demonstrating the predicted characteristics of a soil specialist, reappearing at high abundance in soil enrichments.

Clade E showed an average increase in abundance throughout the enrichment in both enrichment lines but, interestingly, harbored organisms that were enriched to higher levels in the digestate derived line (D line) compared to the SD line (**Fig. 4B**), suggesting that they were suppressed by some organisms originating from the soil. Clade F appeared to contain organisms that were enriched in the SD line but remained at relatively low concentrations in the D line. OTUs of this clade were mostly soil derived organisms and their presence in the D line could be attributed to relic DNA (from the γ-soil) (Nielsen et al 2007) or an artifact of sequence OTU clustering.

**Figure 4:**
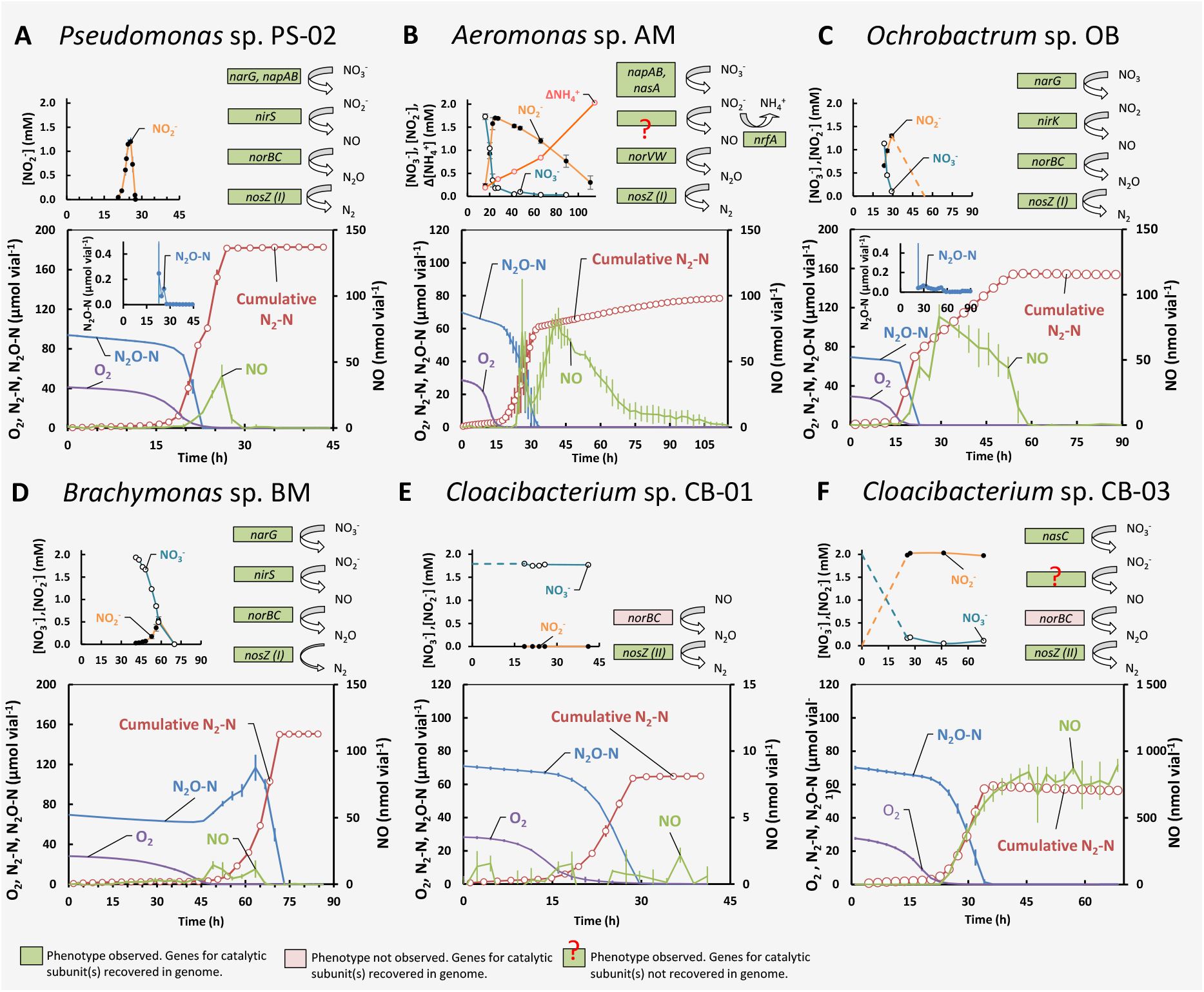
Denitrification genes and denitrification phenotypes of isolated organisms. Gas kinetics of O_2_, N_2_O-N, NO, and cumulative N_2_-N (adjusted for leakage and sampling) in denitrifying phenotype experiments in 120 mL closed vials with He atmosphere containing 50 mL liquid growth medium supplemented with 1 mL O_2_, 1 mL N_2_O and 2 mM NO_3_^−^. Liquid concentrations of NO_2_-, NO_3_^−^ and/or NH_4_^+^ (small panels, dashed lines = estimated by N-mass balance) and all genes coding for catalytic subunits of N-reductases recovered from the Prokka annotated genomes. **<u>Panel A</u>**: *Pseudomonas* sp. PS-01 (n = 2) grown in SS medium. PS-02 demonstrated strict control of gaseous denitrification intermediates throughout the incubation. **<u>Panel B</u>**: *Aeromonas sp*. AM (n = 3) grown in SS medium. AM demonstrated a DNRA+NOS phenotype, converting NO_3_^−^ to NO_2_^−^ and NH_4_^+^. Denitrification was ongoing throughout, but at a low and constant rate (0.4 μmol N_2_-N h^−1^ vial^−1^, **Fig. S22**). **<u>Panel C</u>**: *Ochrobactrum* sp. OB (n = 2) grown in SS medium. OB demonstrated strict control of gaseous intermediates throughout the incubation. **<u>Panel D</u>**: *Brachymonas sp*. BM (n = 2) grown in Sistrom’s succinate medium. BM demonstrated a full-fledged denitrifying phenotype where N_2_O was kept at high levels throughout the incubation. **<u>Panel E</u>**: *Cloacibacterium* sp. CB-01 (n = 3) grown in NB medium. CB-01 had a truncated denitrifying phenotype respiring primarily N_2_O. **<u>Panel F</u>**: *Cloacibacterium* sp. CB-03 (n = 2) grown in NB medium. CB-03 had a truncated denitrifying phenotype converting N_2_O to N_2_, and nitrate to nitrite.

### Eco-physiological genome analysis of isolated organisms

Throughout the enrichment series, the N_2_O-respiring organisms were apparently growing under C-substrate limiting conditions most of the time (**Fig. 2A**), and tracing the OTUs circumscribing the isolated organisms throughout the enrichment cycles showed that many of these organisms grew to, and maintained, high abundance throughout the repeated transfers, i. e. growing in both materials (**Fig. 3EFG**). Acquisition of less accessible nutrients for growth and proliferation could therefore in part explain why the isolated organisms outperformed other species throughout the enrichments. To explore this metabolic versatility, we annotated carbohydrate-active enzymes (CAZymes) and peptidases/proteases in the isolate genomes using the dbCAN meta server (Zhang et al 2018) and the MEROPS database (Rawlings et al 2010), respectively, and further analyzed the predicted proteins for putative signal peptides using SignalP5.0 (Raut et al 2021).

A range of genes coding for CAZymes were identified in all isolates (**Tab. S9**), several of which are known to target complex carbohydrates also contained putative signal peptides, indicating that these proteins are transported to the cell exterior. Isolates **CB-01** and **CB-03** seemed to have CAZymes focused on the breakdown of plant materials, coding enzymes involved in degradation of cellulose, cellulose derivatives and starch (**Tabs. S9-10**). This was supported by the presence of the carbohydrate binding module (CBM) 48, that binds to various linear and cyclic α-glucans derived from starch and glycogen (Koay et al 2010, Chaen et al 2012). **AM** had a large repertoire of genes encoding CAZymes with multiple CMBs associated with binding to cellulose (CBM5, Kezuka et al 2006), starch/glycogen (CBM48), peptidoglycans and chitin (CBM50, Onaga and Taira 2008) (**Tabs. S9-10**). All isolates, except **BM**, had genes encoding cellulases, and genes encoding glycogen synthase (EC: 2.4.1.21 and 2.4.1.11) were recovered in **PS-02**, **AM**, **CB-01**, **CB-03** and **OB**. **AM** and **CB-01/CB-03** had genes encoding glycogen operon protein glgX homolog (EC: 3.2.1.-) and glycogen phosphorylase (EC: 2.4.1.1), both catalyzing breakdown of glycogen. This may be associated with a fitness advantage during the dual culture enrichment as glycogen metabolism has been shown to improve *E. coli* fitness when experiencing changing environments (Sekar et al 2020). Contrastingly to the other isolates, **BM** did not appear to be geared towards extracellular degradation of complex carbohydrates, nor contained genes involved in glycogen metabolism (**Tabs. S9-10**) and might be dependent on harvesting easily available carbohydrates in the sterilized growth media. Interestingly, the ability of the isolated strains to grow in sterilized digestate was strongly related to the number of genes coding for proteases and CAZymes (**Fig. S32H**)

All isolates encoded peptidases containing putative signal sequences, but the relative proportion of these varied between the isolates; with **CB-03** having the largest proportion of predicted peptidases containing putative signal sequences, followed by **AM** and **CB-01** (**Tab. S11**). Extracellular secreted peptidases may also reflect environmental adaptations; the isolates contained peptidases active at a more neutral pH range and known low-pH active peptidases (Nguyen et al 2019) were not recovered. This falls in line with the inherent pH of the environments from which the isolates were obtained – neutral/alkaline digestate and weak acidic soil. Interestingly, the genomes **CB-01** and **CB-03** had several characteristics similar to predatory bacteria, such as overrepresentation of genes for peptidases, genes for the complete mevalonate pathway for isoprenoid production, histidine kinase (EC: 2.7.13.3), serine protease (EC: 3.4.21.107), FMN NADH reductase (EC: 1.7.1.17, only recovered in CB-01) and Dipate enol-lactone hydrolase (EC: 3.1.1.24) (Pasternak et al 2013). In contrast, **AB**, **BM**, **OB** and **PS-02** encoded the complete MEP/DOXP pathway, common for non-predatory bacteria (Pasternak et al 2013). We speculate that these genomic features may have contributed to **CB-01** and **CB-03** achieving dominance throughout the enrichment, but, to the best of our knowledge predatory traits of *Cloacibacterium* sp. has not been reported, and further experimentation would be required to confirm this.

Species within the genus *Ochrobactrum* can fix nitrogen symbiotically in root-nodules (Trujillo et al 2005, Zurdo-Pineiro et al 2007, Imran et al 2014). Some genes coding for proteins involved nodule formation where recovered in *Ochrobacter* sp. **OB** genome (**Tab. S12**), but not for the catalytic subunits of nitrogenase.

### Characterizing the isolates denitrifying regulatory phenotypes (DRP) and genotype

All isolates did encode the gene for Nos, *nosZ* (clade I or II, Hein and Simon 2019), as well as several other denitrification genes (**Fig. 4**). Although organisms with a full-fledged denitrification pathway can both produce and reduce N_2_O, they may be strong sinks for N_2_O in the environment, depending on their denitrification regulatory phenotype (**DRP**, Bergaust et al 2011), which is shaped by the regulatory network controlling their stepwise reactions of denitrification, both at the transcriptional (Spiro 2012) and metabolic (Mania et al 2020) level.

To characterize the DRP of our isolated strains, they were raised under strictly oxic conditions to secure absence of any denitrification proteins, and transferred to gas-tight vials with liquid medium containing 2 mM NO_3_^−^, and with He, O_2_ and N_2_O in the headspace. As these stirred cultures were allowed to deplete the oxygen and switch to anaerobic respiration, they were monitored for O_2_, NO, N_2_O and N_2_ in the headspace and NO_3_^−^, NO_2_^−^ and NH_4_^+^ in the liquid. Measured gases in incubations supplemented with 1 ml O_2_, 1 mL N_2_O and 2 mM NO_3_^−^, alongside with measured liquid concentrations of NO_2_^−^ NO_3_^−^ and NH_4_^+^ and genes coding for catalytic subunits, are shown for each isolate in **Fig. 4**.

The genomes of *Pseudomonas* sp. **PS-02**, *Ochrobactrum* sp. **OB** and *Brachymonas* sp. **BM**, predicted a full-fledged dentification pathway, i.e. reduction of NO_3_^−^ to dinitrogen gas, which was verified (**Fig. 4ACD**). However, theregulatory phenotypes were profoundly different: **PS-02** reduced available NO_3_^−^ and N_2_O concomitantly, before initiating NO_2_^−^-reduction (**Fig. 4A**). Nos activity was higher relative to the other N-reductases at the oxic/anoxic transition as there was only miniscule, transient accumulation of N_2_O during denitrification (NO_2_→NO→N_2_O→N_2_), and the preferential reduction of N_2_O was maintained if cultured with NO_2_^−^, with or without N_2_O in the headspace (**Figs. S17** – **S20**). The phenotype of **OB** (**Fig. 4C**) was very similar to that of **PS-02** and was maintained under a variety of conditions (**Figs. S21** – **S24**). **BM**, however, reduced most of the available NO_3_^−^ to N_2_O at first (**Fig 4D**), and this trait was retained if cultured with NO_2_^−^, with or without N_2_O in the headspace (**Figs. S25** – **S27**). This suggested that while **BM** would be a source of N_2_O in the environment, **PS-02** and **OB** would be strong sinks.

DNRA organisms with *nosZ* could be attractive inoculants since they reduce NO_3_^−^ to NH_4_^+^ rather than to N_2_, thus retaining plant-available N in the soil (Rütting et al 2011), and at the same time scavenging N_2_O produced by other organisms. The AM isolate simultaneously reduced the available NO_3_^−^ to NO_2_^−^ and N_2_O to N_2_ after O_2_-depletion (**Fig. 4B**, analyzed in more detail in **Fig. S28**), and subsequently reduced NO_2_^−^ to NH_4_^+^ and trace amounts of N_2_. This indicated dissimilatory nitrate reduction to ammonium (DNRA), which was corroborated by the presence of *nrfA* in the genome, coding for a key enzyme of DNRA (Cytochrome c552 nitrite reductase, EC: 1.7.2.2) (Einsle, 2011). It also carried a *nasD* gene that showed high sequence similarity (protein blast) with NirB (NADH-dependent nitrite reductase) of a related *Aeromonas media* strain. Genes for the nitrite reductases NirS/K were not identified, and the source for the produced NO remains unresolved. The **AM** genome also apparently lacked genes for the nitrate reductase NarGHI, while genes coding for periplasmic nitrate reductase Nap (*napAB*) and N_2_O reductase Nos (*nosZ*, clade I) were present. It also encoded the gene *nasA*, which showed high sequence similarity (protein blast) to a nitrate reductase of a related *Aeromonas media* strain. NasA is a constituent of the nitrate assimilatory system (Nas) and functions as a nitrate reductase in nitrate assimilation in a wide range of bacteria (Jiang et al 2015). The phenotypic analysis (**Fig. 4B**) showed that NO_3_^−^ and N_2_O were clearly reduced at the same time in incubations with the **AM** isolate. This contrasts earlier findings that Nos outcompetes Nap (for electrons) in denitrifying bacteria (Mania et al 2020), and it could be speculated that the gene annotated as NasA in **AM** may be responsible for the reduction of NO_3_^−^ to NO_2_^−^ that took place concomitantly with N_2_O reduction. However, none of the other genes generally found in the *nas* operon of bacteria, such as *nasFEC* and *B*, were detected in the genome analysis.

The genotypes of *Cloacibacterium* sp. **CB-01** and **CB-03** predicted a truncated denitrification pathway (NO→N_2_O→N_2_), and one (CB-03) was also equipped with genes for assimilatory NO_3_^−^ reductase (*NasC*, EC: 1.7.99.4) and a nitrite/nitrate transporter (*narK*). This was all verified by experiments showing stoichiometric conversion of N_2_O to N_2_ and reduction of NO_3_^−^ to NO_2_^−^ by **CB-03** (**Fig. 4EF**) and corroborated by experiments under a variety of conditions (**Figs. S29-S31**). The early onset of NO_3_^−^ reduction, before depletion of oxygen, suggesting that NasC was active under oxic conditions in this isolate, which was also reported for *Paracoccus denitrificans* (Pinchbeck et al 2019). Of the two isolates, **CB-01** makes for a particularly promising N_2_O-reducing soil inoculant. Both **CB-01** and **CB-03** were circumscribed by the dominating OTU1 of Clade A (**Fig. 3E**) which dominated in both D and SD enrichment lines.

### Performance of isolated organisms as sinks for N_2_O in soil

To produce inocula for testing isolates’ capacity as N_2_O sinks in soil, they were grown aerobically to high cell densities in autoclaved digestate (**Fig. S32**). The estimated the cell density at the end of the 45 h incubation was ranged from 0.5 to 1.4 mg dry-weight mL^−1^ (~3-7 · 10^10^ cells mL^−1^) for the different isolates, the lowest value recorded for *Brachymonas* sp. **BM** (0.5) while Aeromonas sp. **AM** reached highest (1.4). Interestingly, the capacity of the isolates to grow was strongly correlated with the number of genes coding for CAZymes and proteases in their genomes (**Fig. S32H**).

To assess the N_2_O sink capacity of these aerobically grown organisms, they were inoculated to soil in vials with He atmosphere (with traces of O_2_), which were monitored for O_2_, NO, N_2_O and N_2_. Since the effect of such inoculation confounds the effect of the isolates and the effect of the available carbon in the autoclaved digestate, we included a set of control treatments. Thus, the experiment included 5 different pre-amendments: 1) Autoclaved digestate enriched with isolate by aerobic growth, 2) live digestate sampled directly from the anaerobic digester, 3) live digestate heat-treated to 70 °C for two hours (70 °C digestate) to reduce the activity of native N_2_O producers (Jonassen et al 2021) 4) digestate in which CB-01 was grown, subsequently heated to 70 °C for two hours (70 °C O_2_ dig) to give comparable amounts of carbon added to the soil, and 5) soil without any amendments. The *I_N2O_*-emissions ratio, which is the area under the N_2_O-curve divided by the area under the N_2_O+N_2_-curve (Liu et al 2014; Russenes et al 2016) expressed as a percentage, was used as a proxy to assess the treatment effects in the amended soils (**Fig. 5**).

**Figure 5:**
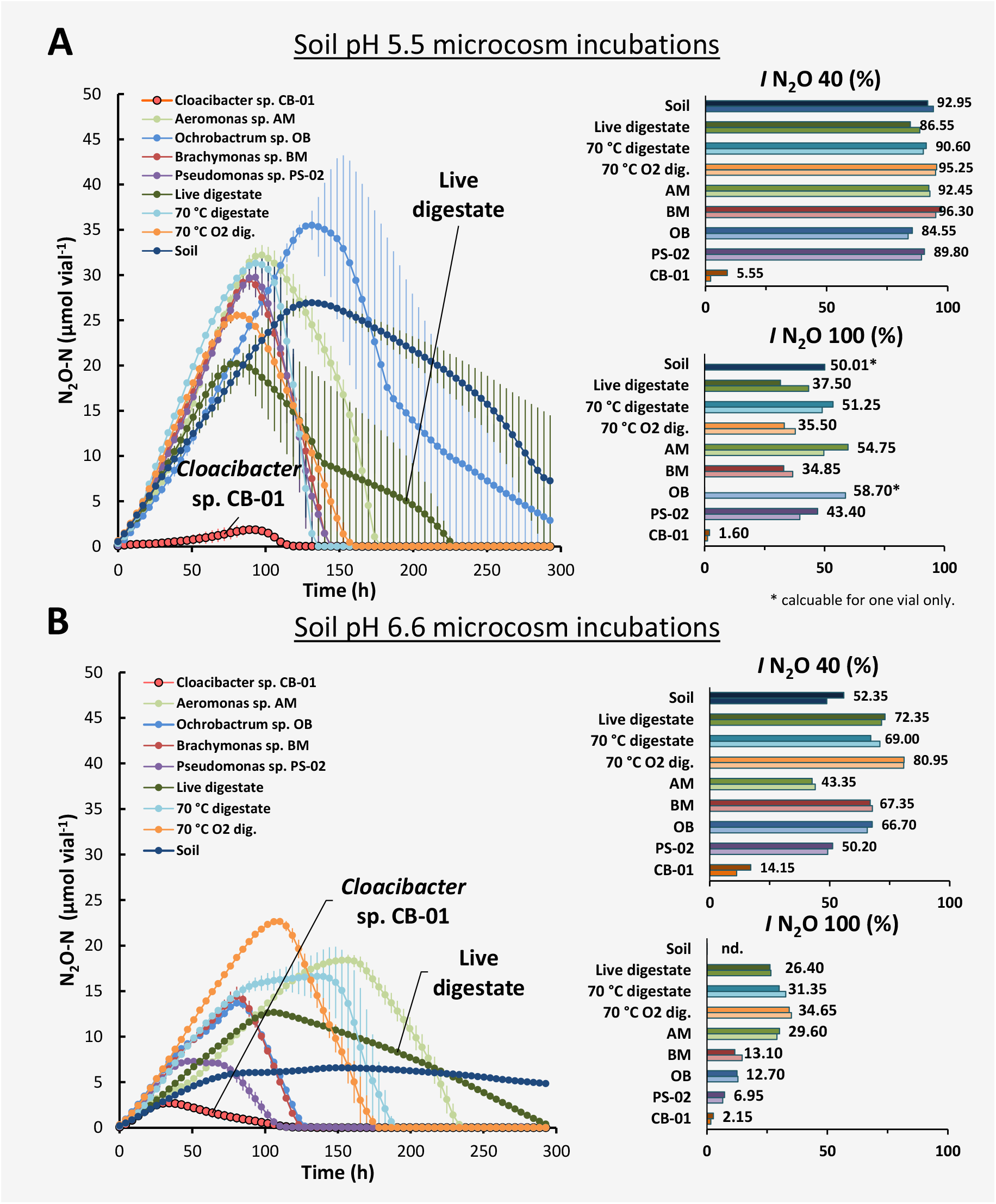
Gas kinetics during incubation of soils (**Panel A**: soil pH=5.5, **Panel B**: soil pH=6.6) amended with various pretreated digestates (0.06 mL g^−1^ soil): “Live digestate” = digestate directly from the anaerobic digester, “70 °C digestate” = live digestate heat treated to 70 °C for two hours, “70 °C O_2_ dig.” = autoclaved digestate used to grow strain CB aerobically, and then heat treated to 70 °C for two hours. **AM**, **BM**, **OB**, **PS-02** and **CB-01**: autoclaved digestate in which strains **AM, BM, OB, PS-02 and CB-01** had been grown aerobically.) to cell densities of 1.39, 0.51, 0.79, 0.81 and 0.72 mg cell dry-weight mL^−1^, respectively (**Fig. S32**). The main graphs (panels A&B) show N_2_O kinetics for each treatment (n = 2). The two bar graphs to the right show the N_2_O indexes (***I_N2O_**,* single vial values), which is the area under the N_2_O-curve divided by the area under the N_2_O+N_2_-curve expressed as a percentage. Two ***I_N2O_*** values are shown: one for the timespan until 40% of the NO_3_^−^ - N is recovered as N_2_+N_2_O+NO-N (**I_N2O 40%_**), and one for 100% recovery (**I_N2O 100%_**). More details (including N_2_ and NO kinetics) are shown in **Figs. S33 and S34**.

As expected, ***I_N2O_*** values were generally higher in the pH 5.5 soil than in the pH 6.6 soil (**Fig. 5B**), and the isolates BM, OB, PS-02 lowered ***I_N2O_*** only in the soils with pH 6.6. CB-01, however, resulted in extremely low ***I_N2O_*** values in both soils, clearly outperforming any of the control treatments. We tested if the ability of **CB-01** to act as a strong N_2_O sink in the pH 5.5 soil could be due to acid tolerance, by growing the organisms in stirred (600 rpm) liquid medium with pH ranging from 5.5 to 7, and found no evidence for acid tolerance, neither for growth nor for the synthesis of functional N_2_O-reductase (**Fig. S31**). An alternative explanation of the acid-tolerant N_2_O-sink effect of **CB-01** could be that the cells were embedded in flocks/biofilms in the digestate, protected against low soil-pH by the buffer-capacity of the matrix. Strains of *Cloacibacterium* are known to secrete extracellular polymeric substances (Nohua et al 2015) and found in high abundance in biofilms of natural (Pang et al 2016), which lends support towards the hypothesis of matrix mediated shielding effects. This points towards the advantages of biofilm formation or other attachment strategies in generating favorable micro niches and so gaining advantage over competitors in a low pH environment.

Whilst our eco-physiological genome analysis revealed that several isolates had the genetic potential to utilize complex carbon sources and encoded several traits that might secure survival in a competitive situation, agricultural inoculants are most definitely invaders of the soil microbial community, and any longer-term establishment is dependent on the resistance by the residential community against alien species. The likelihood of a successful invasion is related to the resident community richness, referred to as the diversity-invasion effect (Mallon et al 2018), and reflects the key challenges of an invading organism; growth and establishment by utilizing resources not utilized by the resident community, or forcefully “overtake” a resident niche through competition or antagonism.

To assess the ability of our isolates to persist in soil and to retain their N_2_O reduction capacity, a second experiment was set up with identical treatments to those in **Fig. 5**, but storing the amended soils for 1 month before testing the denitrification kinetics. A fertilization event was simulated by the addition of 50 μmol NO_3_^−^, 1 mg ground plant material g^−1^ soil, and 0.5 mL O_2_ before sealing vials and monitoring denitrification kinetics throughout depletion of oxygen and the transition to anoxia. In this experiment the effect of the inoculated isolates on N_2_O - emissions was evaluated based on maximum N_2_O accumulation (no treatment reduced all available N-oxides, making it impossible to calculate *I_N2O_* emission indexes) (**Figs. S35-36**). Whilst none of the inoculants significantly differed from the controls in pH 5.5 soil, PS-02 outperformed the other inoculants at pH 6.6. In fact, the soil treated with **PS-02** performed better after 30 days soil storage (maximum N_2_O for PS-02 was ~1/10 of other treatments, **Fig. S36**) than immediately after amendment in the first soil experiment (**Fig. 5**). Likewise, maximum N_2_O for **CB-01** treatment in pH 6.6 soil was approximately 2/3 of other amendments, but the difference was not statistically significant (p > 0.05).

A dose-response experiment with the isolates **CB-01**, **PS-02**, and **OB** was conducted to determine the minimum dose needed to obtain substantial reduction of N_2_O production in soil. The isolates were grown aerobically in autoclaved digestate (pH adjusted to 7.5) as explained above, the cell density achieved was assessed by the cumulated oxygen consumption (explained in detail in **Fig. S37**), and the cell density was adjusted to 0.3 mg cell dry-weight mL^−1^ for all three strains, by dilution with autoclaved digestate. These enriched digestates were then used in an amendment experiment identical to that presented in **Fig. 5**, but with three different doses of enriched digestates (0.6, 0.3 or 0.15 mL; triplicates for each level), which is equivalent to an inoculation intensity of 18, 9 and 4.5 μg cell dry-weight g^−1^ soil, or 9, 4.5 and 2.3·10^7^ cells g^−1^ soil, assuming the same dry-weight per cell as *Paracoccus denitrificans* (2·10^−13^ g dw cell^−1^). The highest inoculation intensity in this experiment is approximately 50% of that used in the previous experiments (**Fig. 5**). The experiment included controls, amended with equivalent doses of sterile pre-aerated autoclaved digestate.

The results (summarized in **Fig. S38** and **Tab. S13**) showed a strong and dose-dependent effect of *Cloacibacterium* sp. **CB-01** on the N_2_O accumulation, exemplified with the peak N_2_O concentration (Max N_2_O), which was reduced by 96, 64 and 20% (compared to the control without bacteria) by the inoculation levels 0.6, 0.3 and 0.15 mL digestate vial^−1^, respectively (p<0.05 for all contrasts). *Pseudomonas* sp. **PS-02** and *Ochrobacter* sp. **OB** had weaker effects on Max N_2_O, but statistically significant (p<0.05) at all inoculation levels. The ***I_N2O_*** showed the same patterns, but several contrasts (bacteria versus control) lacked statistical significance for the lowest inoculation level.

Our inoculation levels were 2.7, 4.5 and 9·10^7^ cells g^−1^ soil, which is within the upper range of inoculation levels used by Domeignoz-Horta et al (2016), who inoculated soils with 10^6^ and 10^8^ *Dyadobacter fermentans* – cells g^−1^ soil. *Dyadobacter fermentans* carry *nosZ* Clade II, but no other denitrification genes, which makes it comparable to our *Cloacibacterium* sp. CB-01, and a comparison of the performance of the two strains is interesting: inoculation with 10^8^ *D. fermentans* - cells g^−1^ resulted in a reduction in the N_2_O/(N_2_O+N_2_) product ratio which is similar to what was achieved by the two highest inoculation levels with *Cloacibacterium*, i.e. 0.45-0.9·10^8^ cells g^−1^. Thus, the two strains appear to have similar capacities for acting as sinks for N_2_O in soil. However, a closer inspection of the data reveals that *Dyadobacter* did not affect the N_2_O-emission in soils with pH below 6.6, while *Cloacibacterium* performed well in our acid soil (pH 5.5, **Fig 5A**). This could indicate that *Cloacibacterium* sp. **CB-01** has a more robust N_2_O sink capacity in low-pH soils. As suggested previously, this is probably not due to an inherent acid tolerance, but rather a combined effect of the organisms’ tendency to aggregate and form biofilms, and the relatively high pH of the digestate (7.6). The matrix in which cells are embedded prior to inoculation to soils is probably a crucial issue.

### Concluding remarks

The hierarchical clustering of 16S rRNA-based OTUs demonstrated that the dual enrichment effectively selected “generalist organisms” capable of growth by N_2_O-respiration both sterilized digestate and soil, already after 3-4 transitions, as predicted by the model. (**Figs. S1** to **S5**).

Among the isolates, *Cloacibacterium* sp. **CB-01** stand out as particularly interesting because it grew well both in soil and digestates and was unable to denitrify *sensus stricto* (lacking the genes for dissimilatory NO_3_^−^ and NO_2_^−^ reduction). In addition, it proved a strong N_2_O sink even in the acid soil (pH 5.5), where the other isolates appeared unable to synthesize functional N_2_O reductase, as is the case for most organisms (Liu et al 2014, Lim et al 2021). Testing the pH-response of CB-01 in pure culture showed no particular tolerance to low pH (**Fig. S30**). We speculate that its ability to reduce N_2_O in low pH soil is due to the ability of this organism to localize in alkaline microinches supplied by the digestate material, possibly through the production of a biofilm, a trait known to be common to members of this genus (Tiirola et al 2009, Revetta et al 2013, Biswas et al 2014, Pang et al 2016). The ability to retain activity in low pH soils is very desirable for agriculture due to the issue of soil acidification, driven by N-input and subsequent base cation depletion in agricultural soils (Tian et al 2015), and the uncertainty in net GHG-emission reduction of the few viable treatment options such as liming (Wang et al 2018, Hénault et al 2019) to mitigate N_2_O derived from denitrification in these soils, but at a possible expense of increased emissions of carbonate-CO_2_ (Wang et al 2021). The second isolate to show promise is **PS-02**. While **PS-02** can act as both a source and sink for N_2_O, it showed the benefit of eliciting an N_2_O-emissions reduction for an extended period after soil amendment. An interesting possibility and a future perspective is the option of combining **PS-02** and **CB-01**, to secure effective elimination immediately after fertilization (**CB-01**) as well as a more long-lasting effect (**PS-02**).

Not all the members of the generalist Clade A OTUs were obtained as pure cultures and extended isolation efforts may uncover yet more organisms with good qualities for an amended digestate material. Further, this enrichment technique could easily be expanded to new soils and new digestates to develop amendments suited to specific local materials and conditions. Future research investigating the performance of digestate amendments derived from these isolates would be valuable to accurately quantify the N_2_O-emissions reduction effects under field conditions.

## Acknowledgements

This work was financially supported by the Norwegian Research Council (project number: 260868). We wish to thank Elisabeth G. Hiis for indispensable assistance with the automated gas incubation and measurement system.

