## Supplementary materials for "A novel dual enrichment strategy provides soil- and digestate-competent N_2_O-respiring bacteria for mitigating climate forcing in agriculture"

### 1 Dual enrichment, conceptual model

The idea was to enrich N_2_O-reducing organisms that are able to grow both in soil and digestate by dual enrichment culturing, i.e. sequential anaerobic batch incubations (with N_2_O) in the two substrates, starting with unsterilized digestate, with or without unsterilized soil. At the end of the batch incubation, a fraction is used to inoculate the next batch where the substratum is gamma-sterilized soil. At the end of this incubation, a fraction is then used to inoculate the next batch where the substratum is autoclaved digestate, and so on. Each batch incubation should be long enough to secure depletion of easily available carbon sources, to secure competition between the populations.

To visualize the selection of organisms depending on experimental conditions (length of incubation, fraction of one batch transferred to the next) and the properties of the organisms, we made a simple mathematical model with three conceptual types of organisms:

- ***D*** = digestate specialist: fast growth in digestate, gradual death in the soil
- ***S*** = soil specialist: fast growth in soil, gradual death in the digestate
- ***G*** = generalist: growth on both substrata, but slower than the specialists (in their preferred substrate)

All three were assumed to compete for the same pool of carbon substrates, and the competition was implemented by assuming logistic growth for each depending on the total increase in cell density (i.e. the sum of all three populations), and first-order death rates. The differential equations for the growth and death of the three population in a single batch are:

$\frac{dN_{D}}{dt}=N_{D}*r_{D}\left( 1-\frac{N_{t}}{K} \right)-N_{D}*d_{D}$	(1)

$\frac{dN_{S}}{dt}=N_{S}*r_{S}\left( 1-\frac{N_{t}}{K} \right)-N_{S}*d_{S}$	(2)

$\frac{dN_{G}}{dt}=N_{G}*r_{G}\left( 1-\frac{N_{t}}{K} \right)-N_{G}*d_{G}$	(3)

Using ***D*** as an example to explain the model: ***N_D_*** is the population size of D (cells mL^-1^), ***r_D_*** (h^-1^) is its maximum growth rate (high for digestate, low/zero for soil), ***Nt*** is the summed growth of all three populations, ***K*** is the substratum’s carrying capacity (i.e. the maximum cell number that can be produced in the substratum), *d_D_* (h^-1^) is the first order death rate. The growth and death rates are substrate-specific: for cultivation in digestate, D has high ***r_D_*** and low (or zero) ***d_D_*** , while the opposite is the case for cultivation in soil: ***r_D_*** is low (or zero), ***d_D_*** is high.

Thus, in digestate, D will increase at a high rate as long as N_t_<<K, the rate decline as N_t_ converge to K, reach zero when *N_t_*=*K*(1-d_D_/r_D_) and decline if *N_t_*>*K*(1-d_D_/r_D_), provided that ***d_D_***> 0. In soil, the abundance is constant if ***r_D_*** for soil is set to zero, and D will die out (first order) by a rate given by ***d_D_***. The two other populations (equations 2 and 3) were simulated the same way, with the same ***K***-value as for D, but with individual substrate-specific growth- and death rates. The model calculates ***N_t_*** by summing up the net increase of the three populations, while any decline is not affecting ***N_t_***.

The model was implemented in excel, simulating the growth of each population by forward Euler. A simulation example is shown in **Fig. S1**, illustrating features of D, S and G type organisms.

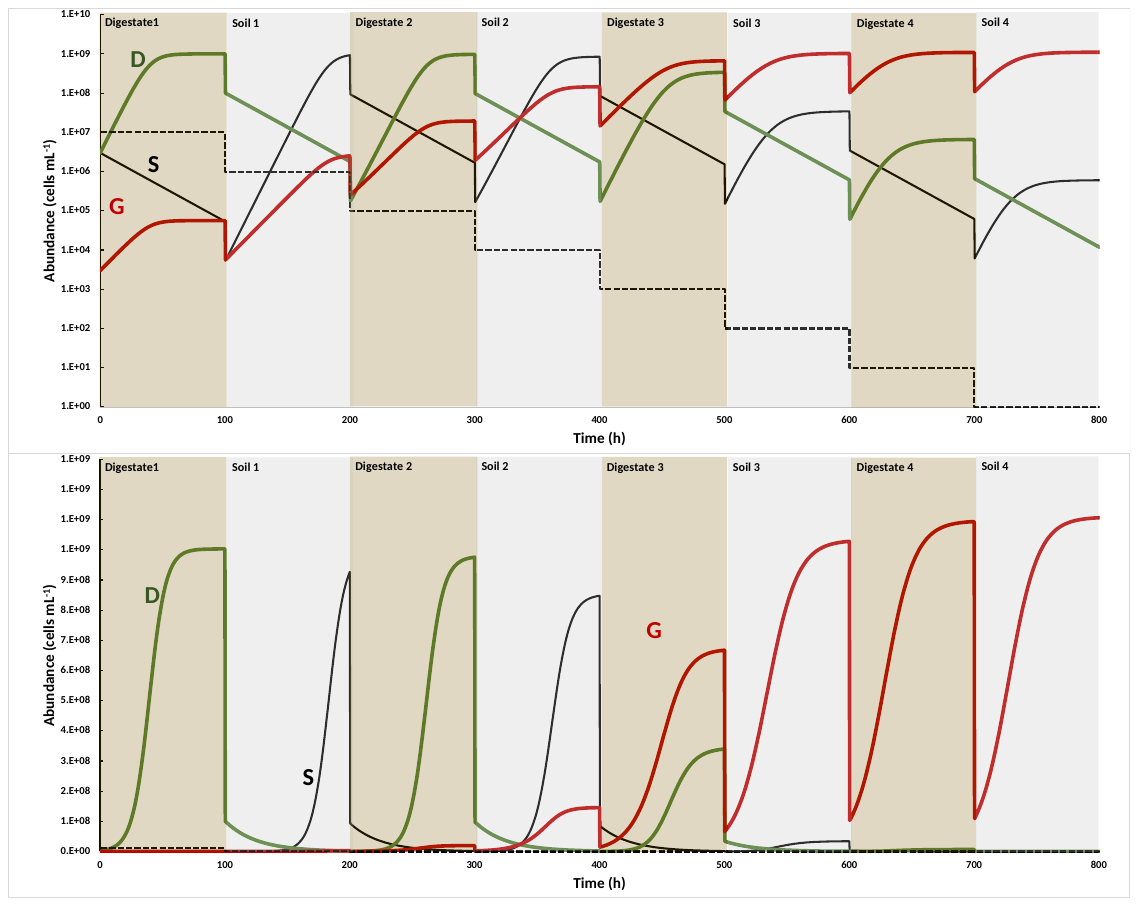

**Figure S1: Simulation of the competition between three populations through a series of enrichment cultures.** The three populations (S, D and G, see text for explanation) were simulated with the parameter values shown in **Table S1**, 100 hours incubation time for each batch and transfer of 10% of the culture volume to the next batch. Top panel shows abundance on log scale, bottom panel on a linear scale. D and S are sustained at stable levels (but fluctuating with substrates) only until G-abundance reaches significant levels. Thereafter they decline and approach extinction (< 1 cells mL^-1^) if continuing the enrichment through 6-7 more batches (result not shown). The dashed line is the predicted dilution of a population which neither grows nor dies.

**Table S1:** Codes for substrate-specific rate constants (equation 1-3), and values used for simulation shown in Fig. S1, including the initial cell abundance for the soil specialist (S), the digestate specialist (D), and the generalist (G). *K*=10^9^ cells mL^-1^ both for soil and digestate.

| **Organism** | **Initial** | **Rate constants (h^-1^) in:** | | | | |
| --- | --- | --- | --- | --- | --- | --- |
|  | **abundance** | **Digestate** | | **Soil** | | |
|  | **(cells mL^-1^)** | ***r*** | ***d*** | ***r*** | ***d*** | |
| **D** | 3.E+06 | ***r_D_dig_*** = 0.15 | ***d_D_dig_*** = 0 | ***r_D_soil_*** = 0 | | ***d_D_soil_*** = 0.04 |
| **S** | 3.E+06 | ***r_S_dig_*** = 0 | ***d_S_dig_*** = 0.04 | ***r_S_soil_*** = 0.15 | | ***d_S_soil_*** = 0 |
| **G** | 3.E+03 | ***r_G_dig_*** = 0.075 | ***d_G_dig_*** = 0 | ***r_G_soil_*** = 0.075 | | ***d_G_soil_*** = 0 |

In the following, the sensitivity of the model to parameter values was tested by changing one parameter at a time, using the parameter values and initial population densities in **Tab. S1** as default.

The fraction transferred from one batch to the next had a significant effect on the selective pressure as shown in **Fig. S2**.

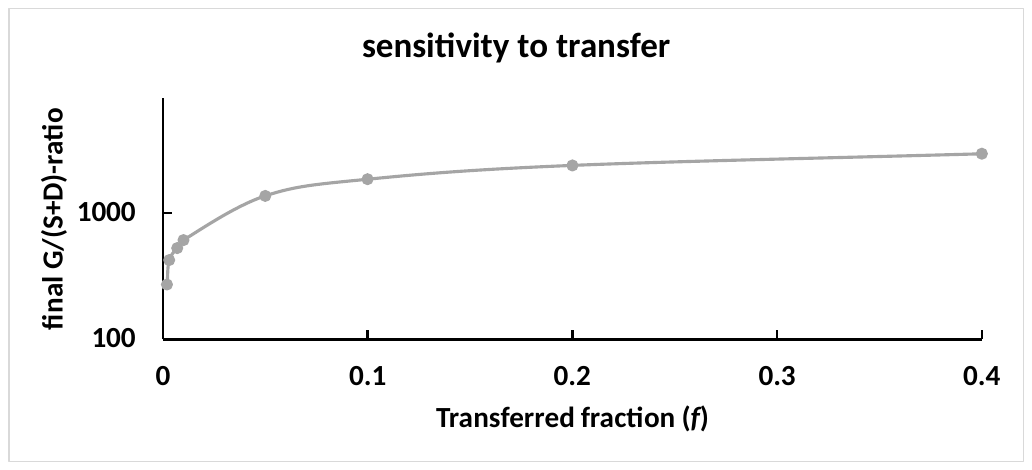

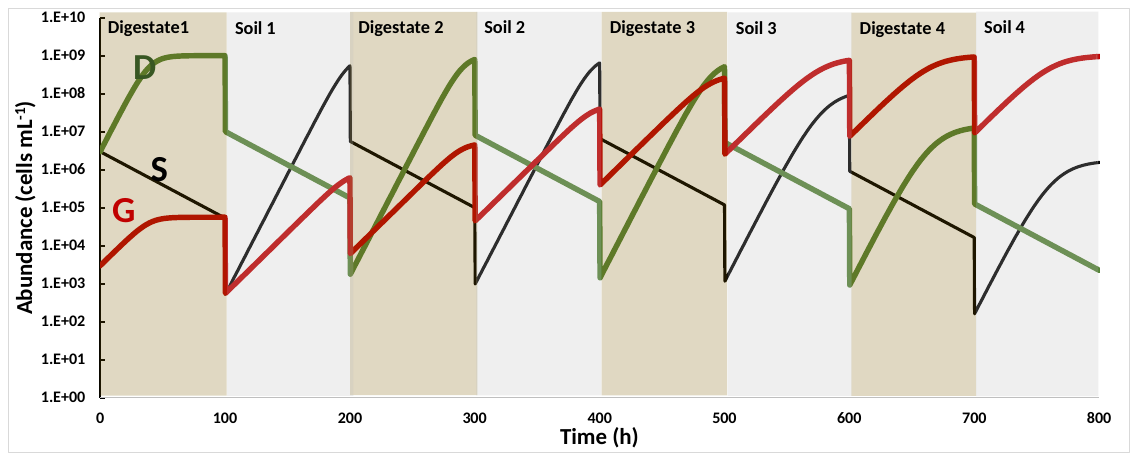

**Figure S2: Selective pressure depends on the fraction (*f*) transferred from one batch to the next.** The top panel shows the final G/(S+D) abundance ratio (i.e. in Soil 4, **Fig. S1**), for different values of ***f***, all other parameters were identical to that in **Fig. S1.** The G/(S+D) ratio is a measure of the selective pressure favoring G over S and D, and this shows drastic decline as ***f*** decrease below 0.1 (i.e. 10% transfer). The reason for this is that at very low ***f***, **N_t_** remains << K throughout most of the time, hence growth is not limited by substrate availability, resulting in lower selection pressure. The phenomenon is illustrated in the bottom panel which shows a simulation for ***f***= 0.01. Selective pressure could be restored for ***f*** = 0.01 by increasing the time span for each batch cultivation (result not shown).

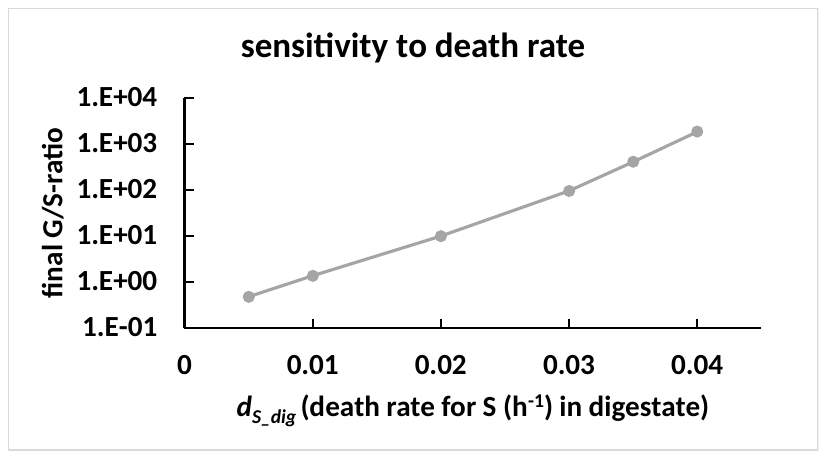

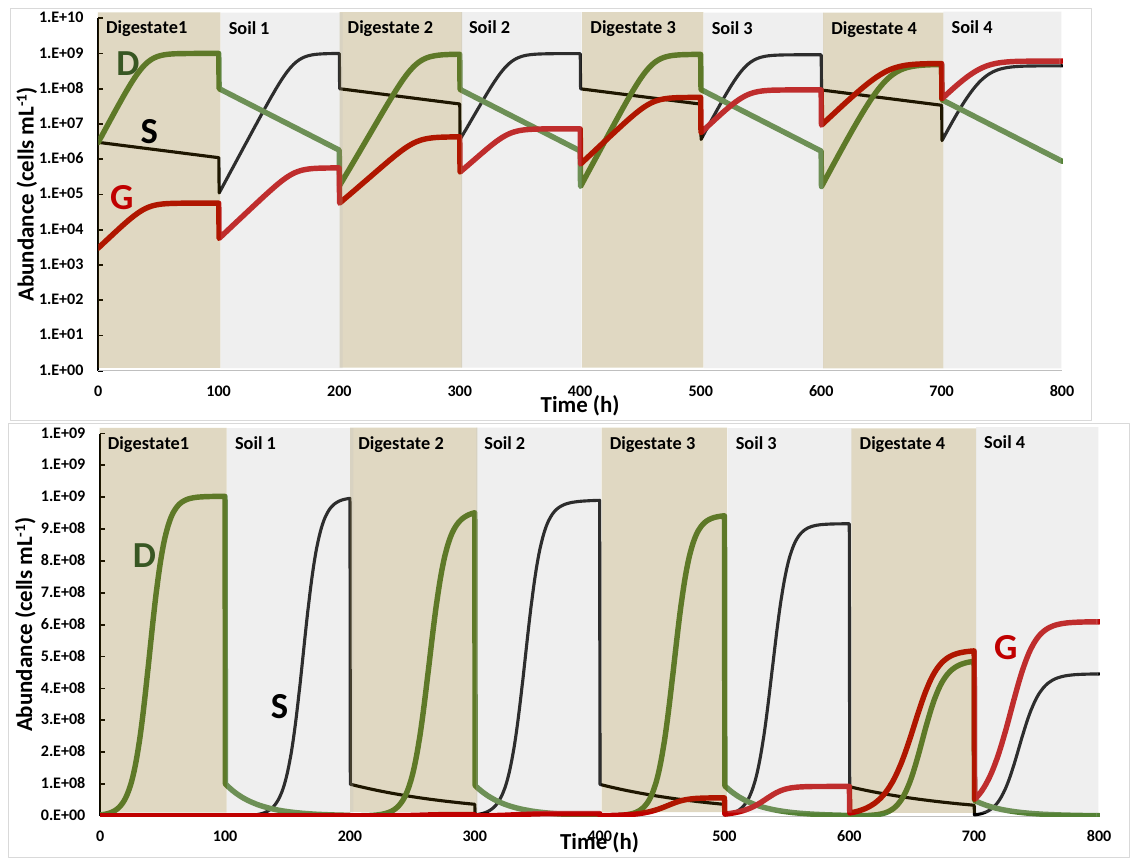

**Figure S3: Sensitivity to death rate of S in digestate (*d_S­_Dig_).*** To explore the sensitivity to death rate for S in digestate (***d_S­_Dig_***), simulations were run with different values (all other parameters as in **Tab. S1**). Top panel shows the final G/S ratio plotted against ***d_S­_Dig_*** (h^-1^). Bottom panel shows simulation for ***d_S_Dig_*** = 0.01 h^-1^ (Log scale in upper panel and linear scale in the lower). Although G will ultimately become dominant at any ***d_S_Dig_***>0, and ultimately exclude both S and D (NB: *d_D__soil* = 0.04 h^-1^ in these simulations), this would require a continuation of the dual enrichment (tested by simulation of a sequence of 24 batches, result not shown).

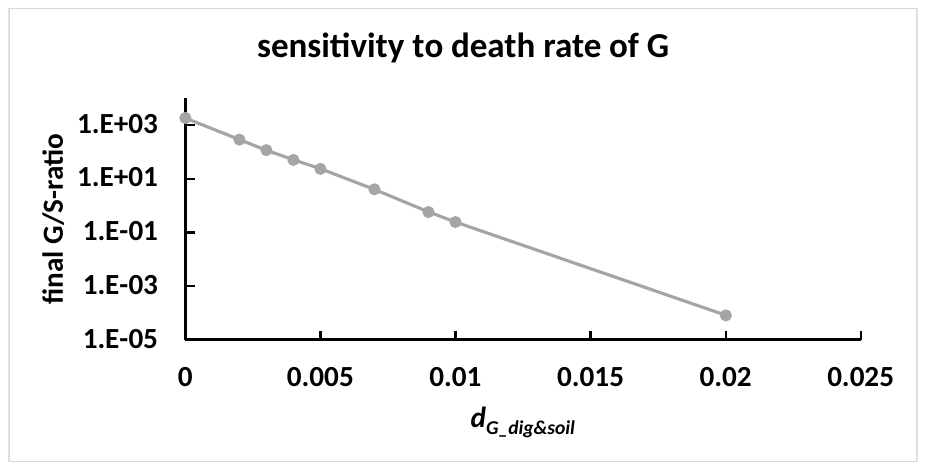

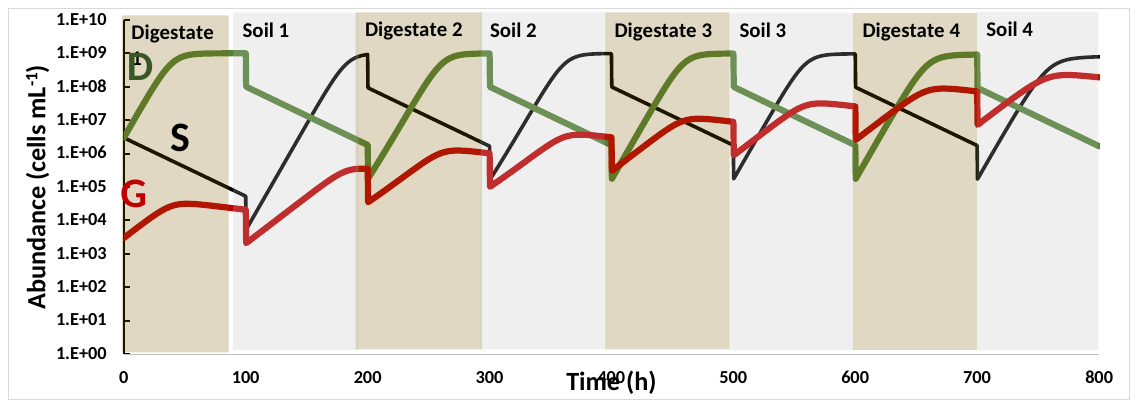

**Figure S4: Sensitivity to death rate of G**. The top panel explores the effect of cell death in G: Parameter values as in **Table S1,** except for the death rates of G in digestate (***d_G-Dig_***) and soil (***d_G-soil_***) which were both stepped up from 0 to 0.02 h^-1^ ( ***d_G-Dig_*** = ***d_G-soil_*** for each simulation), and the result is shown as the G/S abundance ratio after 800 h (i.e. at the end of the last enrichment in soil), plotted against ***d_G-Dig&soil_*_._** The result suggests selection against G when ***d_G-Dig_*** = ***d_G-soil_*** = 0.02 h^-1^ but this is not the case: by simulating a continuation of the enrichment through 20*8=160 enrichments for ***d_G-Dig_*** = ***d_G-soil_*** = 0.02 h^-1^, the G/S abundance ratio increased slowly but steadily, reaching 13 at the end (result not shown). This illustrates that although a competitive generalist can become dominant by dual enrichment culturing through 7-8 steps even at very low initial abundance (**Fig S1**), it would take very many batch cultivations for a less competitive organism to become dominant if it’s initial abundance is low. The bottom panel shows the simulated population dynamics for ***d_G-Dig_*** = ***d_G-soil_*** = 0.01 h^-1^: in this case, G almost reached dominance at the end of the first 8 batch cultivations. Extended simulation showed that G reached >10 times higher than S and G after 4 additional batches (result not shown).

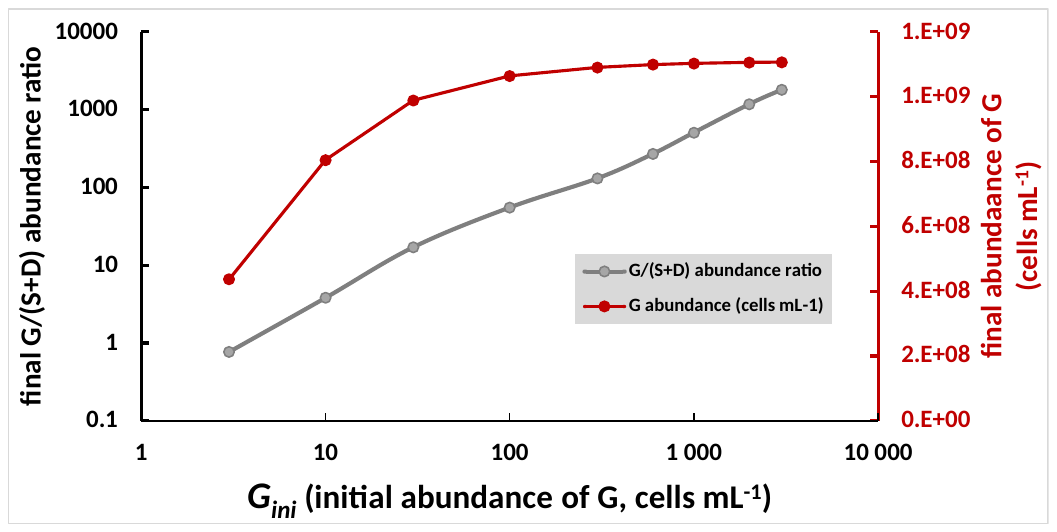

**Figure S5: Minimum initial abundance of a competitive G to become dominant.** To the explore the minimum initial abundance for competitive generalist (as modelled in **Fig. S1**) to become dominant through 8 cycles, simulations were run with different initial G-abundancies (***G_ini_***) , and the result is shown as the final G/(S+D) abundance ratio (log scale) and the final abundance of G after 8 batches, plotted against ***G_ini_*** (log scale). This illustrates that the final G abundance approach it’s maximum (K=10^9^ cells mL^-1^) at ***G_ini_*** around 100 cells mL^-1^, while the G/(S+D) ratio continued to increase with increasing ***G_ini_*** due to earlier onset of decline for D and S.

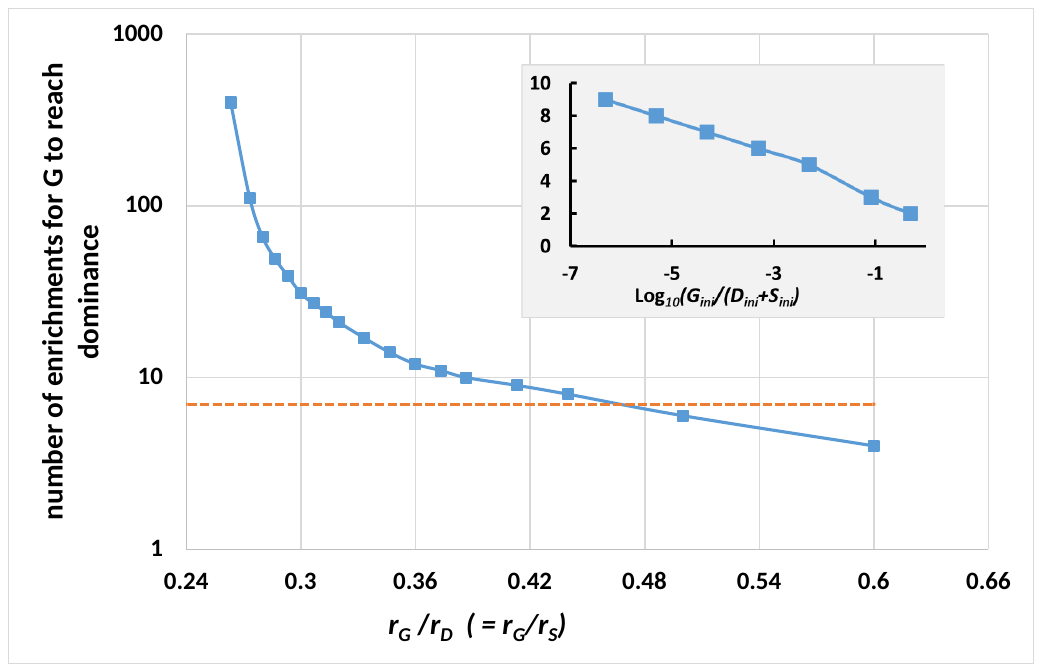

**Figure S6: Number of enrichments needed for weakly and strongly competitive generalist to become dominant.** To explore the minimum growth rate for G (r_G_) to become dominant within 7-8 sequential enrichment cultures, and to explore the number of enrichment cultures that would be needed to reach dominance for weakly competitive generalists, simulations were run with a series of ***r_G_*** values, but otherwise with the same parameters as in Table S1. The simulations were run until **G** reached dominance, arbitrarily defined as **G** > 10*(**D**+**S**). The panel shows the number of sequential enrichment cultures needed to reach G-dominance, plotted against *r_G_/r_S_* (which is equal to *r_G_/r_D_* because *r_D_*=*r_S_*=0.15 h^-1^ for all simulations). The dashed line mark 7 enrichment cultures. This shows the minimum ***r_G_*/*r_D_*** -ratio for **G** to be dominant in the 7^th^ enrichment is ~0.47, i.e. the minimum rG= 0.07 h^-1^ (***r_G_***=0.15*0.47= 0.07 h^-1^). For a generalist with rG<0.07 h^-1^, higher number of enrichment cultures are needed. NB. In the present simulations, the initial population ***G_ini_*** is 1000 times lower than ***S_ini_*** and ***D_ini_***. For higher ***G_ini_,*** lower number of enrichments are needed. The minimum ***r_G_/r_D_*** ratio for G to be competitive is 0.26 (***r_G_/r_D_* = *r_G_/r_S_*** < 0.26 **results** in washout of G). The exercise shows that 7-8 enrichment cultures would be enough to enrich a competitive generalist, even at lower initial numbers than that used in previous simulations. The latter is illustrated in the inserted panel, showing the necessary number of enrichments for G to become dominant for ***r_G_***= 0.075 h^-1^ (i.e. ***r_G_/r_D_***=***r_G_/r_S_***= 0.5), plotted against the log_10_ value of the relative initial abundance of G (***G_ini_/(S_ini_+D_ini_)***).

### 2 Supplementary materials and methods

**Materials for inoculum and growth substrates**

**Table S2: Digestate characteristics at the time of sampling for enrichment culturing and soil incubations**. The digestate was taken from an anaerobic digester of a municipal WWTP (same as that used by Jonassen et al 2021). Enrichment culturing and soil incubations were done with freshly sampled digestate (Sampling 1-4). Digestate characteristics were analyzed in the NS-EN ISO/IEC 17025 accredited laboratory belonging to the WWTP.

|  | **Digestate characteristics** | | | | | | |
| --- | --- | --- | --- | --- | --- | --- | --- |
|  | pH | % dry weight ^a^ | LOI ^b^  (% of DW) | TAK ^c^  (meq L^-1^) | VFA ^c^  (meq L^-1^) | VFA/TAK | NH_3+_NH_4_^+^  (mg-N L^-1^) |
| Sample 1^d^ | 7.61 | 4.12 | 55.93 | 186 | 16.7 | 0.090 | nd. |
| Sample 2^d^ | 7.73 | 3.97 | 55.45 | 199 | 17.3 | 0.087 | nd. |
| Sample 3^d^ | 7.84 | 3.87 | 55.46 | 201 | 16.3 | 0.081 | nd. |
| Sample 4^d^ | 7.60 | 3.73 | 54.87 | 189 | 15.2 | 0,080 | 1486 ± 7^e^  (1883 ± 3) |
| Average^f^ | **7.70** | **3.92** | **55.17** | **194** | **16.4** | **0.085** | **1824** |

^a^ Dry weight % expressed as percentage of wet weight (determined according to EN15934, given by WWTP). ^b^ Loss of ignition (volatile solids) as percentage of dry weight (determined according to EN15935, given by WWTP). ^c^ VFA = volatile fatty acids. TAK = total alkalinity (determined by titration described in EN12176:1998, given by WWTP). ^d^ Sample 1: sample used in enrichment experiment (live digestate inoculum, D_A-G.1_ and SD_A-G.1_, shown in **Fig. 2A** in main paper and Figs. **S8** and **S9**. Source of autoclaved digestate used as growth substrate in enrichment culturing. Sample 2 was used for aerobic growth of isolates, (**Fig. S32**). Sample 3 was used in soil incubations (live digestate) (**Fig. 5** in main paper and **S33-36**), Sample 4 was used in soil dose response experiment (**Figs. S38-38**). ^e^ Aeration of autoclaved digestate stripped off NH_4_^+^ (concentration in live digestate given in parenthesis). ^f^ Yearly average digestate characteristics given by the WWTP. Ammonium concentrations measured at the WWTP was measured as described by Greenberg et al (1980).

**ddPCR**

**Table S3**: PCR cycle settings for 16S ddPCR with primer pairs PRK341F/PRK806R.

| Time: | Temperature (°C): | Description: | |
| --- | --- | --- | --- |
| 5 min | 95 | Denaturation | |
| 30 s | 95 | Denaturation | 40 cycles |
| 30s | 55 | Annealing |  |
| 45 s | 72 | Extension |  |
| 5 min | 4 | Signal stabilization | |
| 5 min | 90 | Signal stabilization | |
| Indef. | 4 | Hold step | |

**Media composition**

Sistrom’s succinate medium (**SS**): contained (L^-1^) 3.48 g K_2_HPO_4_, 0.195 g NH_4_Cl, 4 g succinic acid, 0.10 g glutamic acid, 0.04 g aspartic acid, 0.5 g NaCl, 0.2 g nitrilotriacetic acid, 0.3 g MgSO_4_ · 7H_2_O, 0.015 g CaCl_2_ · 7H_2_O, 0.002 g FeSO_2_ · 7H_2_O, 0.1 mL trace element solution and 0.1 mL vitamin solution. The trace element solution contained (g L^-1^): 17.65 g EDTA (triplex 3), 109.5 g ZnSO_4_ · 7H_2_O, 50 g FeSO_4_ · 7H_2_O, 15.4 g MnSO_4_ · H_2_O, 3.92 g CuSO_4_ · 5H_2_O, 2.48 g Co(NO_3_)_2_ · 6H_2_O and 1.14 g H_3_BO_3_. H_2_SO_4_ was added until the solution cleared. The vitamin solution contained (g L^-1^) 10.0 g nicotinic acid, 5.0 g thiamine HCl and 0.10 g Biotin. Solid media agar plates were produced by addition of 1.5 wt.% agar. R-2A medium (**R-2A**): contained (L^-1^) 0.5 g casein acid hydrolysate, 0.5 g dextrose, 0.3 g K₂HPO₄, 0.025 g MgSO₄, 0.5 g proteose peptone, 0.3 g sodium pyruvate 0.5 g starch (soluble), 0.5 g yeast extract. **R-2A** (Merck 17209) was used for preparing agar plates. Tryptic soy broth (**TSB**): containing (L^-1^) 17 g casein peptone, 3 g soya peptone, 5 g NaCl, 2.5 g Na_2_HPO_4_ and 2.5 g dextrose (Sigma Aldrich 22092-500G). 1.5 wt. % agar plates were made with 0.1X strength TSB. Nutrient broth (**NB,** Merch): containing (L^-1^) 15 g yeast extract, 3.0 g NaCl, 1 g dextrose. 1.5 wt.% agar plates were made with 0.2X strength NB.

### 3 Enrichment culturing

#### 3.1 D-line

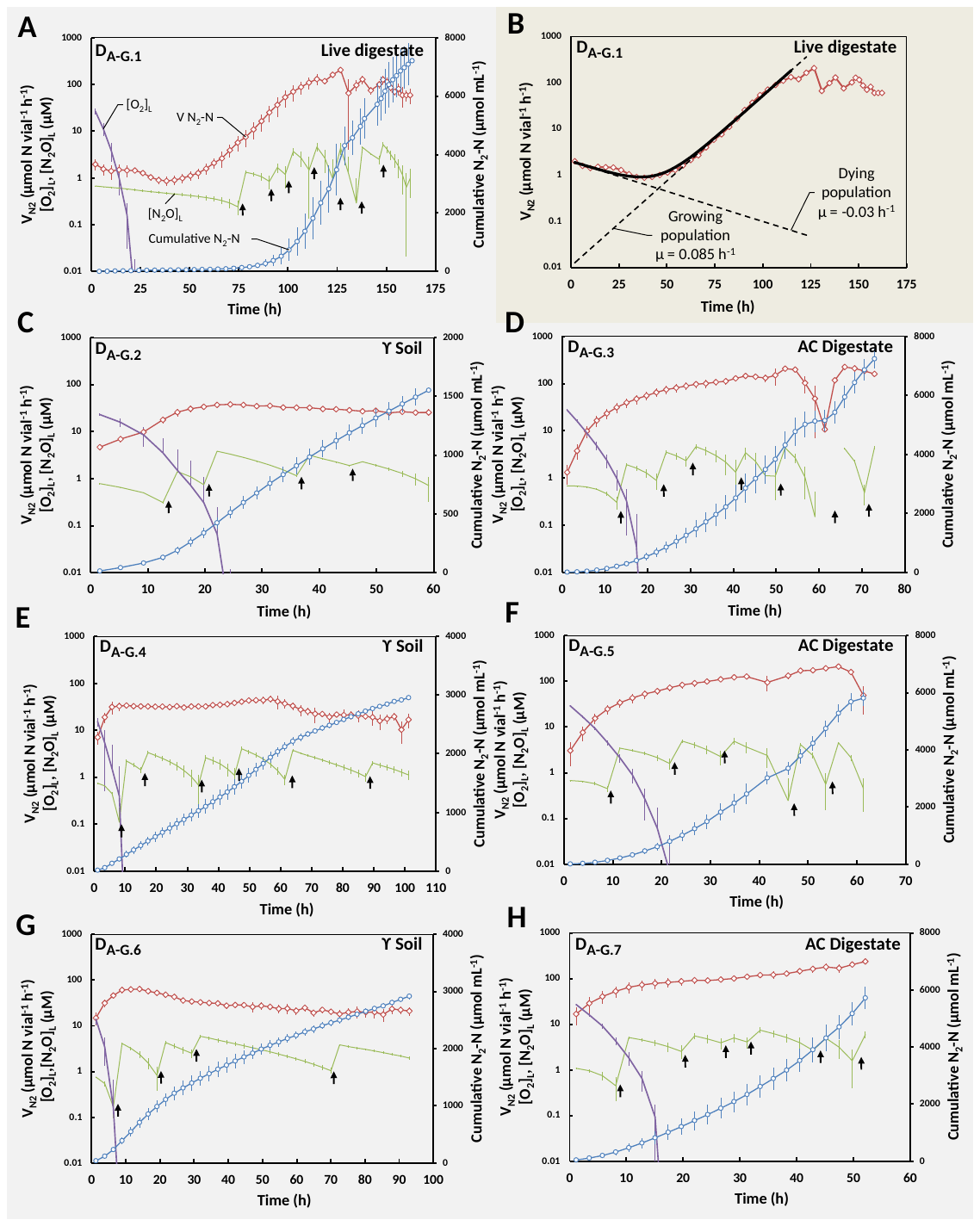

**Figure S7: Enrichment culturing starting with live digestate – (D-line).** Panel A&B shows the result for the initial enrichment culturing by anaerobic incubation of live digestate. Panel A shows N_2_-N production rates, cumulative N_2_-N produced, and liquid concentration of N_2_O-N and O_2_ throughout enrichment, while panel B shows the measured N_2_ production rate on a log scale, together with a fitted model assuming a dying and a growing population as developed by Jonassen et al (2021). The panels C-H shows the same data as in panel A, for the subsequent enrichment cultures line D_A-G.2_ to D_A-G.7_. Each enrichment culture was started by transferring 10 weight % of material from the previous enriched culture (D_A-G.j_ to D_A-G.j+1_). Black arrows: exogenous addition of N_2_O. Error bars is displayed as standard deviation (n = 7).

#### 3.2 SD-line

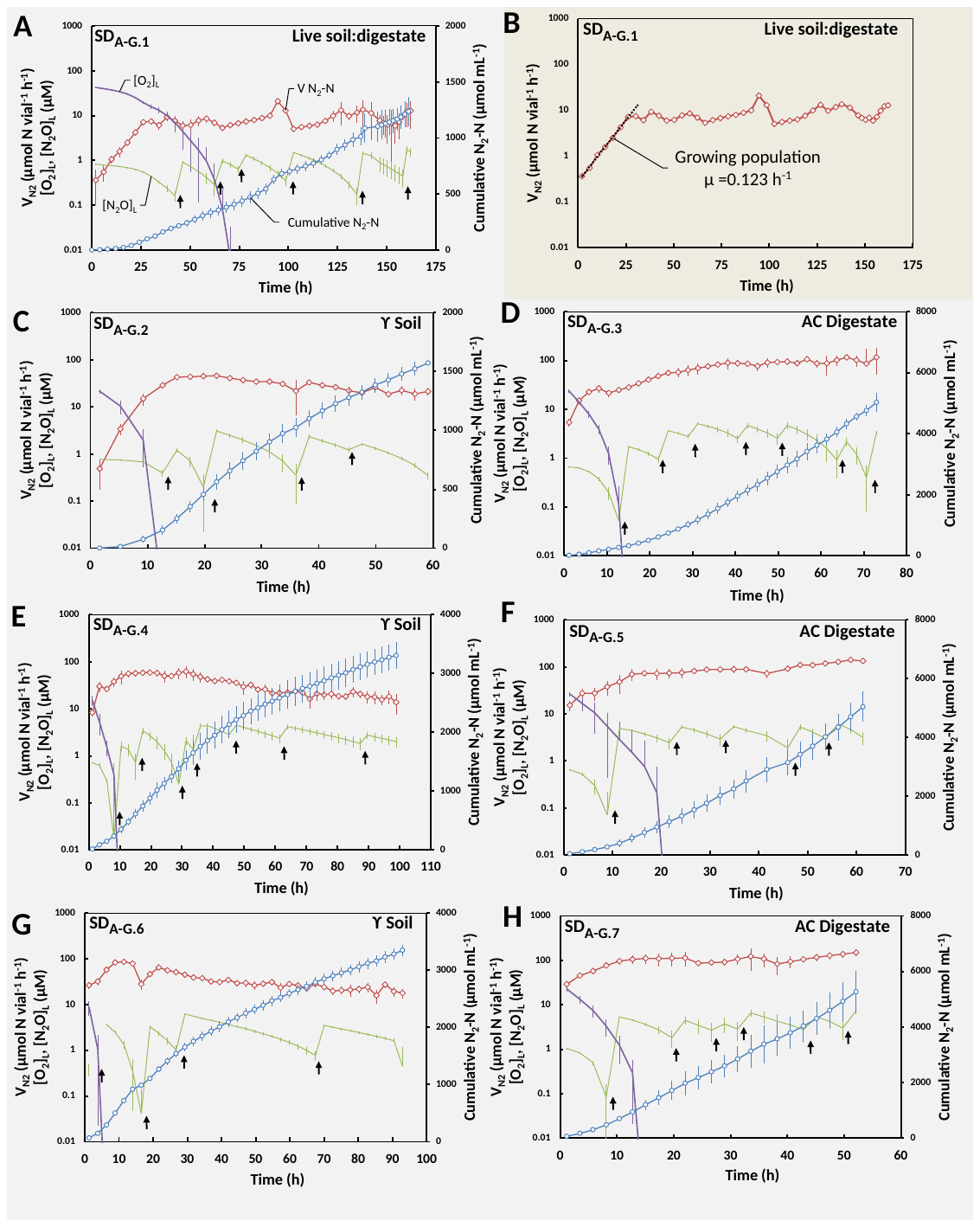

**Figure S8: Enrichment culturing starting with live digestate + live soil (SD-line).** Panel A&B shows the result for the initial enrichment culturing by anaerobic incubation of live digestate mixed with live soil. Panel A shows N_2_-N production rates, cumulative N_2_-N produced and liquid concentration of N_2_O-N and O_2_ throughout enrichment, while panel B shows the measured N_2_ production rate on a log scale. In contrast to the D line (**Fig. S7**), the N_2_ production increased exponentially from the very start. The panels C-H shows the same data as in panel A, for the subsequent enrichment cultures line SD_A-G.2_ to SD_A-G.7_. Each enrichment culture was started by transferring 10 weight % of material from previous enriched culture (SD_A-G.j_ to SD_A-G.j+1_). Black arrows: exogenous addition of N_2_O. Error bars is displayed as standard deviation (n = 7).

**Estimation of cells surviving the passage between digestate and soil**

The N_2_O reduction kinetics in the series of enrichment cultures were used to obtain crude estimates of the fraction of organisms surviving the transfer from one substrate to the next (from soil to digestate, and *vice versa*). The calculation was based on the assumption that all active organisms have equal growth yield (***Y*** = cells mol^-1^electrons) and maximum growth rate (***µ_max_***, h^-1^), hence also cell-specific maximum respiration rate (***v_max_***, mol electrons cell^-1^ h^-1^).

The estimated number of N_2_O-respiring respiring cells at the end of enrichment ***n***, $N_{n(end)}$ is:

$N_{n(end)}=\frac{V_{e_{n(0)}}}{v_{max}}+E_{n-cum}\cdot Y$	(1)

where $V_{e_{n(0)}}$is the initial rate of electron transport to N_2_O reductase for enrichment ***n*** , and $E_{n-cum}$ is the cumulated electron flow to N_2_O during enrichment ***n***.

The estimated number of N_2_O-respiring respiring cells at the beginning of the next enrichment, ***n+1***, $N_{n+1(0)}$, is:

$N_{n+1(0)}=V_{e_{n+1}(0)}/v_{max}$	(2)

where $V_{e_{n+1}(0)}$ is initial rate of electron transport to N_2_O reductase for enrichment ***n+1***.

Since 10 % of the material in culture n was transferred to culture n+1, we have that the estimated fraction of N_2_O-reducing organisms surviving this transfer, ***f***, is:

$f=N_{n+1(0)}/(0.1\cdot N_{n(end)})$	(3)

Combining equation 1,2 and 3, and the fact that *Y ·* *v_max_* = *µ_max_*, we have that

$f=10\cdot V_{e_{n+1(0)}}/(V_{e_{n}(end)} +E_{n-cum}\cdot\mu_{max})$	(4)

*While E_n-cum_* was measured for each enrichment culture, while *V_en(0)_* and *V_en+1(0)_* were not, since each enrichment was initiated with ~1.4 vol% O_2_ in the headspace, resulting in a mixture of aerobic and anaerobic respiration during the first 10-15 hours until O_2_ was depleted (**Figure S9B**). To estimate *V_en(0)_* and *V_en+1(0)_* to be used to estimate ***f*** (equation 4), the measured rates of electron flow rate to N_2_O immediately after oxygen depletion were extrapolated back to time 0, assuming exponential growth rate *µ_max_*=0.1h^-1^. The implicit assumption is the absence of any lag phase after transfer, thus $V_{e_{n+1(0)}}$ , hence ***f*** should be considered minimum estimates.

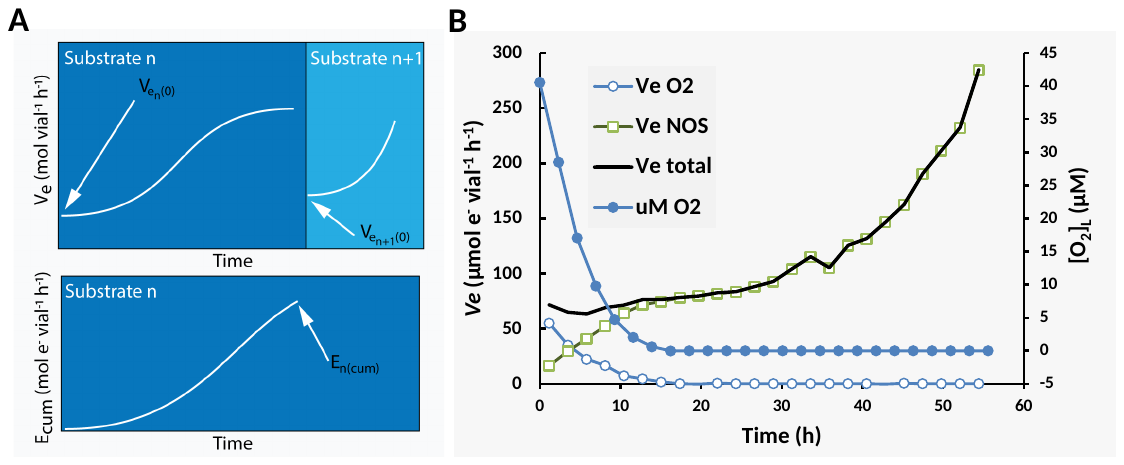

**Figure S9** Panel A: Example kinetics of how the N_2_O reduction kinetics in the series of enrichment cultures were used to estimate the % of active organisms surviving the transfer from one substrate to the next (from soil to digestate, and *vice versa*). The electron flow to N_2_O reductase increased in *Substrate n* (soil or digestate) exponentially to begin with, leveling off gradually as growth becomes substrate limited. At the end of the enrichment culturing in *Substrate n*, 10 % is transferred to enrichment culturing in *Substrate n+1* (soil after digestate, or digestate after soil). The fraction of N_2_O reducers in *Substrate n* which survives the transfer to *Substrate n+1* is to be calculated based on measured initial volumetric electron flow rate to N_2_O (***V_e n+1_***) in *Substrate n+1*, and the *Substrate n*- data: initial volumetric electron flow rate to N_2_O (***V_e_* _n_**) and the cumulated electron flow (***E_n-cum_***) using equation 7 (see text preceding Fig S4). Panel B: The panel shows an example for enrichment culture vial D_A.1_ (enrichment line A in digestate, enrichment number 1): The measured oxygen concentration declined rapidly during the first 10 hours. The estimated electron flow rate to oxygen (Ve O2) declined accordingly and was gradually replaced by the electron flow to N_2_O (Ve NOS). Ve total is the sum of the electron flow to oxygen and N_2_O.

#### 3.3 SIMPER analysis on 16S amplicon data

**Table S4**: SIMPER analysis results output of the top 10 OTUs contribution to the explained variance in the D lines.

| Taxon | Clade | Avgerage dissimil. | Contrib. (%) | Cumulat. (%) | Mean abundance (%) | | | | | | |
| --- | --- | --- | --- | --- | --- | --- | --- | --- | --- | --- | --- |
|  |  |  |  |  | SD_A-G.1_ | SD_A-G.2_ | SD_A-G.3_ | SD_A-G.4_ | SD_A-G.5_ | SD_A-G.6_ | SD_A-G.7_ |
| OTU1 | A | 0.0659 | 48.5 | 48.5 | 0.228 | 0.155 | 0.127 | 4.96 | 7.58 | 41.4 | 32.9 |
| OTU3 | C | 0.0196 | 14.4 | 62.9 | 26.1 | 9.19 | 0.20 | 0.074 | 0.017 | 0.009 | 0.015 |
| OTU6 | A | 0.0131 | 9.64 | 72.6 | 0.104 | 6.29 | 21.9 | 20.0 | 13.9 | 8.27 | 8.36 |
| OTU2 | A | 0.0116 | 8.56 | 81.1 | 8.56 | 18.8 | 28.6 | 15.8 | 22.5 | 7.87 | 17.1 |
| OTU4 | C | 0.0064 | 4.70 | 85.8 | 15.0 | 5.08 | 0.229 | 0.054 | 0.011 | 0.007 | 0.008 |
| OTU5 | A | 0.0046 | 3.35 | 89.2 | 0.059 | 0.124 | 7.30 | 4.78 | 13.2 | 3.52 | 7.35 |
| OTU8 | A | 0.0038 | 2.78 | 92.0 | 0.039 | 0.031 | 0.014 | 2.32 | 2.91 | 6.58 | 6.98 |
| OTU14 | A | 0.0014 | 1.01 | 93.0 | 0.035 | 0.34 | 5.05 | 1.20 | 5.17 | 0.60 | 4.20 |
| OTU7 | C | 0.0010 | 0.72 | 93.7 | 5.28 | 3.53 | 0.067 | 0.023 | 0.004 | 0.004 | 0.003 |
| OTU29 | A | 0.0009 | 0.68 | 94.4 | 0.023 | 1.72 | 1.5 | 5.39 | 2.01 | 4.63 | 1.23 |

**Table S5**: SIMPER analysis results output of the top 10 OTUs contribution to the explained variance in the SD lines.

| Taxon | Clade | Avgerage dissimil. | Contrib. (%) | Cumulat. (%) | Mean abundance (%) | | | | | | |
| --- | --- | --- | --- | --- | --- | --- | --- | --- | --- | --- | --- |
|  |  |  |  |  | SD_A-G.1_ | SD_A-G.2_ | SD_A-G.3_ | SD_A-G.4_ | SD_A-G.5_ | SD_A-G.6_ | SD_A-G.7_ |
| OTU1 | A | 0.1033 | 61.9 | 61.9 | 0.14 | 4.2 | 14.8 | 47.0 | 43.5 | 55.3 | 39.3 |
| OTU11 | A | 0.0133 | 7.98 | 69.9 | 2.30 | 11.2 | 4.94 | 1.46 | 2.85 | 2.19 | 11.7 |
| OTU2 | A | 0.0105 | 6.31 | 76.2 | 0.36 | 2.86 | 20.7 | 5.13 | 13.4 | 3.98 | 8.91 |
| OTU6 | A | 0.0061 | 3.63 | 79.9 | 0.05 | 4.66 | 17.7 | 7.71 | 8.11 | 4.32 | 6.0 |
| OTU3 | C | 0.0057 | 3.42 | 83.3 | 14.10 | 4.61 | 0.134 | 0.023 | 0.014 | 0.010 | 0.008 |
| OTU4 | C | 0.0053 | 3.16 | 86.4 | 13.30 | 4.2 | 0.205 | 0.020 | 0.008 | 0.006 | 0.003 |
| OTU17 | A | 0.0040 | 2.37 | 88.8 | 0.25 | 12.1 | 2.21 | 2.15 | 0.321 | 0.441 | 0.106 |
| OTU7 | C | 0.0031 | 1.87 | 90.7 | 10.30 | 4.08 | 0.156 | 0.014 | 0.004 | 0.004 | 0.002 |
| OTU5 | A | 0.0029 | 1.72 | 92.4 | 0.03 | 0.469 | 10.7 | 2.62 | 5.93 | 1.82 | 4.24 |
| OTU8 | A | 0.0019 | 1.14 | 93.5 | 0.01 | 0.118 | 0.694 | 3.76 | 4.17 | 6.05 | 6.02 |

### 4 Genome sequencing, phylogeny and *eco-physiological genome analysis* of isolated organisms

**Table S6**: SMRT® link software (PacBio®) output (assembly parameters, alignment to draft assembly, polished assembly, coverage) and CheckM calculated completeness (presence of single copy marker genes) and contamination (multiple single copy marker genes) of isolated organisms.

| Isolates: | CB-01 | CB-03 | BM | AM | PS-02 | OB |
| --- | --- | --- | --- | --- | --- | --- |
| Assembly Parameters | | | | | | |
| Method | HGAP4 | HGAP4 | HGAP4 | HGAP4 | Microbial Assembly | Microbial Assembly |
| Seed Coverage | 30 | 30 | 30 | 22 | 20 | 15 |
| Expected Genome Length | 2 740 000 | 2 740 000 | 2 710 000 | 4 630 000 | 4 340 000 | 4 830 000 |
| Alignment to Draft Assembly | | | | | | |
| Percent Aligned Bases | 95.41% | 92.53% | 95.66% | 84.55 % | 93.60 % | 76.39 % |
| Number of Subreads (aligned) | 202 702 | 304 312 | 456 704 | 226 629 | 166 555 | 192 289 |
| Number of Polymerase Reads (aligned) | 13 088 | 20 485 | 43 392 | 18 713 | 14 278 | 14 180 |
| Polymerase Read Length Mean (aligned) [bp] | 50 243 | 50 040 | 37 734 | 38 599 | 38 419 | 42 567 |
| Polymerase Read Length Max (aligned) | 134 672 | 130 398 | 121 149 | 122 221 | 125 758 | 123 538 |
| Polished Assembly | | | | | | |
| Polished Contigs | 1 | 1 | 2 | 1 | 5 | 54 |
| Maximum Contig Length [bp] | 2 979 886 | 2 718 917 | 2 711 532 | 4 571 002 | 4 016 625 | 297 087 |
| Sum of Contig Lengths [bp] | 2 979 886 | 2 718 917 | 2 754 828 | 4 571 002 | 4 494 782 | 4 640 821 |
| Coverage | | | | | | |
| Mean Coverage | 207 | 354 | 552 | 146 | 113 | 120 |
| Missing bases (%) | 0.00 % | 0.00 % | 0.00 % | 0.00 % | 0.00 % | 0.00 % |
| CheckM quality parameters | | | | | | |
| Completeness | 100 % | 100 % | 99.77 % | 99.97 % | 99.96 % | 80.32 % |
| Contamination | 0.74 % | 0.25 % | 0 % | 0 % | 1.16 % | 0.43 % |

**Table S7**: Isolates of N_2_O reducing bacteria isolated from the final enrichment (samples D_7_ and SD_7_) in autoclaved digestate: The isolates were circumscribed by OTUs based on 16S identity (> 97 %). Average OTU abundance (7 biological replicates ± standard deviation) is shown for each OTU circumscribing isolated organisms for enrichment D _A-G.1_/SD _A-G.1_ (first enrichment in live material), D _A-G.1_/SD _A-G.1_ (last gamma sterilized soil enrichment) and D _A-G.1_/SD _A-G.1_ (final enrichment in autoclaved digestate).

| Isolate: | Circumscribed by OTU | Clade | % 16S rDNA seq. ident. (Overlapping region: 404 - 429 bp) | Average OTU abundance (%) ± standard deviation (n = 7) | | | | | |
| --- | --- | --- | --- | --- | --- | --- | --- | --- | --- |
|  |  |  |  | D_A-G.1_ | D_A-G.6_ | D_A-G.7_ | SD_A-G.1_ | SD_A-G.6_ | SD_A-G.7_ |
| *Cloacibacterium* sp. **CB-01** | 1 | A | 99.8 % | 0.23 ± 0.02 | 41.4 ± 9.6 | 32.9 ± 10.3 | 0.14 ± 0.04 | 55.3 ± 2.0 | 39.3 ± 9.6 |
| *Cloacibacterium* sp. **CB-03** | 1 | A | 99.8 % |  |  |  |  |  |  |
| *Azonexus* sp. **AN*** | 2 | A | 98.2 % | 8.56 ± 0.54 | 7.87 ± 1.82 | 17.1 ± 2.87 | 0.36 ± 0.16 | 17.1 ± 2.87 | 8.91 ± 3.12 |
| *Pseudomonas*  sp. **PS-02** | 8 | A | 99.8 % | 0.04 ± 0.05 | 6.58 ± 4.85 | 6.98 ± 5.52 | 0.011 ± 0.002 | 3,98 ± 0.55 | 6.02 ± 3.12 |
| *Aeromonas*  sp. **AM** | 19 | A | 99.5 % | 0.031 ± 0.026 | 2.15 ± 2.17 | 2.26 ± 2.27 | 0.017 ± 0.007 | 6.05 ± 1.74 | 4.49 ± 0.97 |
| *Brachymonas*  sp. **BM** | 37 | A | 100 % | 0.004 ± 0,003 | 1.125 ± 0.313 | 0.63 ± 0.12 | 0.0075 ± 0.0065 | 3.68 ± 0.96 | 0.308 ± 0.159 |
| *Ochrobactrum* sp. **OB** | 74 | D | 100 % | 0.003 ±0,003 | 0.227 ± 0.055 | 0.003 ± 0.002 | 0.0017 ± 0.0013 | 0.57 ± 0.15 | 0.005 ±  0.003 |

* The genome of *Azonexus sp*. AN was not sequenced: 16S rDNA sequence identity with OTUs was determined by alignment of 16S OTU sequence and 16S from Sanger sequencing of PCR amplicons amplified using 27F and 1492R universal primer pairs (see main text).

**Table S8:** ddPCR quantification of 16S copy numbers on pooled samples (A to G) of DNA extracts from D_A-G.j_ and SD_A-G.j_ (j = 1-7), D0 and SD0, and sterile growth substrates used throughout the enrichment (AC-dig and ϒ-Soil) and standard error (n = 3).

| **Line** | **Material:** | **Sample:** | **16S/vial:** | **Stdev/sqrt(n)** |
| --- | --- | --- | --- | --- |
| D | Live digestate | D_0_ | 3.50E+11 | 7.0E+09 |
|  | Live digestate | D_A-G.1_ | 1.80E+11 | 4.7E+09 |
|  | ϒ Soil | D_A-G.2_ | 1.40E+11 | 7.9E+09 |
|  | AC-Dig | D_A-G.3_ | 1.60E+11 | 4.4E+09 |
|  | ϒ Soil | D_A-G.4_ | 2.10E+11 | 1.4E+09 |
|  | AC-Dig | D_A-G.5_ | 2.00E+11 | 7.1E+09 |
|  | ϒ Soil | D_A-G.6_ | 2.50E+11 | 7.4E+09 |
|  | AC-Dig | D_A-G.7_ | 2.00E+11 | 6.3E+09 |
| SD | Live dig:soil mix | SD_0_ | 2.60E+11 | 2.1E+09 |
|  | Soil:mix after enr. | SD_A-G.1_ | 1.60E+11 | 9.9E+09 |
|  | ϒ Soil | SD_A-G.2_ | 1.10E+11 | 6.9E+09 |
|  | AC-Dig | SD_A-G.3_ | 1.10E+11 | 5.0E+09 |
|  | ϒ Soil | SD_A-G.4_ | 2.60E+11 | 9.7E+09 |
|  | AC-Dig | SD_A-G.5_ | 1.70E+11 | 7.3E+09 |
|  | ϒ Soil | SD_A-G.6_ | 2.70E+11 | 3.0E+09 |
|  | AC-Dig | SD_A-G.7_ | 1.90E+11 | 4.0E+09 |
| Growth substrate | AC-Dig | Growth substrate | 7.00E+10 | 6.8E+09 |
|  | ϒ Soil | Growth substrate | 1.60E+10 | 1.8E+08 |

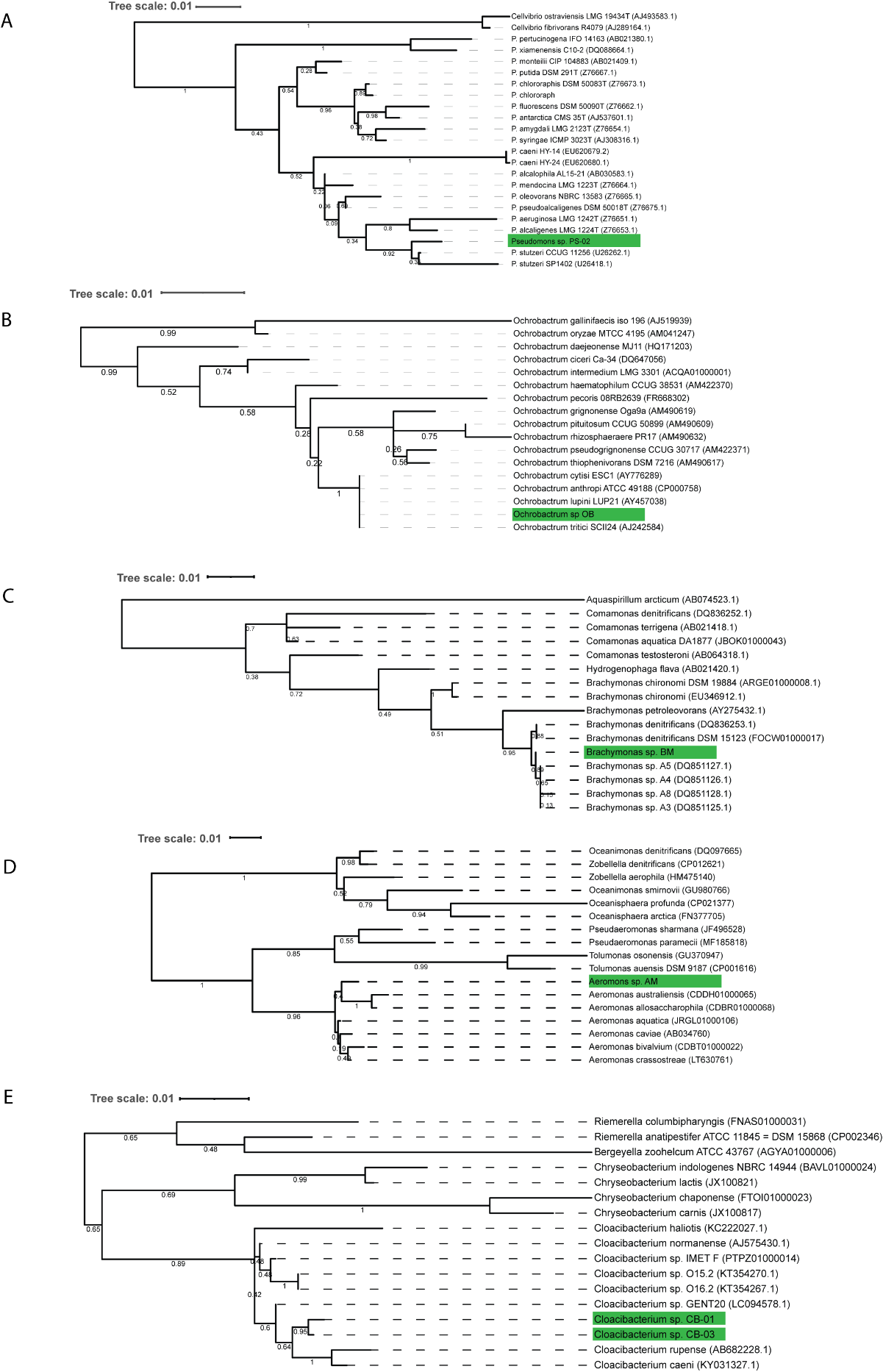

**Figure S10**: Representative characterized strains and close relatives of the isolated organisms were used to build a neighbor‐joining tree (100 bootstrap samplings) from a ClustalW alignment of 16S rDNA sequences. Numbers represent the percentage of bootstrap samplings that generate at each node. Species names are followed by the accession numbers of their 16S rDNA sequences. Panel A: *Pseudomonas* sp. PS-02 (green box), Panel B: *Ochrobactrum* sp. OB (green box), Panel C: *Brachymonas* sp. BM (green box), Panel D: *Aeromonas* sp. AM (green box) and Panel E: *Cloacibacterium* sp. CB-01 and CB-03 (green boxes).

**Table S9:** The identified proteins predicted as carbohydrate-active enzymes (CAZymes) in the genomes of AM, BM, CB-01, CB-03, PS-02, and OB. The CAZymes were automatically annotated through the dbCAN meta server (Feb 2021), which integrates three tools/databases (i.e. HMMER, DIAMOND, Hotpep) and SignalP. The CAZymes assignment includes the enzyme classes Glycoside Hydrolysis (GHs), Glycosyl transferases (GTs), Polysaccharide lyases (PLs), Carbohydrate Esterases (CEs) and enzymes with Auxiliary Activity (AAs) in addition to Carbohydrate-Binding Modules (CBM). Identical annotation in ≥ 2 tools was required for a CAZY assignment to be considered robust. Prokka annotations of corresponding genes is given in **Supplementary Data S1**.

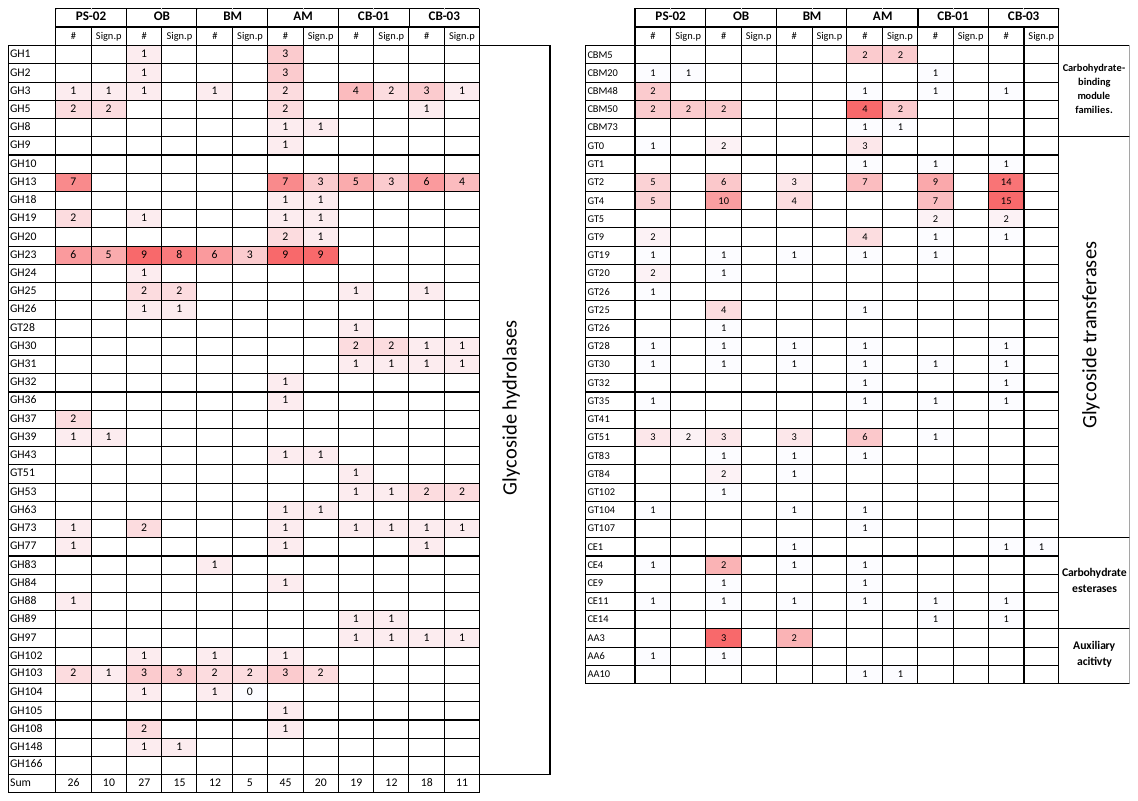

**Table S10:** A selection of identified enzymes predicted as carbohydrate-active enzymes (CAZymes) from the dbCAN meta-server and corresponding PROKKA annotations of enzymes targeting extracellular carbohydrates (indicated by identified signal sequence for membrane trans allocation (Signal P)) and genes encoding proteins involved in glycogen metabolism in the genomes of PS-02, OB, BM, AM, CB-01 and CB-03. BM did not contain annotated CAZymes of particular relevance. A complete list of dbCAN grouped genes is given together with corresponding Prokka annotations for each genome in **Supplementary Data S1**

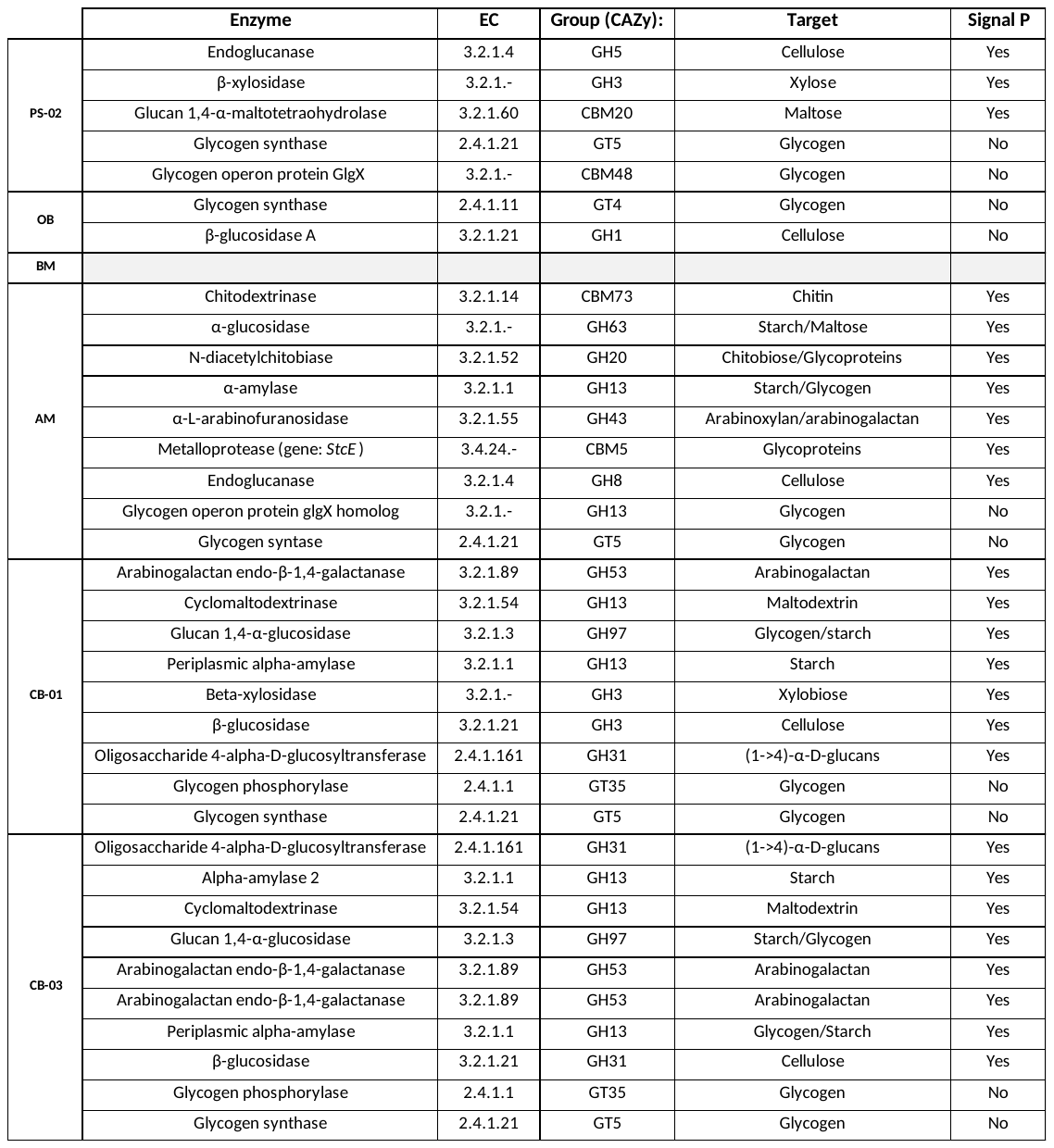

**Table S11:** MEROPS annotated peptidases in the genomes of PS-02, OB, BM, AM, CB-01 and CB-03. Every protein sequence was screened for presence of signal sequences in SignalP 5.0 and the MEROPS subfamilies were collapsed to families. Prokka annotations of corresponding genes is given in **Supplementary Data S1**

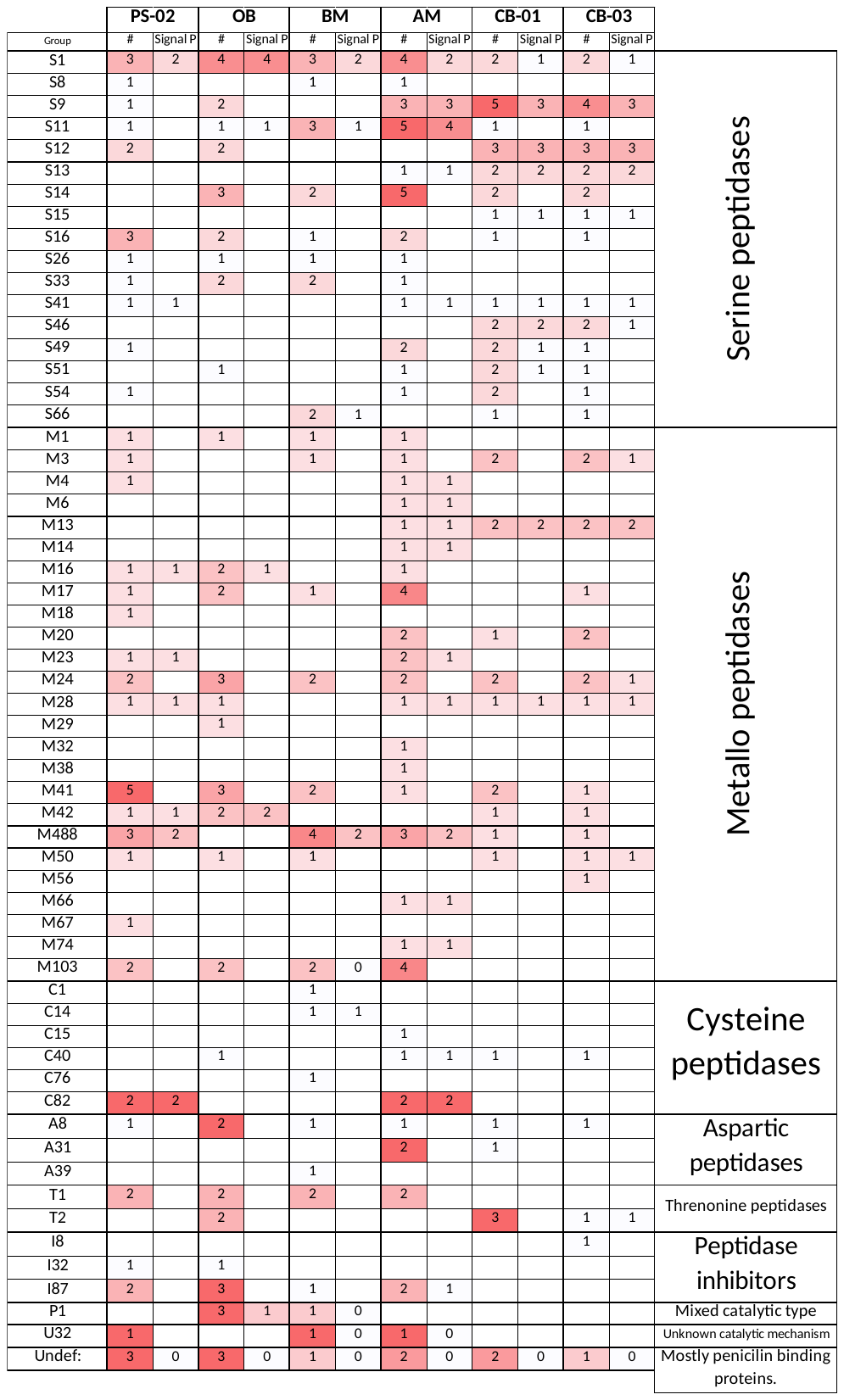

**Table S12**: Prokka annotated genes coding for proteins involved nodule formation and nitrogenase activity in genome of *Ochrobacter* sp. OB*.

| Gene: | Detected: | Onthology: | Reference: |
| --- | --- | --- | --- |
| *nodN* | Yes |  | Surin and Downing 1988, Baey et al 1992 |
| *nodM* | Yes | Glutamine-fructose-6-phosphatase (EC: 2.6.1.16) | Surin and Downing 1988, Baey et al 1992 |
| *CC_0717* | Yes | Nodulin related protein CC_0717. Unknown function. |  |
| *nifS* | Yes | Cysteine desuluronase (EC: 2.8.1.7) | Kennedy and Dean 1992 |
| *nifU* | Yes | NifU-like domain protein | Kennedy and Dean 1992 |
| *nifDKH* | No | Catalytic subunits of nitrogenase | McGlynn et al 2013 |
| *anfG* | No | Catalytic subunits of nitrogenase | McGlynn et al 2013 |
| *vnfDKGH* | No | Catalytic subunits of nitrogenase | McGlynn et al 2013 |
| *fixK* | Yes | Nitrogenase transcriptional regulator | Li et al 2010 |
| *fixN* | Yes | ec:1.9.3.1- Cytochrome-c oxidase. | David et al 1988 |
| *fixL* | Yes | Sensor protein fixL (EC 2.7.3.-) | Monson et al 1995. |

* Quality checking the assembled genome of OB with CheckM did reveal that only 80 % of the single copy marker genes were recovered from the genome of OB (**Tab. S8**), and it is conceivable that potential missing parts of the genome could contain more nod- and nitrogen fixation related genes, as well as other genetic encoded traits as discussed in main text.

### 5 Assessment of growth or death of OTU’s

To assess growth or decline of each OTU within the 6 clades, we calculated the relative increase for each consecutive enrichment culture as *R*= ln(_N(i)_/(N_(i-1)_*0.1) where N_i_ is the estimated copy number per vial at the end of enrichment i and N_(i-1)_ is the estimated copy numbers at the end of the foregoing enrichment (both estimated by the relative abundance of the OTU in question and the total copy numbers of 16SrDNA quantified by digital PCR (**Tab. S10**). The multiplication with 0.1 is because 10% of the content of one enrichment culture was transferred to the next. *R* for the initial enrichment in live digestate (with and without live soil added is ln(N/N_0_), where N is the abundance after enrichment and N_0_ is the abundance measured at the onset of this enrichment). The result for the 6 clades is shown in **Figs. S11-S16.**

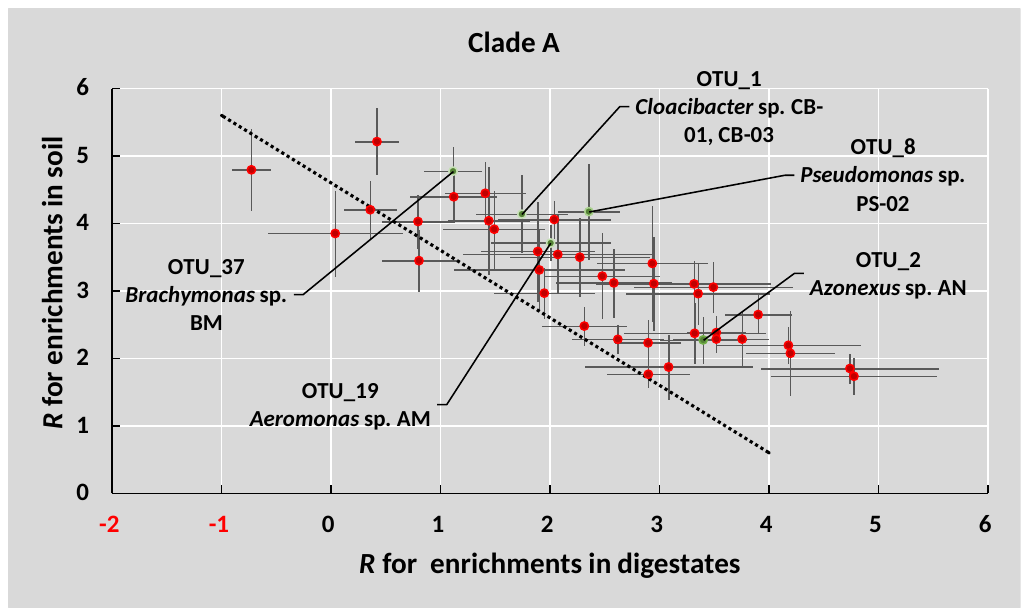
**Figure S11 Assessment of growth/death (soil versus digestate) of OTU’s within Clade A**. For the 42 OTUs of Clade A the increase or decrease in copy numbers for each individual enrichment vial was estimated as *R*= ln(_N(i)_/(N_(i-1)_·0.1). For each OTU, the average *R* for the enrichments in autoclaved digestate (*R_digestate_*) and γ-sterilized soil (*R_soil_*) were calculated. The panel shows *R_soil_* plotted against *R_digestate_*, with standard error (n=8) marked by vertical and horizontal lines. The dashed line is the plotted equation *R_soil_*+*R_digestate_*= 4.6, which marks a division between OTU’s that are sustained (R_soil_+R_digestate_ > 4.61) or gradually washed out (R_soil_+R_digestate_ < 4.61) throughout the dual enrichment cultures. The plotting of *R_soil_* and *R_digestate_* for individual OTU’s shows that the OTU’s within Clade A span a continuum from “Soil specialists” (high *R_soil_*, low/negative *R_digestate_* values) through “Generalists” (similar *R_soil_* and *R_digestate_* values >2) and further on to “Digestate specialists” (low R_soil_, high R _dig_). Interestingly several of the isolates qualifies as Generalists, while *Azonexus* is more of a Digestate specialist, as suspected (Jonassen et al 2021). *R_soil_* was negatively correlated with *D_digestate_* (regression function: *R_soil_*=4.7-0.6·*R_digestate_*, r^2^=0.7, p<0.01).

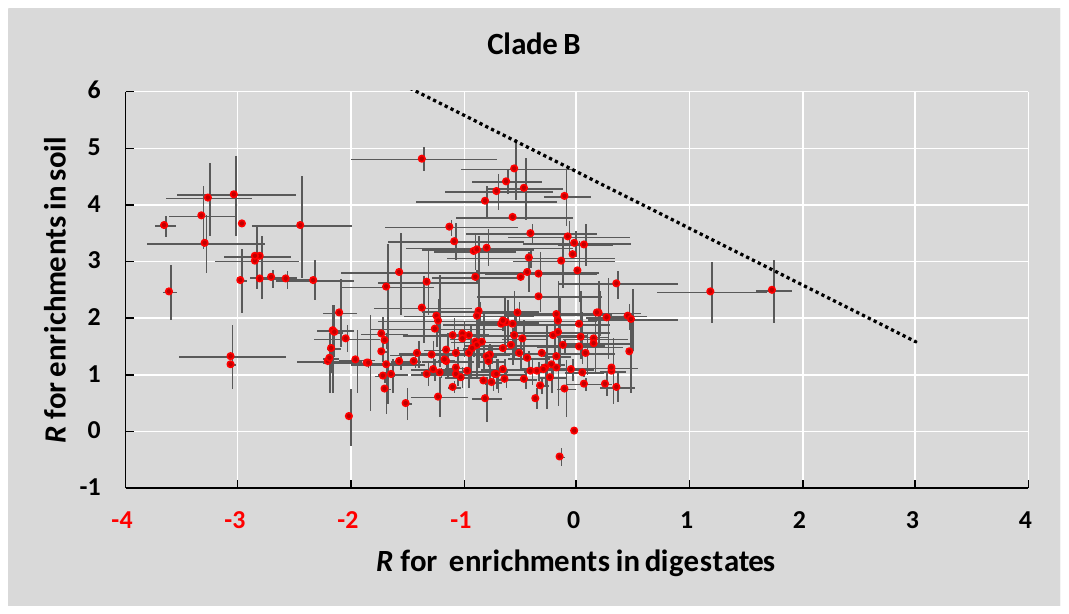

**Figure S12: Assessment of growth/death (soil versus digestate) of OTU’s within Clade B**. For the 172 OTUs of Clade B the increase or decrease in copy numbers was estimated as R= ln(_N(i)_/(N_(i-1)_·0.1). For each OTU, the average R for the enrichments in autoclaved digestate and γ-sterilized soil were calculated. The panel shows R for Soil plotted against R for Digestate, with standard error (n=8) marked by vertical and horizontal lines. The dashed line marks the division between cells that are predicted to die out: R_soil_+R_digestate_ >4.61 for organisms that will increase throughout and <4.61 for organisms that will decline throughout. All OTU’s within this clade are below the line. The distribution of R values suggests the majority dies out due to failure in the digestate, rather than in soil.

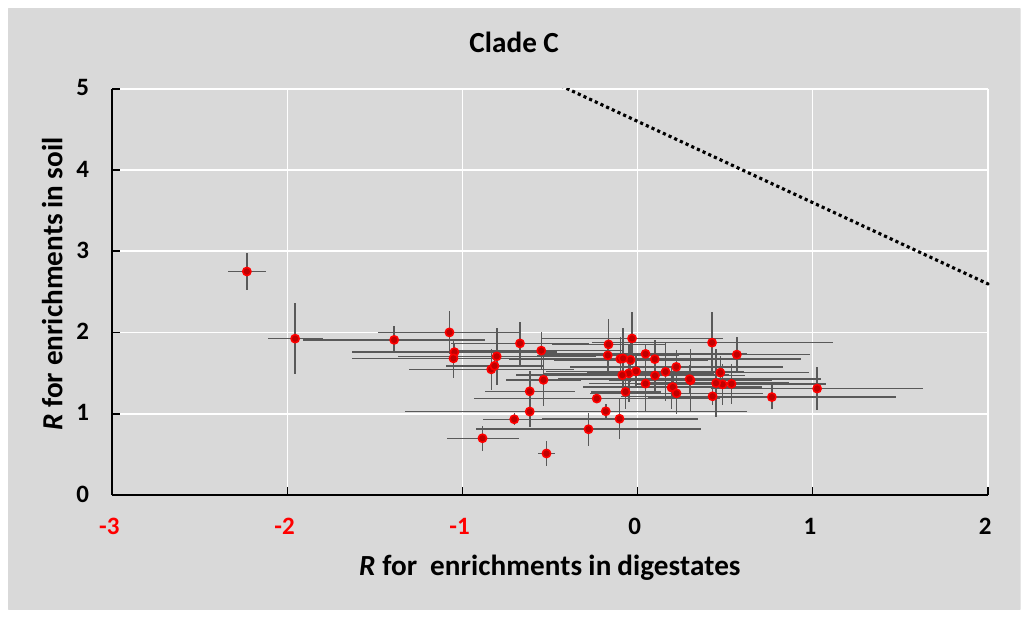
**Figure S13: Assessment of growth/death (soil versus digestate) of OTU’s within Clade C**. For the 51 OTUs of Clade C the increase or decrease in copy numbers was estimated as R= ln(_N(i)_/(N_(i-1)_·0.1). For each OTU, the average R for the enrichments in autoclaved digestate and γ-sterilized soil were calculated. The panel shows R for Soil plotted against R for Digestate, with standard error (n=8) marked by vertical and horizontal lines. The dashed line marks the division between cells that are predicted to die out: R_soil_+R_digestate_ >4.61 for organisms that will increase throughout and <4.61 for organisms that will decline throughout. All OTU’s within this clade are below the line.

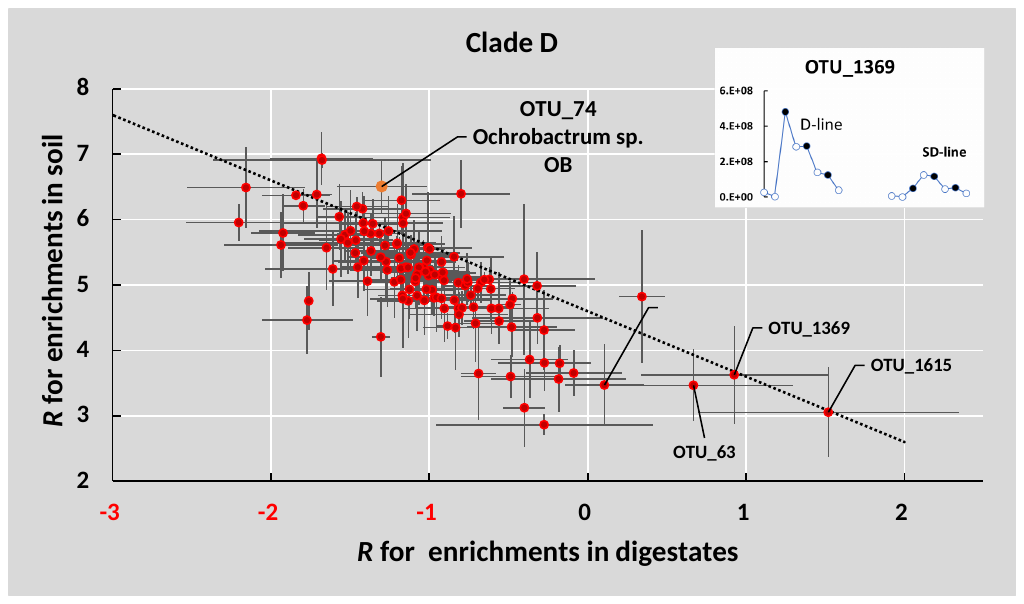
**Figure S14: Assessment of growth/death (soil versus digestate) of OTU’s within Clade D**. For the 128 OTUs of Clade D the increase or decrease in copy numbers was estimated as R= ln(_N(i)_/(N_(i-1)_*0.1). For each OTU, the average R for the enrichments in autoclaved digestate and γ-sterilized soil were calculated. The panel shows R for Soil plotted against R for Digestate, with standard error (n=8) marked by vertical and horizontal lines. For digestate, the majority of OTU’s had R between -2 and -1.5, which indicates a 80-85% decline during the enrichment in digestate. In contrast, R for enrichments in soil ranged from 3-7, indicating that the abundance increased by a factor of 150-1100 (=7-10 cell divisions) during the enrichment in soil. A clear negative correlation between R_soil_ and R_digestate_ is observed (r^2^=0.573, p<0.01). The outliers with apparent growth in digestate (R>0) showed somewhat erratic development of abundance throughout, as illustrated for OTU_1369 (inserted panel). The dashed line marks the division between cells that are predicted to die out (R_soil_+R_digestate_ >4.61 for organisms that will increase throughout and <4.61 for organisms that will decline throughout). The majority is below the line (= declining), while some are sustained or grow slightly, which was the case for the OTU 74 circumscribing the isolated *Ochrobactrum* sp. OB. The distribution of R values suggests that the majority dies out fast in the digestate but grow fast in soil.

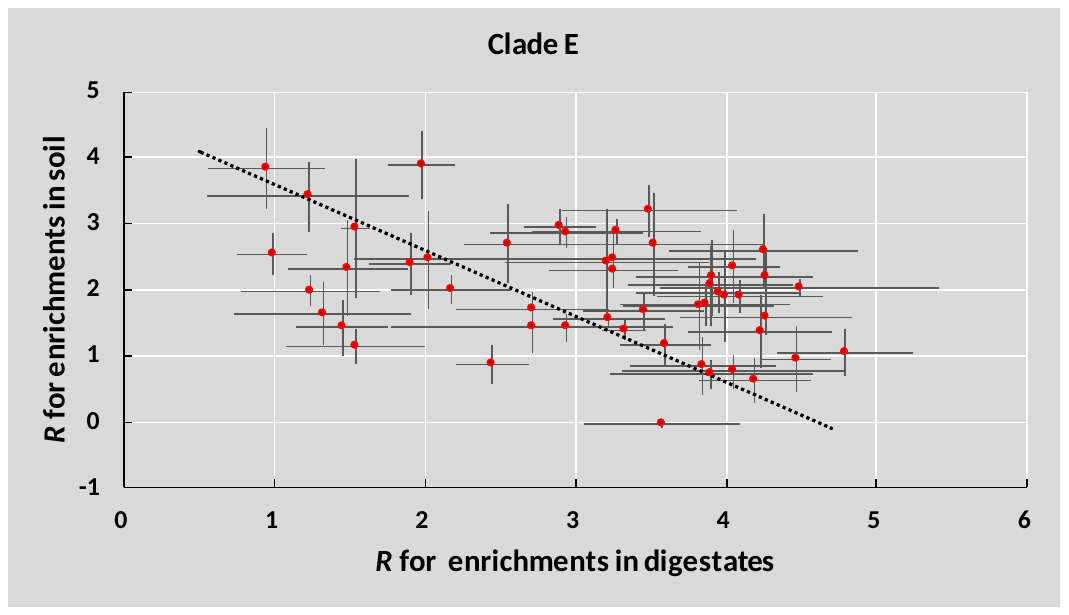

**Figure S15: Assessment of growth and death of OTU’s within Clade E**. For the 50 OTUs of Clade E the increase or decrease in copy numbers was estimated as R= ln(_N(i)_/(N_(i-1)_*0.1). For each OTU, the average R for the enrichments in autoclaved digestate and γ-sterilized soil were calculated. The panel shows R for Soil plotted against R for Digestate, with standard error (n=8) marked by vertical and horizontal lines. The dashed line marks the division between cells that are predicted to die out: R_soil_+R_digestate_ >4.61 for organisms that will increase throughout and <4.61 for organisms that will decline throughout. 35 of the 50 OTU’s within this clade are above the line.

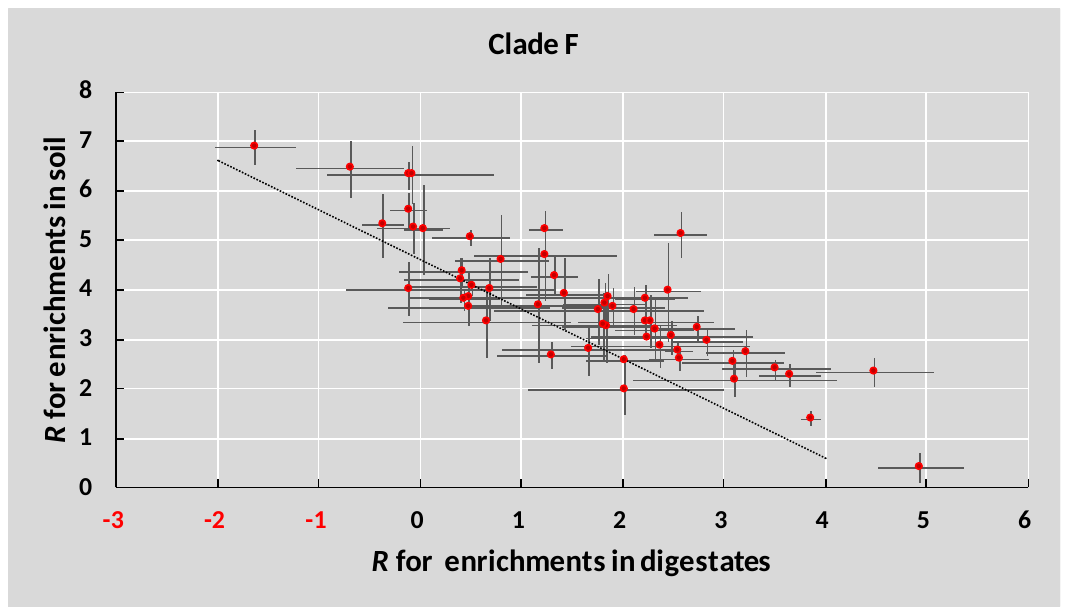

**Figure S16: Assessment of growth and death of OTU’s within Clade F**. For the 57 OTUs of Clade F the increase or decrease in copy numbers was estimated as R= ln(_N(i)_/(N_(i-1)_·0.1). For each OTU, the average *R* for the enrichments in autoclaved digestate and γ-sterilized soil were calculated. The panel shows *R* for Soil (*R_soil_*) plotted against *R* for Digestate (*R_digestate_*), with standard error (n=8) marked by vertical and horizontal lines. The dashed line marks the division between cells that are predicted to die out: *R_soil_*+*R_digestate_* >4.61 for organisms that will increase throughout and <4.61 for organisms that will decline throughout. 46 of the 57 OTU’s within this clade are above the line. *R_soil_* was negatively correlated with *D_digestate_* (regression function: *R_soil_*=5.0-0.8·*R_digestate_*, r^2^=0.7, p<0.01).

### 6 Denitrifying phenotype experiments

#### 6.1 *Pseudomonas sp.* PS-02

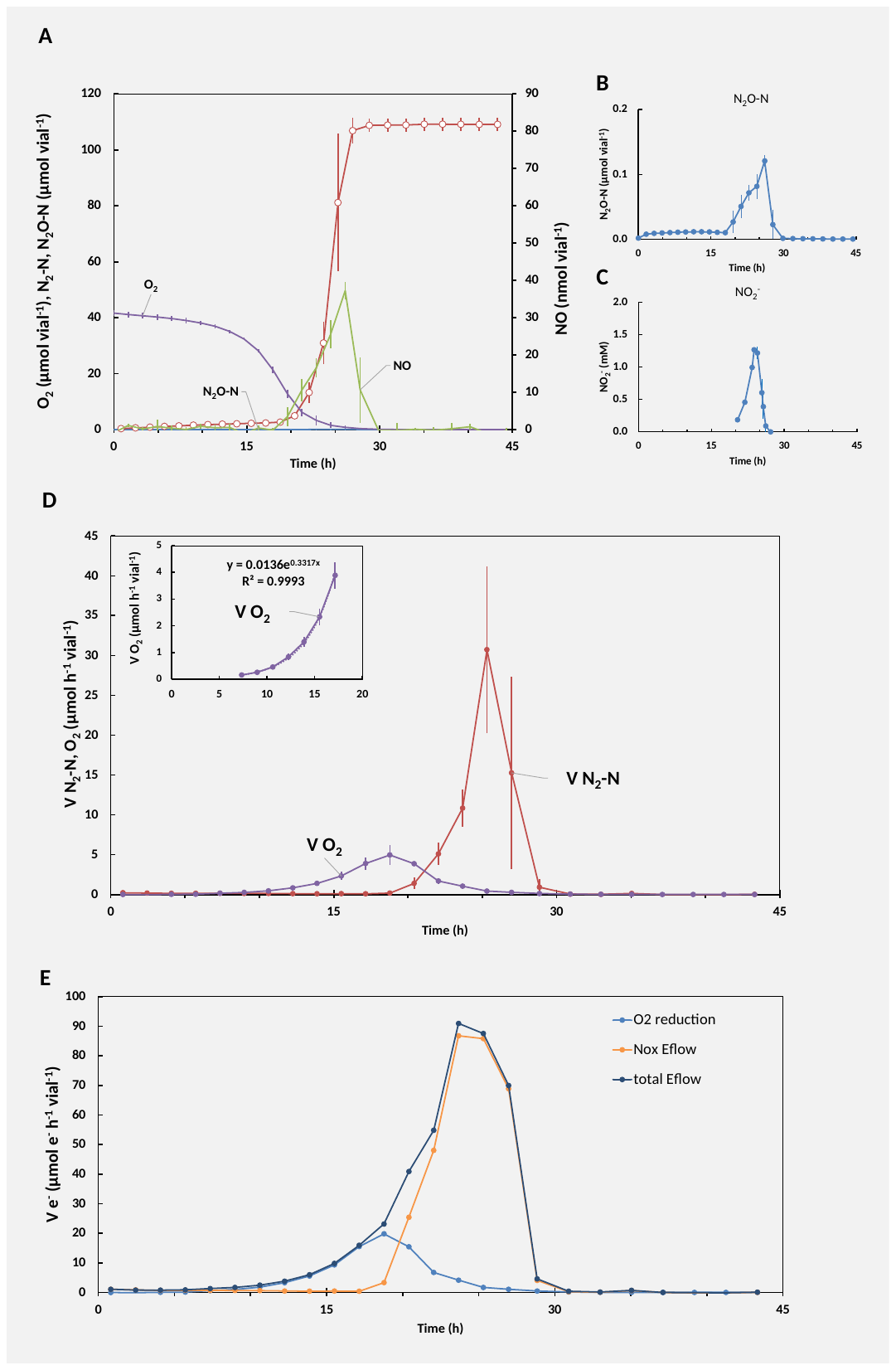

The genome analysis of *Pseudomonas sp*. PS-02 (**Fig. 4A in the main paper**) revealed genes coding for all denitrification-related reductases involved in the sequential reduction of NO_3_^-^ to N_2_: cytoplasmic *nar*, *nirS*, *nor* and *nosZ* (clade I), thus predicting a full-fledged denitrifier. In liquid culture supplemented with NO_3_^-^ or NO_2_^-^ and O_2_ cells transitioned seamlessly from oxic to anaerobic respiration on NO_3_^-^ (**Fig. S17**) or NO_2_^-^ (**Fig. S18**) whilst maintaining strict control of gaseous intermediates NO and N_2_O. NO_2_^-^ accumulated to mM levels when supplied with 2 mM NO_3_^-^ (**Fig. S17**). Cells favored respiration of exogenously supplied N_2_O over NO_3_^-^ (**Fig. 4A main paper**, supplementing kinetics in **Fig. S19**) and NO_2_^-^ (**Fig. S20**), which might indicate that the relative activity of Nos was higher than the other N-reductases at the oxic/anoxic transition, or a delay of Nar (and Nir) expression succeeding the oxic/anoxic transition relative to Nos, as competition for electrons between Nar and Nos was not expected. The strict control of N_2_O under growth on NO_3_^-^ and NO_2_^-^, and with an apparent over capacity for N_2_O respiration over NO_3_^-^ would deem PS-02 as a strong N_2_O sink and an interesting candidate for inoculation of soils.

**Figure S17 Denitrification phenotype of *Pseudomonas* sp. PS-02 when provided with O_2_ and NO_3_.** PS-02 was grown in gas-tight 120 mL vials with 50 mL Sistrom’s succinate medium initially supplemented with 1 mL O_2_ and 2mM NO_3_^-^ at constant temperature and stirring (20 °C, 600 rpm). Initial OD_660_ ≈ 0.001. Panel A**:** Measured gases (N_2_O, NO, O_2_) and calculated cumulative N_2_ throughout the incubation. Panel B: Measured N_2_O throughout the incubation. Panel C: Measured liquid concentration of NO_2_^-^. Panel D: Calculated O_2_ consumption- and N_2_ production rates. Inserted panels: exponential regression of initial rates of O_2_-reduction (oxic phase). Panel E: Calculated the rates of electron flow channeled to O_2_ (O2 reduction), and denitrification (Nox Eflow), and the sum (Total Eflow). The electron flow rate to denitrification was calculated from rates of NO_3_^-^-reduction to NO_2_^-^ (2 mol electrons per mol N), and the three subsequent reduction steps, NO_2_^-^→NO→N_2_O→N_2_ (1 mol electron per mol N for each reaction), as derived from measurements. Error bars: standard deviation (n = 3).

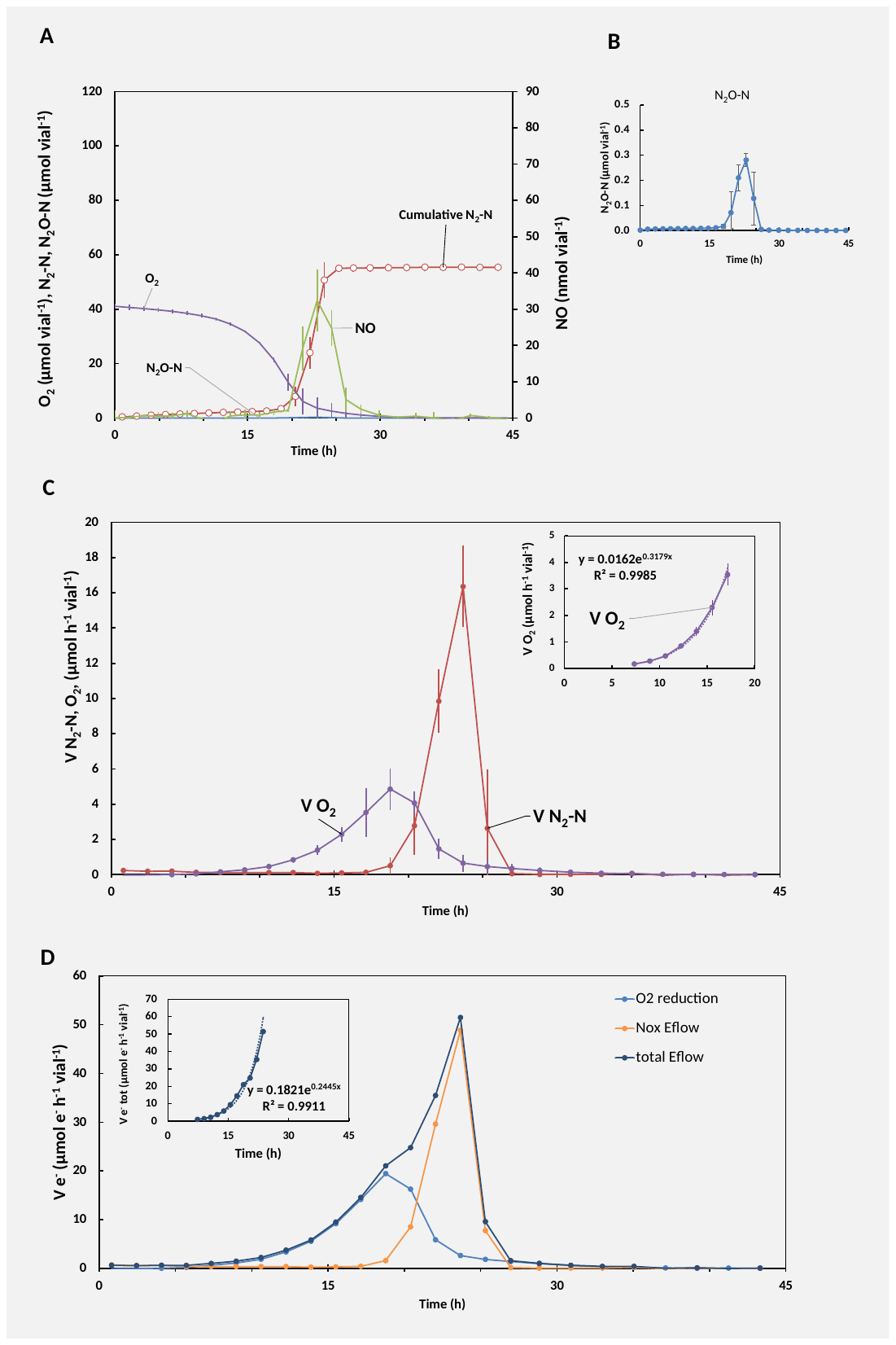

**Figure S18: Denitrification phenotype of *Pseudomonas* sp. PS-02 when provided with O_2_ and NO_2_.** PS-02 was grown in gas-tight 120 mL vials with 50 mL Sistrom’s succinate medium initially supplemented with 1 mL O_2_ and 1mM NO_2_^-^ at constant temperature and stirring (20 °C, 600 rpm). Initial OD_660_ ≈ 0.001. Panel A**:** Measured gases (N_2_O-N, NO, O_2_) and calculated cumulative N_2_-N throughout the incubation. Panel B: Measured N_2_O-N throughout the incubation. Panel C: Calculated O_2_ and N_2_O-N consumption- and N_2_-N production rates. Inserted panels: exponential regression of initial rates of O_2_-reduction (oxic phase). Panel E: Calculated electron flow rates of total electrons channeled to O_2_ (O2 reduction), the NO_x_ reductases (Nox Eflow), and the sum of the total electron flow to terminal oxidases (Total Eflow). Inserted panel: exponential regression of total electron flow from oxic to anoxic phase. Error bars displayed as standard deviation (n = 3).

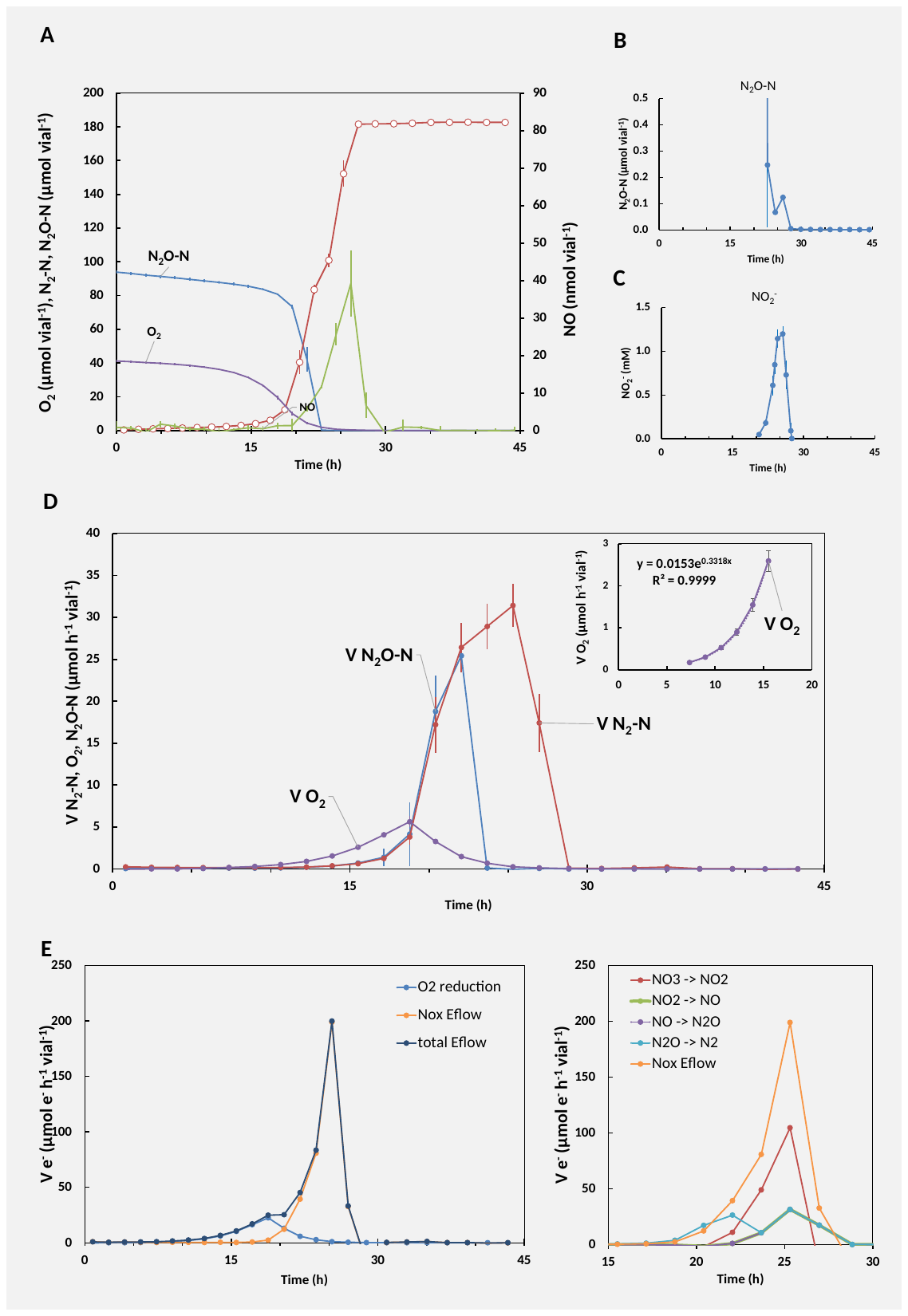

**Figure S19: Denitrification phenotype of *Pseudomonas* sp. PS-02 when provided with O_2_, N_2_O and NO_3_.** PS-02 was grown in gas-tight 120 mL vials with 50 mL Sistrom’s succinate medium initially supplemented with 1 mL O_2_, 1 mL N_2_O, and 2mM NO_3_^-^ at constant temperature and stirring (20 °C, 600 rpm). Initial OD_660_ ≈ 0.001. Panel A**:** Measured gases (N_2_O-N, NO, O_2_) and calculated cumulative N_2_-N throughout the incubation. Panel B: Measured N_2_O-N throughout the incubation. Panel C: Measured liquid concentration of NO_2_^-^. Panel D: Calculated O_2_ and N_2_O-N consumption- and N_2_-N production rates. Inserted panels: exponential regression of initial rates of O_2_-reduction (oxic phase). Error bars displayed as standard deviation (n = 3). Panels E; Left panel: Calculated electron flow rates of total electrons channeled to O_2_ (O2 reduction), the NO_x_ reductases (Nox Eflow), and the sum of the total electron flow to terminal oxidases (Total Eflow). Inserted panel: exponential regression of total electron flow from oxic to anoxic phase. Right panel: Calculated electron flow rates of total electrons channeled to Nar, Nir, Nor, and Nos and summed electron transfer to the NO_x_ reductases (Nox Eflow).

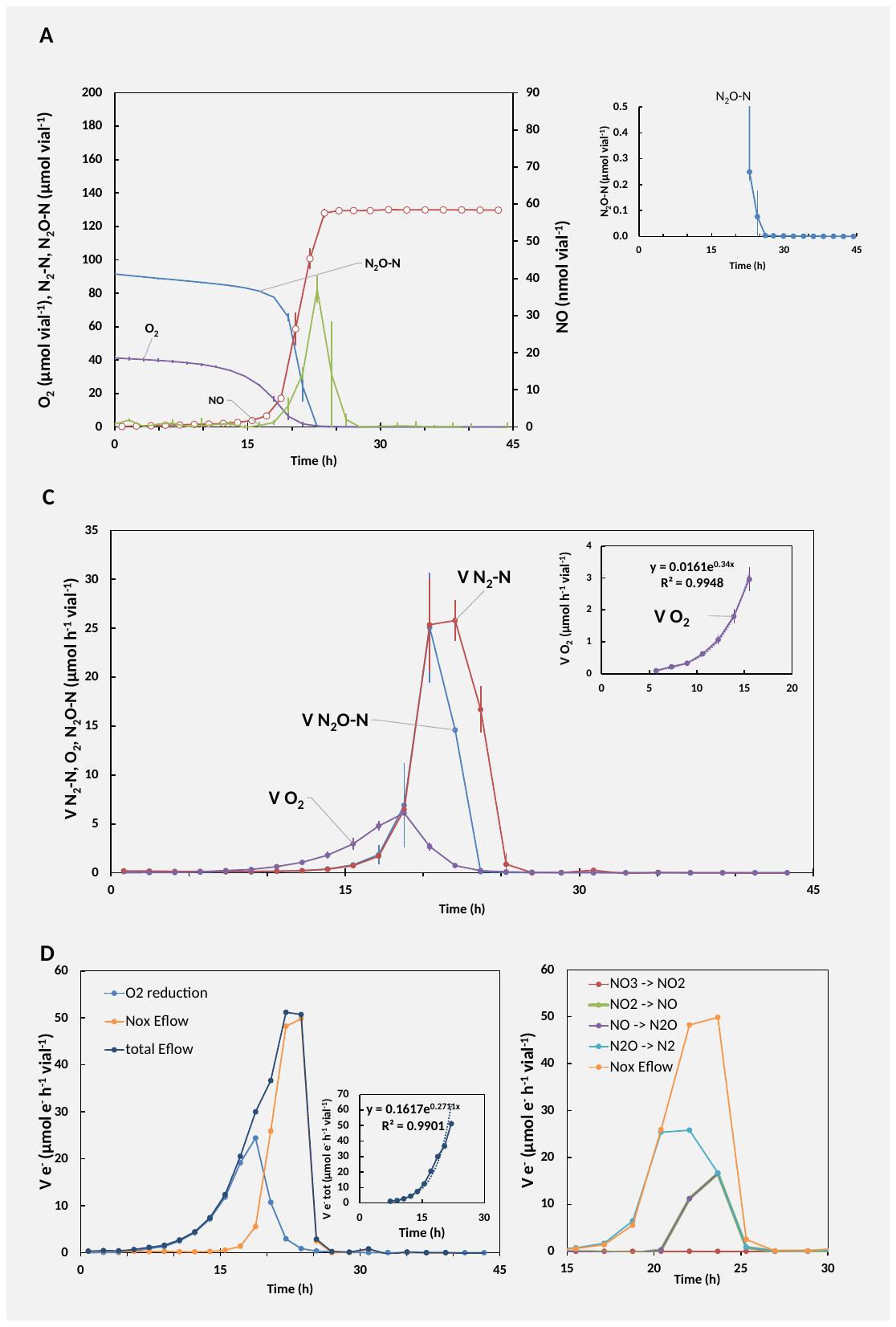

**Figure S20: Denitrification phenotype of *Pseudomonas* sp. PS-02 when provided with O_2_, N_2_O and NO_2_.** PS-02 was grown in gas-tight 120 mL vials with 50 mL Sistrom’s succinate medium initially supplemented with 1 mL O_2_, and 1 mL N_2_O and 1mM NO_2_^-^ at constant temperature and stirring (20 °C, 600 rpm). Initial OD_660_ ≈ 0.001. Panel A**:** Measured gases (N_2_O-N, NO, O_2_) and calculated cumulative N_2_-N throughout the incubation. Panel B: Measured N_2_O-N throughout the incubation. Panel C: Calculated O_2_ and N_2_O-N consumption- and N_2_-N production rates. Inserted panels: exponential regression of initial rates of O_2_-reduction (oxic phase). Error bars displayed as standard deviation (n = 3). Panels D; Left panel: Calculated electron flow rates of total electrons channeled to O_2_ (O2 reduction), the NO_x_ reductases (Nox Eflow), and the sum of the total electron flow to terminal oxidases (Total Eflow). Inserted panel: exponential regression of total electron flow from oxic to anoxic phase. Right panel: Calculated electron flow rates of total electrons channeled to Nar, Nir, Nor, and Nos and summed electron transfer to the NO_x_ reductases (Nox Eflow).

#### 6.2 *Ochrobactrum* sp. OB

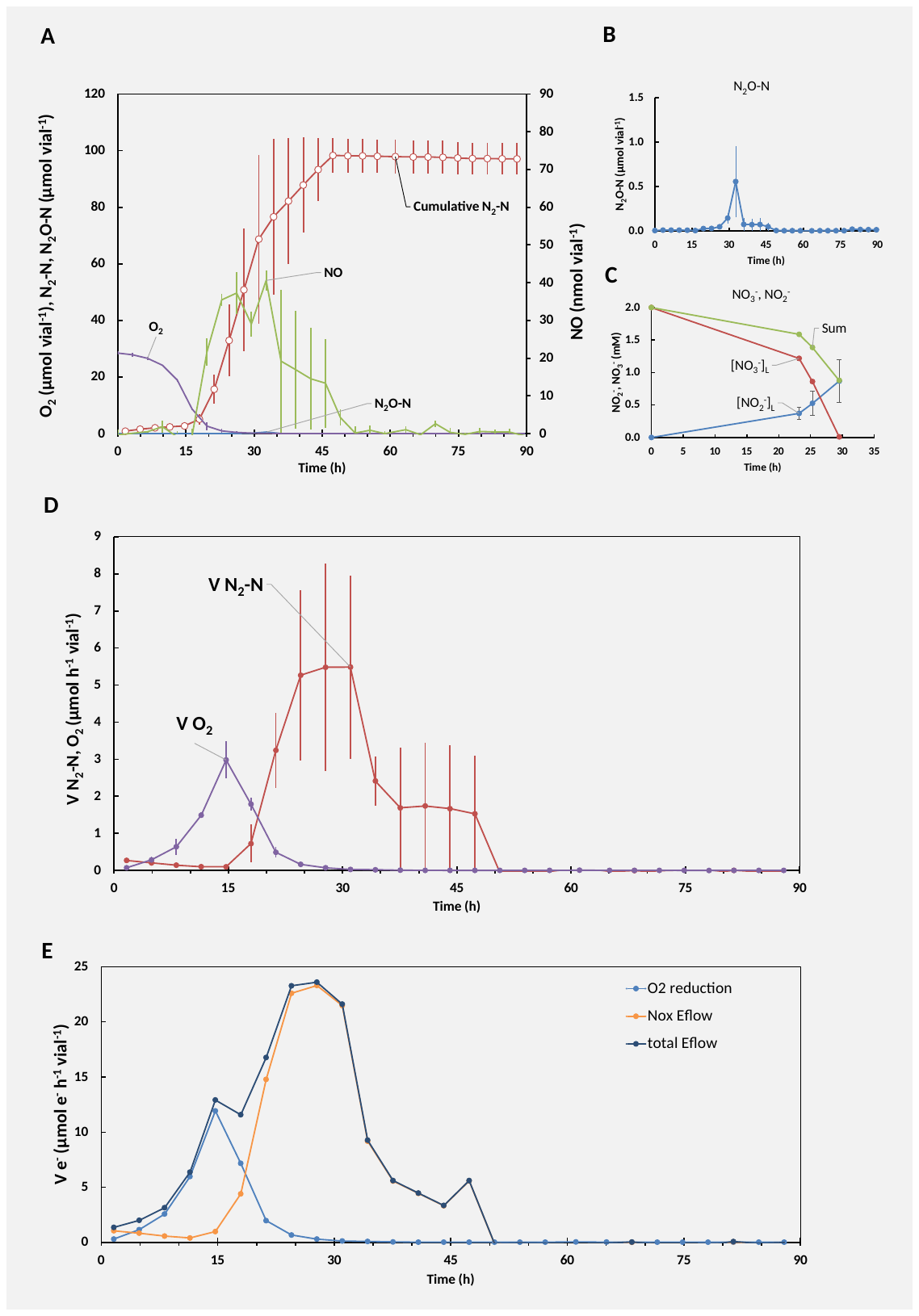
*Ochrobactrum* sp. OB (**Fig. 4C** in main paper) had the genetic capacity of a full-fledged denitrifier carrying cytoplasmic *nar*, periplasmic *nap*, *nirK*, *nor* and *nosZ* (clade I), thus predicting a full-fledged denitrifier, which was also reflected by the isolate’s denitrifying phenotype: it transiently accumulated nitrite, whilst keeping the gaseous intermediates NO and N_2_O low throughout our incubations. A marginal dip in electron flow in the transition between oxic and anoxic respiration of NO_3_^-^ (**Fig. S20**) or NO_2_^-^ (**Fig. S21**) can be understood as a fraction of the respiring cells that did not fully commit to denitrification by not expressing Nir. However, the isolate also demonstrated non-exponential growth (linear increase in N_2_ production rates) when respiring solely on NO_2_^-^, indicating restricted growth. In incubations supplemented with exogenous N_2_O in addition to NO_3_^-^ (**Fig. 4C** in main paper, additional gas kinetics in **Fig. S22**) or NO_2_^-^ (**Fig. S23**) gas kinetics indicated that the reduction of N_2_O was preferred over nitrate and nitrate. This sums up to make *Ochrobactrum* sp. OB a potential N_2_O sink as N_2_O reduction was favored over NO_3_^-^ and NO_2_^-^ while limiting the depletion of oxyanions at higher NO_2_^-^ concentrations.

**Figure S21 Denitrification phenotype of *Ochrobactrum* sp. OB when provided with O_2_ and NO_3_^-^.** OB was grown in gas-tight 120 mL vials with 50 mL Sistrom’s succinate medium initially supplemented with 1 mL O_2_ and 2mM NO_3_^-^ at constant temperature and stirring (20 °C, 600 rpm). Initial OD_660_ ≈ 0.001. Panel A**:** Measured gases (N_2_O-N, NO, O_2_) and calculated cumulative N_2_-N throughout the incubation. Panel B: Measured N_2_O-N throughout the incubation. Panel C: Measured liquid concentration of NO_3_^-^ and NO_2_^-^ and the sum of NO_3_^-^ and NO_2_^-^. Panel D: Calculated O_2_ and N_2_O-N consumption- and N_2_-N production rates. Inserted panels: exponential regression of initial rates of O_2_-reduction (oxic phase). Panel E: Calculated electron flow rates of total electrons channeled to O_2_ (O2 reduction), the NOx reductases (Nox Eflow), and the sum of the total electron flow to terminal oxidases (Total Eflow). Error bars displayed as standard deviation (n = 2).

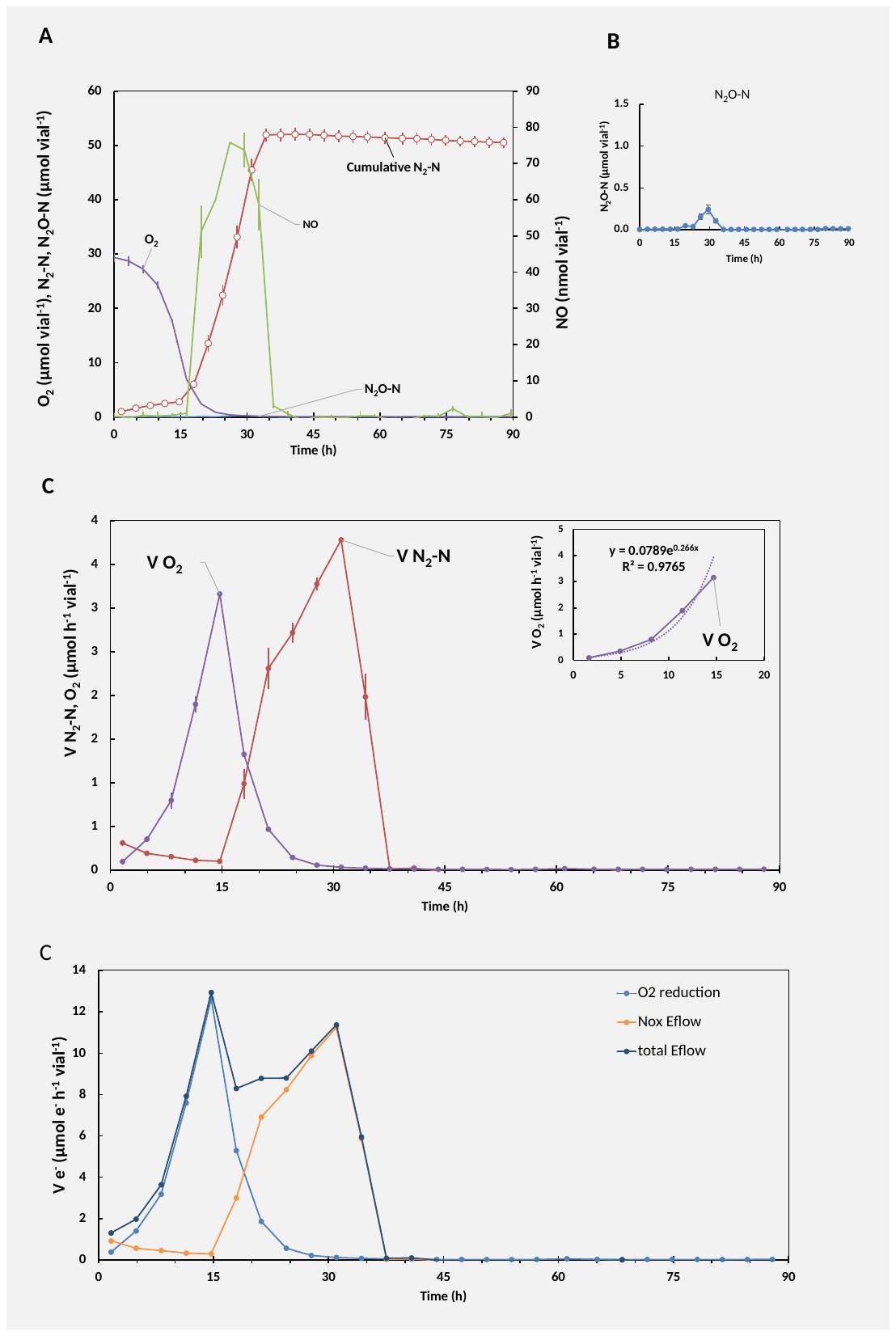

**Figure S22: Denitrification phenotype of *Ochrobactrum* sp. OB when provided with O_2_ and NO_2_.** OB was grown in gas-tight 120 mL vials with 50 mL Sistrom’s succinate medium initially supplemented with 1 mL O_2_ and 1mM NO_2_^-^ at constant temperature and stirring (20 °C, 600 rpm). Initial OD_660_ ≈ 0.001. Panel A**:** Measured gases (N_2_O-N, NO, O_2_) and calculated cumulative N_2_-N throughout the incubation. Panel B: Measured N_2_O-N throughout the incubation. Panel C: Calculated O_2_ and N_2_O-N consumption- and N_2_-N production rates. Inserted panels: exponential regression of initial rates of O_2_-reduction (oxic phase) and N_2_-N production rates. Panel E: Calculated electron flow rates of total electrons channeled to O_2_ (O2 reduction), the NOx reductases (Nox Eflow), and the sum of the total electron flow to terminal oxidases (Total Eflow). Error bars displayed as standard deviation (n = 2).

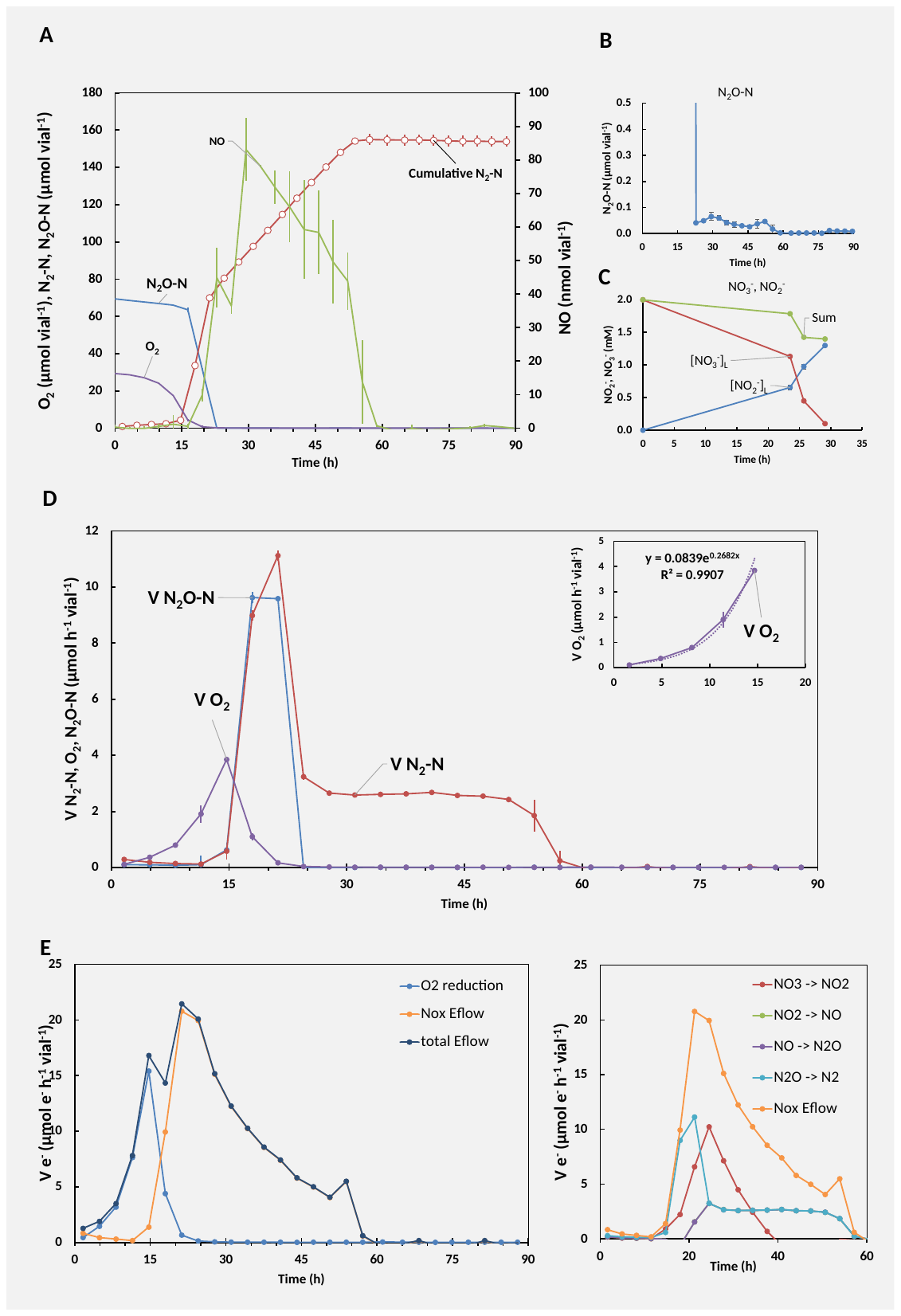

**Figure S23: Denitrification phenotype of *Ochrobactrum* sp. OB when provided with O_2_, N_2_O and NO_3_.** OB was grown in gas-tight 120 mL vials with 50 mL Sistrom’s succinate medium initially supplemented with 1 mL O_2_, 1 mL N_2_O and 2mM NO_3_^-^ at constant temperature and stirring (20 °C, 600 rpm). Initial OD_660_ ≈ 0.001. Panel A**:** Measured gases (N_2_O-N, NO, O_2_) and calculated cumulative N_2_-N throughout the incubation. Panel B: Measured N_2_O-N throughout the incubation. Panel C: Measured liquid concentration of NO_3_^-^ and NO_2_^-^ and the sum of NO_3_^-^ and NO_2_^-^. Panel D: Calculated O_2_ and N_2_O-N consumption- and N_2_-N production rates. Inserted panels: exponential regression of initial rates of O_2_-reduction (oxic phase). Error bars displayed as standard deviation (n = 2). Panel E: Left panel: Calculated electron flow rates of total electrons channeled to O_2_ (O2 reduction), the NO_x_ reductases (Nox Eflow), and the sum of the total electron flow to terminal oxidases (Total Eflow). Inserted panel: exponential regression of total electron flow from oxic to anoxic phase. Right panel: Calculated electron flow rates of total electrons channeled to Nar, Nir, Nor and Nos and summed electron transfer to the NO_x_ reductases (Nox Eflow).

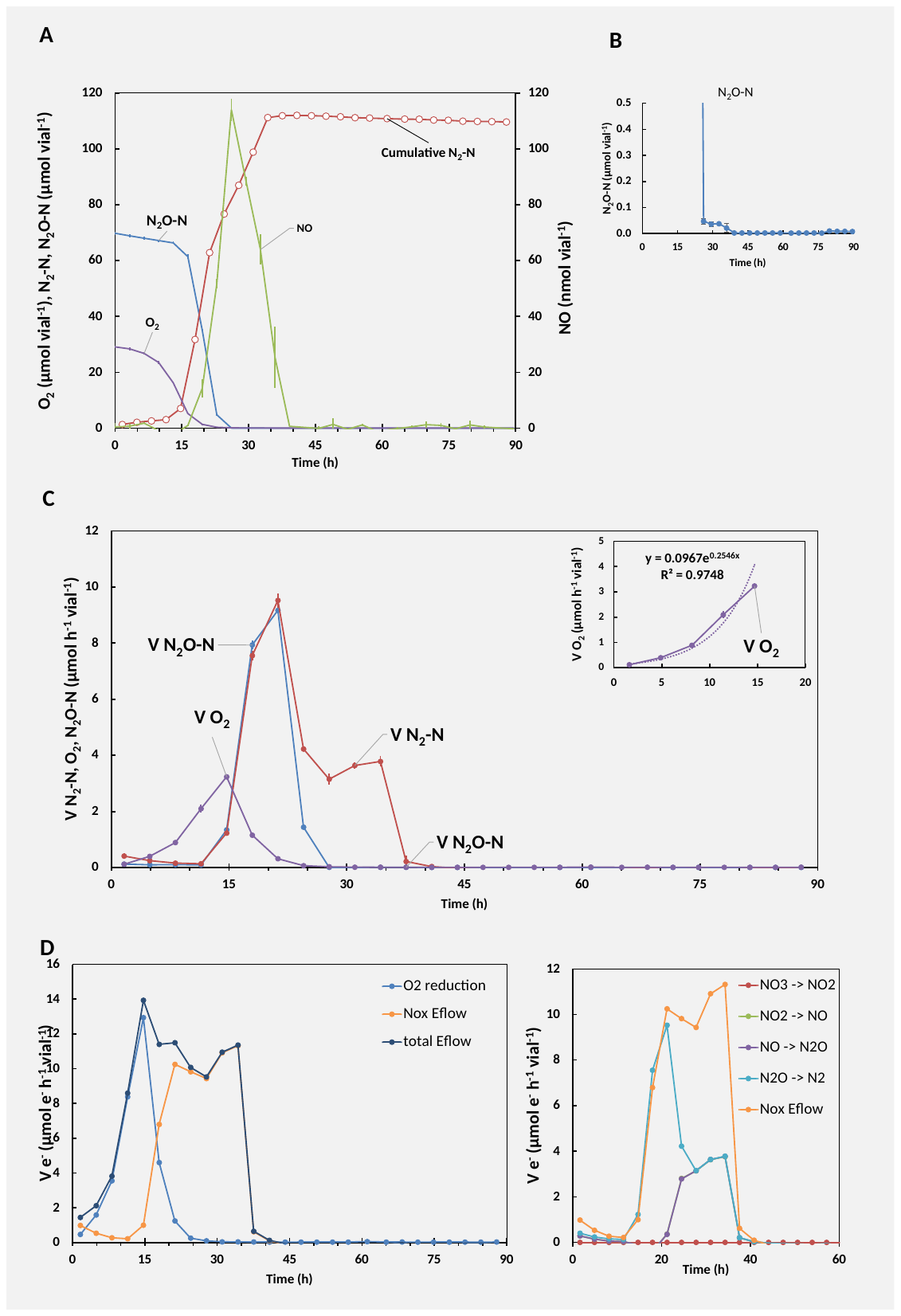

**Figure S24: Denitrification phenotype of *Ochrobactrum* sp. OB when provided with O_2_, N_2_O and NO_2_.** OB was grown in gas-tight 120 mL vials with 50 mL Sistrom’s succinate medium initially supplemented with 1 mL O_2_, 1 mL N_2_O and 1mM NO_2_^-^ at constant temperature and stirring (20 °C, 600 rpm). Initial OD_660_ ≈ 0.001. Panel A**:** Measured gases (N_2_O-N, NO, O_2_) and calculated cumulative N_2_-N throughout the incubation. Panel B: Zoom-in on measured N_2_O-N throughout the incubation. Panel C: Calculated O_2_ and N_2_O-N consumption- and N_2_-N production rates. Inserted panels: exponential regression of initial rates of O_2_-reduction (oxic phase). Error bars displayed as standard deviation (n = 2). Panel E: Left panel: Calculated electron flow rates of total electrons channeled to O_2_ (O2 reduction), the NO_x_ reductases (Nox Eflow), and the sum of the total electron flow to terminal oxidases (Total Eflow). Inserted panel: exponential regression of total electron flow from oxic to anoxic phase. Right panel: Calculated electron flow rates of total electrons channeled to Nar, Nir, Nor, and Nos and summed electron transfer to the NO_x_ reductases (Nox Eflow).

#### 6.3 *Brachymonas sp*. BM

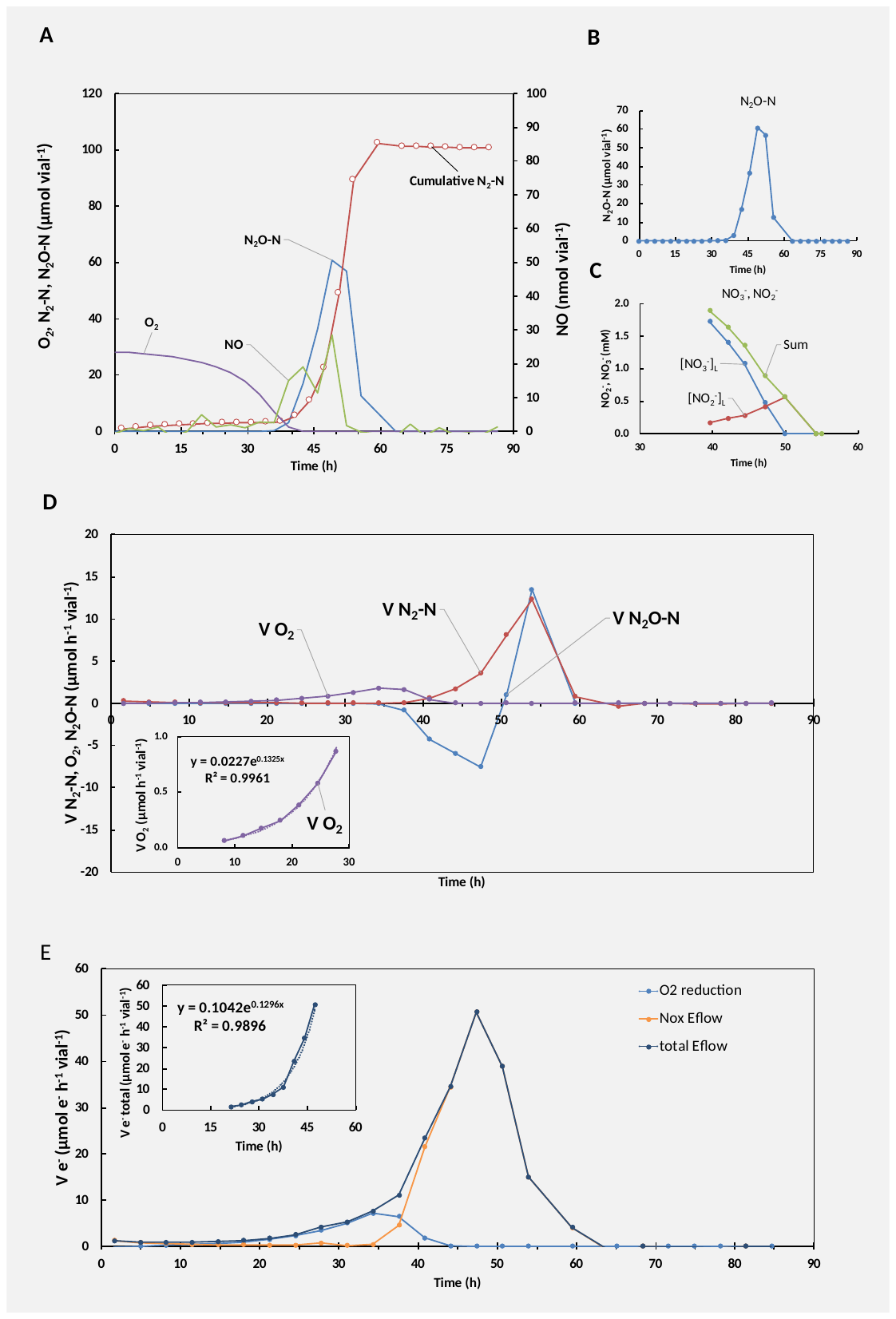
 *Brachymonas* sp. BM demonstrated phenotypic characteristics of a full-fledged denitrifier, but, in contrast to PS-02, BM accumulated significant levels of N_2_O as a response to transitioning from oxic to anoxic conditions. When respiring NO_3_^-^ a continuous electron flow to the terminal oxidoreductases to terminal nitrogen reductases implied that most cells committed to denitrification when transitioning from oxic to anoxic conditions. NO_2_^-^ accumulated to lower levels than the two other full-fledged denitrifying organisms that were isolated (PS-02 and OB) (**Fig. S25**). Parallel incubations supplemented with 1 mL O_2_, 1 mL N_2_O and 2 mM NO_3_^-^ (**Fig. S26,** and **Fig. 4D** in main paper) or 1 mM NO_2_^-^ (**Fig. S27**) demonstrated similar N_2_O accumulation throughout the isolate’s depletion of NO_x_. Thus, BM did not prefer N_2_O reduction over NO_3_^-^ and could be predicted as a net N_2_O source.

**Figure S25: Denitrification phenotype of *Brachymonas* sp. BM when provided with O_2_ and NO_3_.** BM was grown in gas-tight 120 mL vials with 50 mL Sistrom’s succinate medium initially supplemented with 1 mL O_2_ and 2mM NO_3_^-^ at constant temperature and stirring (20 °C, 600 rpm). Initial OD_660_ ≈ 0.001. Panel A**:** Measured gases (N_2_O-N, NO, O_2_) and calculated cumulative N_2_-N throughout the incubation. Panel B: Measured N_2_O-N throughout the incubation. Panel C: Measured liquid concentration of NO_3_^-^ and NO_2_^-^ and the sum of NO_3_^-^ and NO_2_^-^. Panel D: Calculated O_2_ and N_2_O-N consumption- and N_2_-N production rates. Inserted panels: exponential regression of initial rates of O_2_-reduction (oxic phase). Panel E: Calculated electron flow rates of total electrons channeled to O_2_ (O2 reduction), the NOx reductases (Nox Eflow), and the sum of the total electron flow to terminal oxidases (Total Eflow). Inserted panel: exponential regression of total electron flow through transition from oxic to anoxic phase. Results from a single vial shown. Replicate vials showed similar gas kinetics.

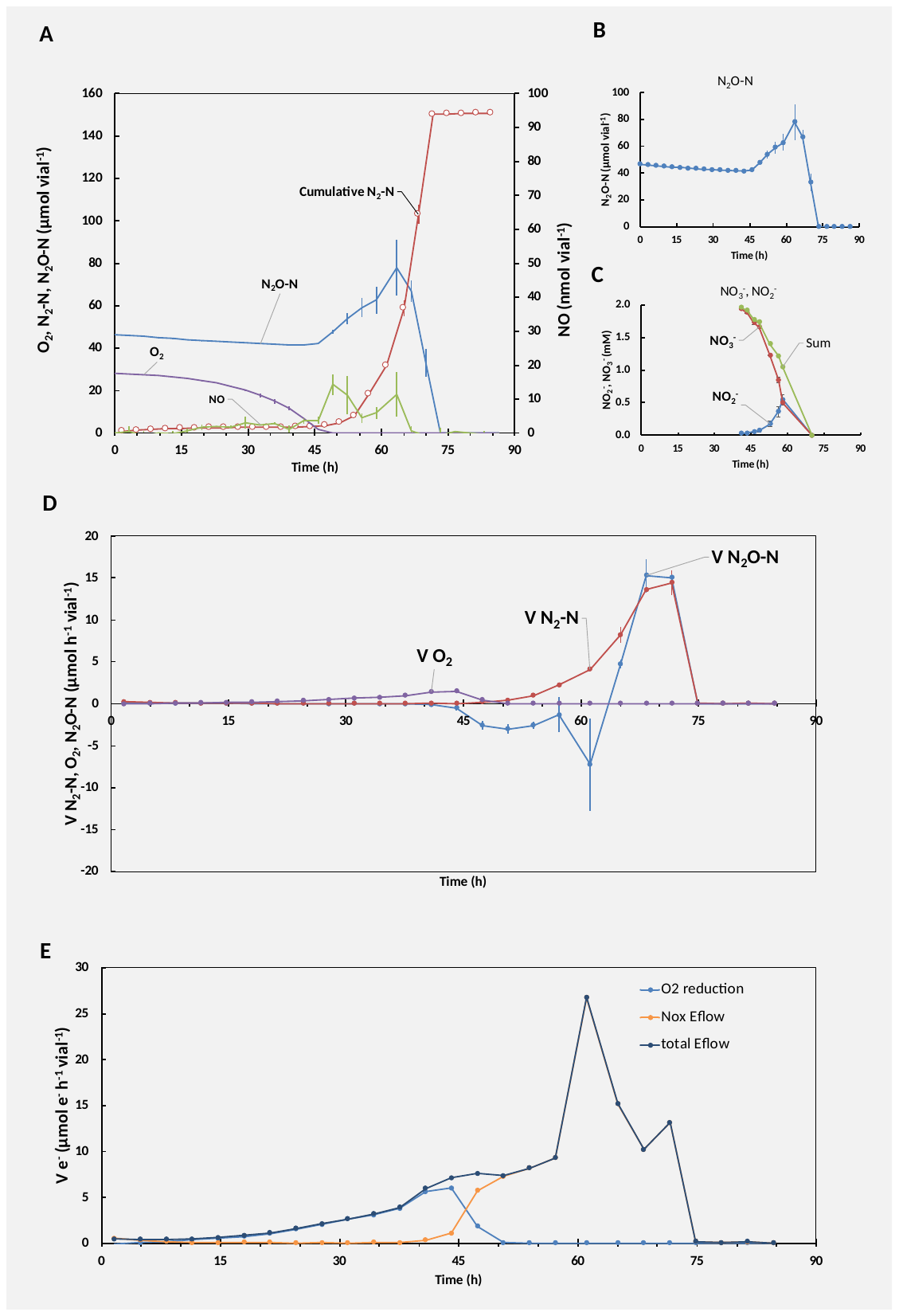

**Figure S26: Denitrification phenotype of *Brachymonas* sp. BM when provided with O_2_, N_2_O and NO_3_.** BM was grown in gas-tight 120 mL vials with 50 mL Sistrom’s succinate medium initially supplemented with 1 mL O_2_, 1 mL N_2_O and 2mM NO_3_^-^ at constant temperature and stirring (20 °C, 600 rpm). Initial OD_660_ ≈ 0.001. Panel A**:** Measured gases (N_2_O-N, NO, O_2_) and calculated cumulative N_2_-N throughout the incubation. Panel B: Measured N_2_O-N throughout the incubation. Panel C: Measured liquid concentration of NO_3_^-^ and NO_2_^-^ and the sum of NO_3_^-^ and NO_2_^-^. Panel D: Calculated O_2_ and N_2_O-N consumption- and N_2_-N production rates. Inserted panels: exponential regression of initial rates of O_2_-reduction (oxic phase) and N_2_-N production. Panel E: Calculated electron flow rates of total electrons channeled to O2 (O2 reduction), the NOx reductases (Nox Eflow), and the sum of the total electron flow to terminal oxidases (Total Eflow). Error bars displayed as standard deviation (n = 2).

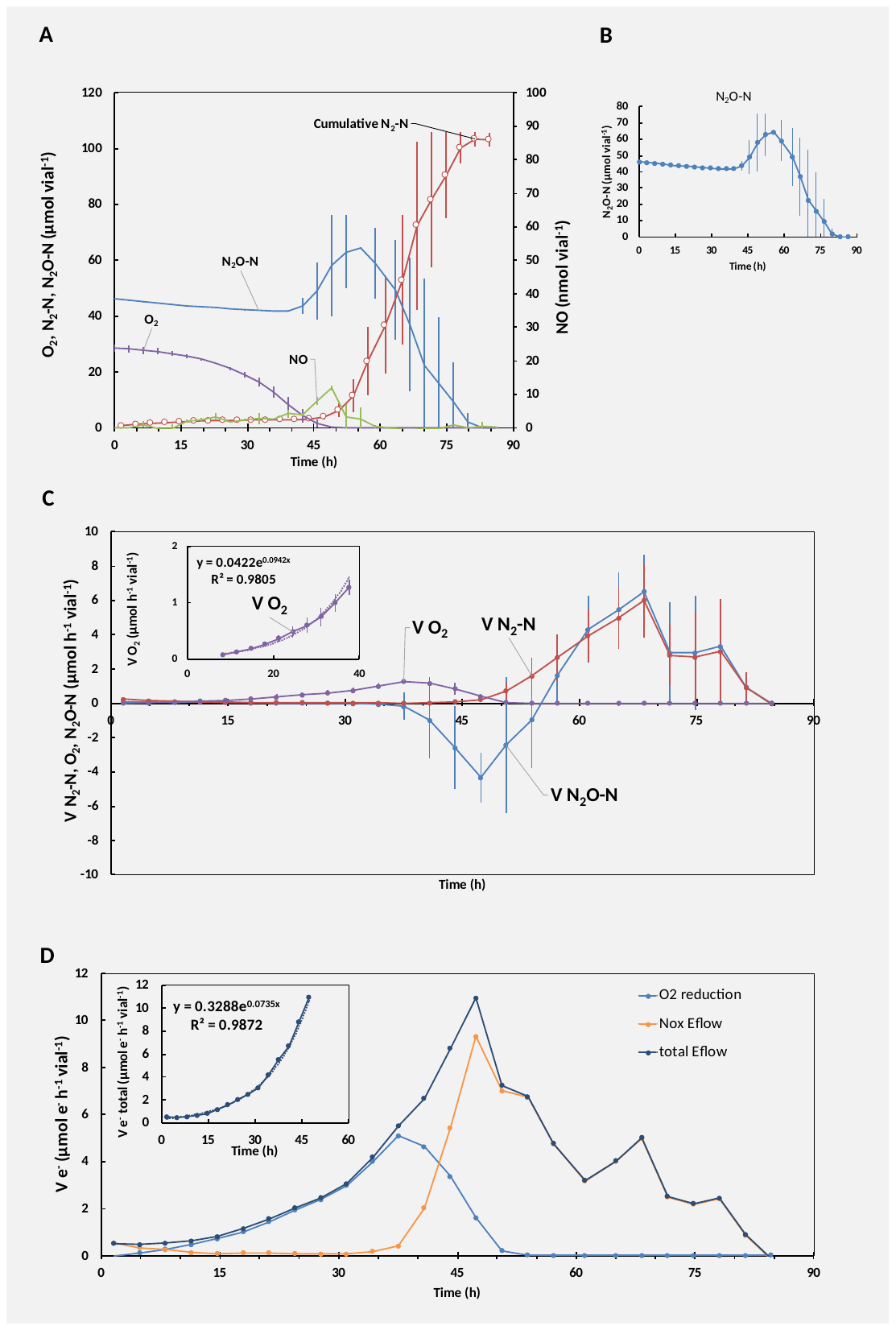

**Figure S27: Denitrification phenotype of *Brachymonas* sp. BM when provided with O_2_, N_2_O and NO_2_.** BM was grown in gas-tight 120 mL vials with 50 mL Sistrom’s succinate medium initially supplemented with 1 mL O_2_, 1 mL N_2_O and 1mM NO_2_^-^ at constant temperature and stirring (20 °C, 600 rpm). Initial OD_660_ ≈ 0.001. Panel A**:** Measured gases (N_2_O-N, NO, O_2_) and calculated cumulative N_2_-N throughout the incubation. Panel B: Measured N_2_O-N throughout the incubation. Panel C: Calculated O_2_ and N_2_O-N consumption- and N_2_-N production rates. Inserted panels: exponential regression of initial rates of O_2_-reduction (oxic phase) and N_2_-N production. Panel D: Calculated electron flow rates of total electrons channeled to O2 (O2 reduction), the NOx reductases (Nox Eflow), and the sum of the total electron flow to terminal oxidases (Total Eflow). Inserted panel: exponential regression of total electron flow through transition from oxic to anoxic phase Error bars displayed as standard deviation (n = 2).

#### 6.4 *Aeromonas* sp*.* AM

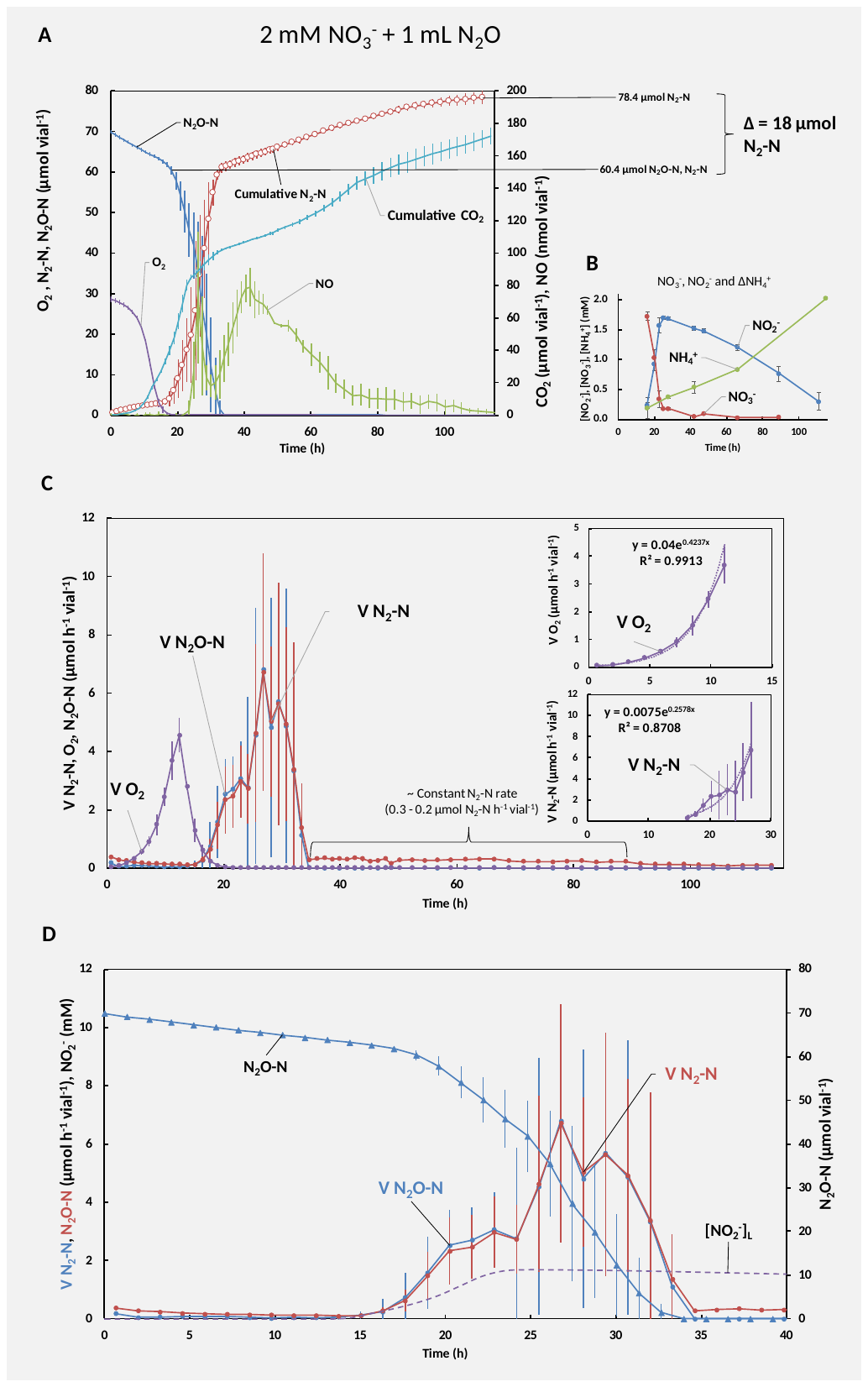
In incubations supplemented with NO_3_^-^ and N_2_O *Aeromonas sp*. AM reduced the available NO_3_^-^ to NO_2_^-^ and N_2_O to N_2_ when transcending from oxic to anoxic conditions (**Fig. S28**, and **Fig. 4B** in main paper). The accumulated NO_2_^-^ was slowly reduced to NH_4_^+^ throughout the incubation, indicative of dissimilatory nitrate reduction to ammonium (DNRA). Following N_2_O depletion AM continued channeling electrons toward denitrification and N_2_-N production indicating denitrification alongside DNRA activity. The phenotype was corroborated by annotation of the *nrfA* gene, coding for a key enzyme of DNRA (Cytochrome c552 nitrite reductase, EC: 1.7.2.2), and denitrification genes coding for periplasmic nitrate reductase (*napAB*) and N_2_O reductase (*nosZ*, clade I) in AN’s genome (**Fig. 4B** in main paper). N_2_O was reduced alongside NO_3_^-^, which would contradict the hypothesis that Nos outcompetes Nap for electrons (Mania et al 2020), however, a nitrate reductase (NasA) was recovered in the genome. The genome also encoded the gene NasD, of which gene product shared high sequence identity (protein blast) with a NADH dependent nitrite reductase of Aeromonas media strain (see main text for details).

**Figure S28: DNRA and denitrifying phenotype of *Aeromonas* sp. AM when provided with N_2_O and NO_3_.** AM was grown in gas-tight 120 mL vials with 50 mL Sistrom’s succinate medium initially supplemented with 1 mL O_2_, 1 mL N_2_O, and 2mM NO_3_^-^ at constant temperature and stirring (20 °C, 700 rpm). Initial OD_660_ ≈ 0.001. Panel A**:** Measured gases (N_2_O-N, NO, O_2_) and calculated cumulative N_2_-N and CO_2_ throughout the incubation. Panel B: Measured liquid concentration of NO_3_^-^ NO_2_^-^ and ΔNH_4_^+^. Panel C: Calculated O_2_ and N_2_O-N consumption- and N_2_-N production rates. Inserted panels: exponential regression of initial rates of O_2_-reduction (oxic phase) and N_2_-N production rates. Panel D: N_2_O-N consumption- and N_2_-N production rates, measured N_2_O-N, and estimated NO_2_^-^ concentration (mM) (estimated based on measurements in Panel B with the Excel Spline function) throughout the first 40 hours of incubation. Error bars displayed as standard deviation (n = 3).

#### 6.5 *Cloacibacterium sp.* CB-01 and CB-03

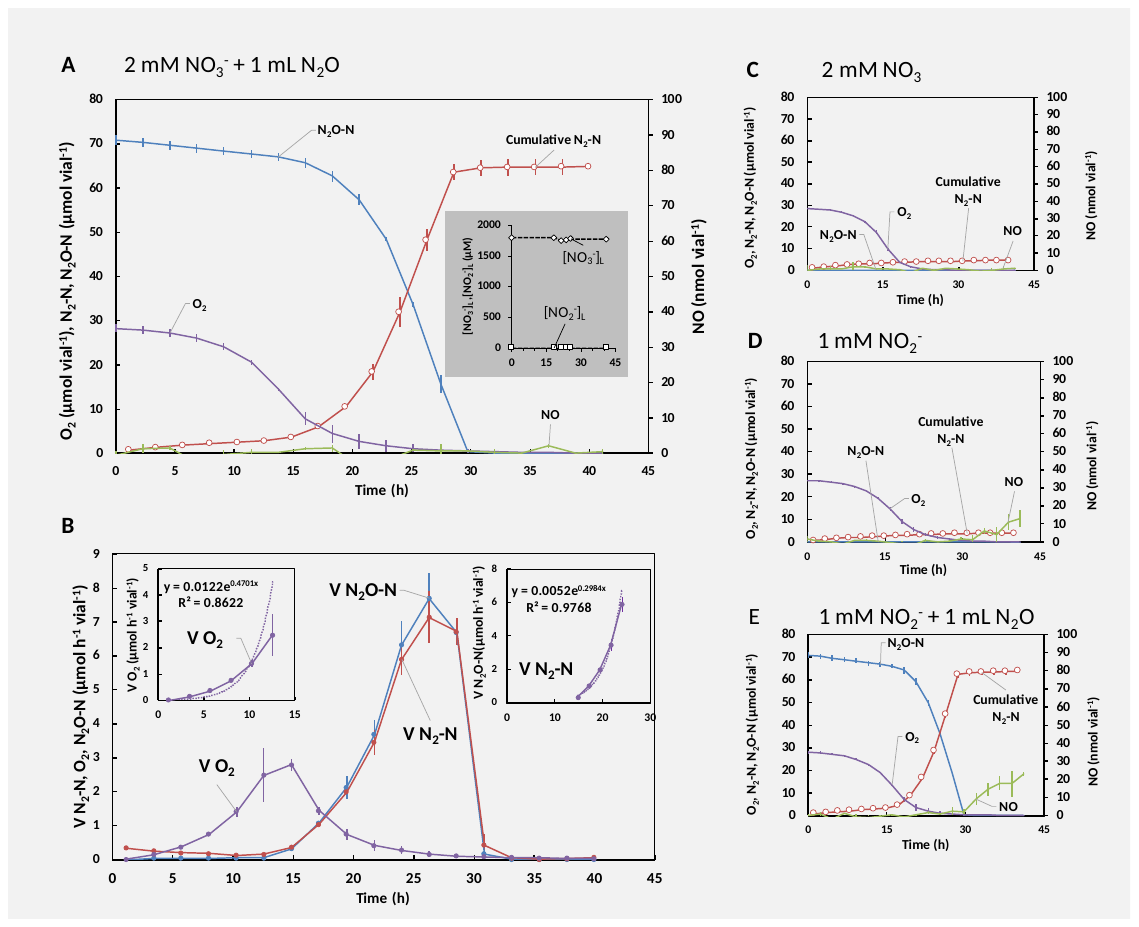

**Figure S29: Denitrification phenotype of *Cloacibacterium* *sp*. CB-01 when provided with O_2_ and combinations of NO_3_, NO_2_ and/or N_2_O.** CB-01 was grown in gas-tight 120 mL vials with 50 mL Nutrient Broth medium initially supplemented with 1 mL O_2_ and combinations of 1 mL N_2_O and/or 2mM NO_3_^-^ and/or 1 mM NO_2_^-^ at constant temperature and stirring (20 °C, 600 rpm). Initial OD_660_ ≈ 0.001. **Panel A:** Measured gases (N_2_O-N, NO, O_2_) and calculated cumulative N_2_-N throughout an incubation initially supplemented with 1 mL O_2_, 2 mM NO_3_^-^ and 1 mL N_2_O. Inserted panel shows measures liquid concentration of NO_3_^-^ and NO_2_^-^. **Panel B**: Calculated O_2_ and N_2_O-N consumption- and N_2_-N production rates. Inserted panels: exponential regression of initial rates of O_2_-reduction (oxic phase) and N_2_-N production rates. **Panel C, D and E**: Measured gases (N_2_O-N, NO, O_2_) and calculated cumulative N_2_-N throughout incubations initially supplemented with 1 mL O_2_ and 2 mM NO_3_^-^, 1 mL O_2_ and 1 mM NO_2_^-^ and 1 mL O_2_, 1 mM NO_2_^-^ and 1 mL N_2_O, respectively. Error bars displayed as standard deviation (n = 2). Error bars for inserted panel in Panel A (measured NO_2_^-^ and NO_3_^-^) displayed as standard deviation (n = 3).

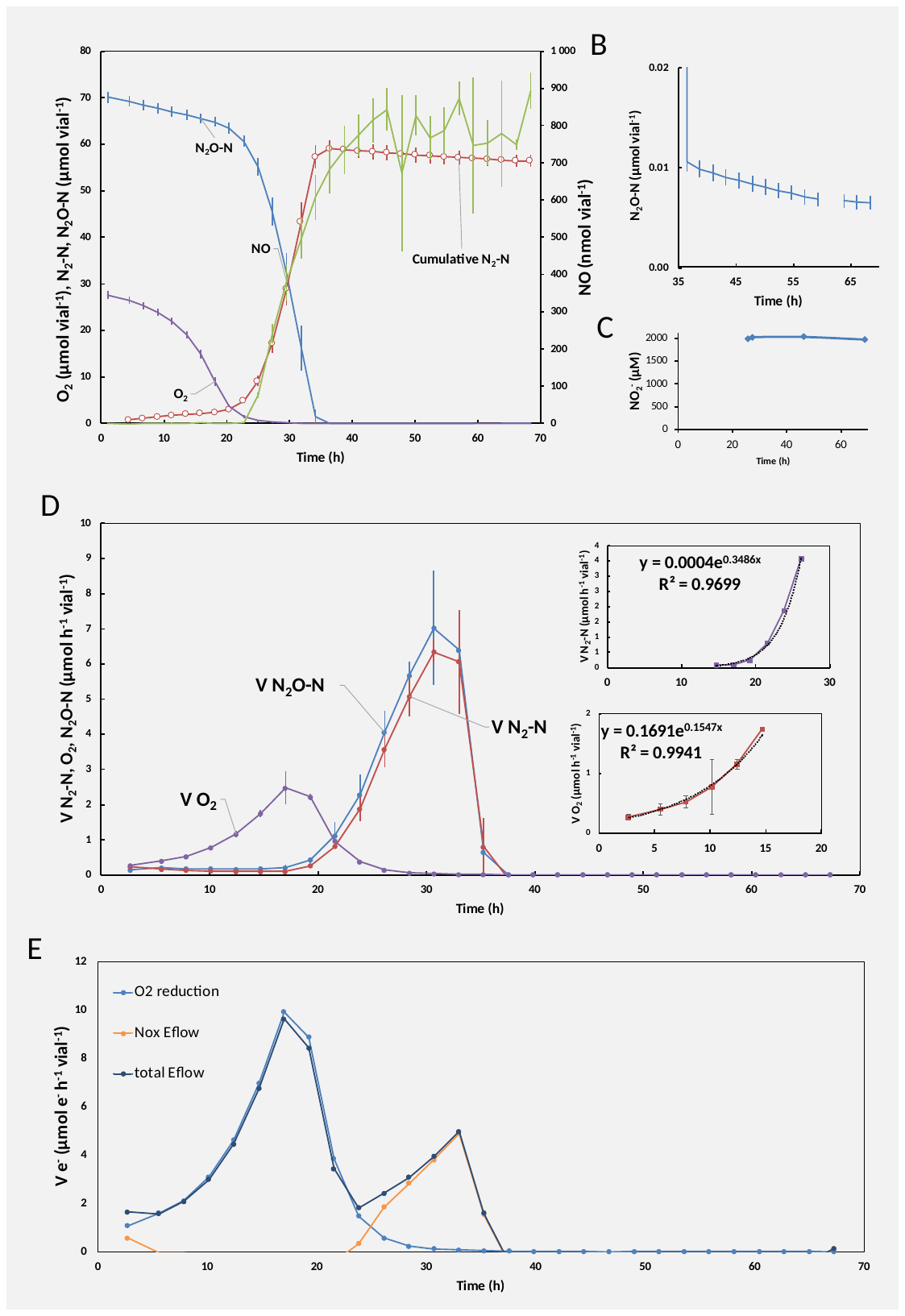

**Figure S30: Denitrification phenotype of *Cloacibacterium* sp. CB-03 when provided with O_2_, N_2_O and NO_3_.** CB-03 was grown in gas-tight 120 mL vials with 50 mL NB medium initially supplemented with 1 mL O_2_, 1 mL N_2_O and 2mM NO_3_^-^ at constant temperature and stirring (20 °C, 600 rpm). Initial OD_660_ ≈ 0.001. Panel A**:** Measured gases (N_2_O-N, NO, O_2_) and calculated cumulative N_2_-N throughout the incubation. Panel B: Zoom in on measured N_2_O-N from t = 35 h and throughout. Panel C: Measured liquid concentration of NO_2_^-^ (µM). Panel D: Calculated O_2_ and N_2_O-N consumption- and N_2_-N production rates. Inserted panels: exponential regression of initial rates of O_2_-reduction (oxic phase) and N_2_-N production rates. Panel E: Calculated electron flow rates of total electrons channeled to O_2_ (O_2_ reduction), the NOx reductases (Nox Eflow) and the sum of the total electron flow to terminal oxidases (Total Eflow). Error bars displayed as standard deviation (n = 2).

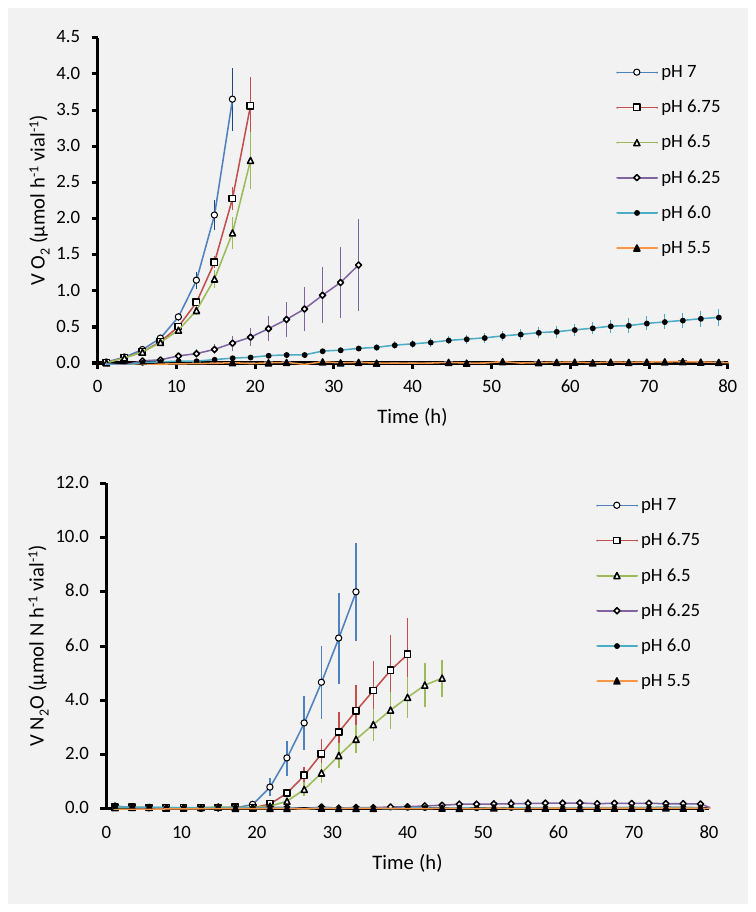

**Figure S31: Denitrification phenotype of *Cloacibacterium* sp. CB-01 when provided with 1 mL O_2_ and 1 mL N_2_O at different pH levels.** CB-01 was grown in gas,tight 120 mL vials with 50 mL NB medium initially supplemented with 1 mL O_2_ and 1 mL N_2_O at constant temperature and stirring (20 °C, 600 rpm). Initial OD_660_ ≈ 0.001. Top panel**:** First period of O_2_ reduction (rate) during incubations with media adjusted to different pH levels. Bottom panel: Panel B: First period of N_2_O reduction (rate) during incubations with media adjusted to different pH. Error bars displayed as standard deviation (n = 3).

### 7 Soil incubations

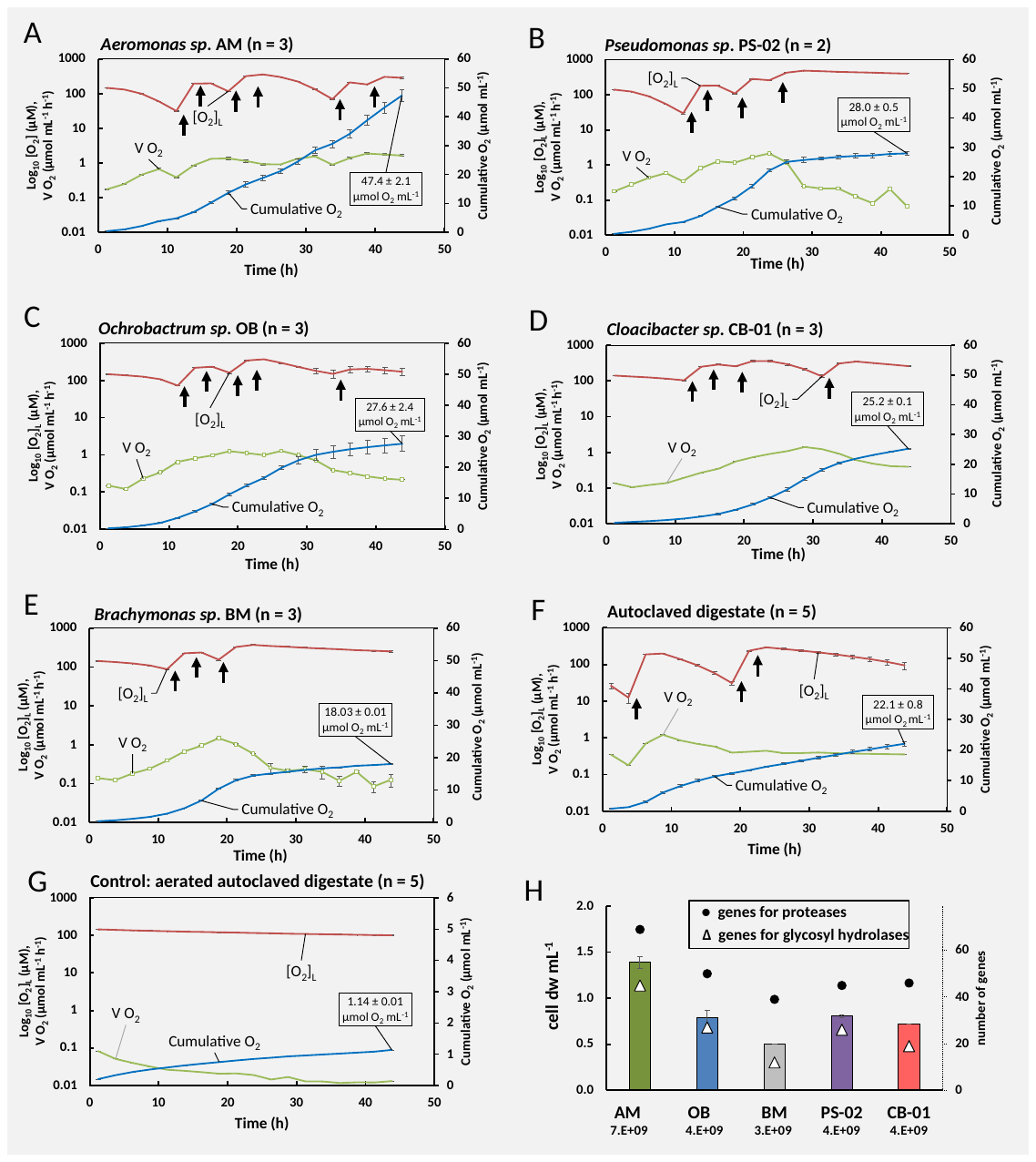

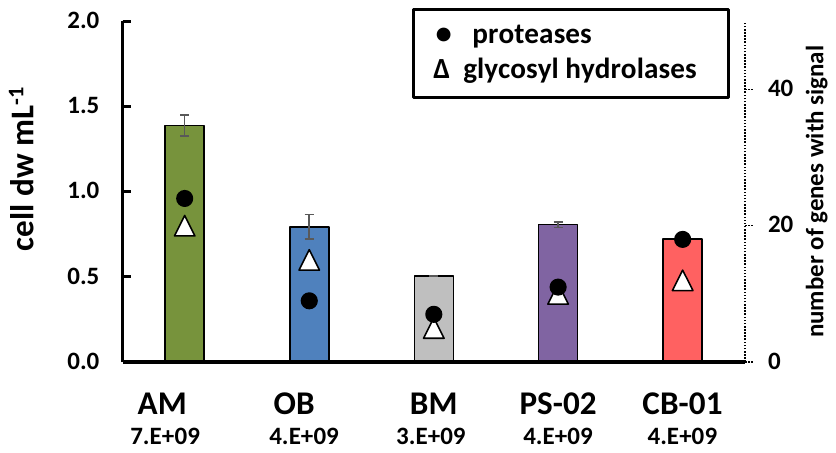

**Figure S32:** **Aerobic growth in autoclaved digestate.** AM, OB, BM, PS-02 and CB-01 were raised aerobically in 50 mL SS (AM, OB, BM and PS-02) or 50 mL NB (CB-01) to high cell densities (OD_660nm_ ~ 1), then transferred (1 mL) to vials with 50 mL stirred (600 rpm) autoclaved pH-adjusted (pH=7.75) and pre-aerated (aerated by pumping sterile filtered air through a stirred suspension for 36 hours) digestate at 20 °C. Oxygen concentration (red), arrows = exogenous O_2_ addition. Cumulative O_2_ reduced = blue. Rate of oxygen consumption = green. Aeration of the autoclaved digestate was necessary to secure near-complete abiotic oxidation of the Fe^2+^ in the digestate, which would otherwise obscure the measurements of O_2_ consumption. **Panel A:** *Aeromonas sp*. AM (n=3). **Panel B:** Pseudomonas sp. PS-02 (n=2). **Panel C:** *Ochrobactrum sp*. OB (n=3). **Panel D:** *Cloacibacterium* sp. CB (n=3). **Panel E:** *Brachymonas sp*. BM (n=3), **Panel** **G:** Non-aerated, pH adjusted (pH = 7.65) autoclaved digestate (n = 5). **Panel F:** Control: Aerated, pH adjusted (pH = 7.75) autoclaved digestate (n = 5). **Panel H**: The cumulated oxygen consumption by each strain was used to estimate the amount of cells produced, assuming that the growth yield for all strains is the same as for *Paracocus denitrificans*, which is 30 g cell dry-weight mol^-1^ O_2_ (based on Bergaust et al (2011): 2·10^-13^ g dry-weight per cell, growth yield = 1.5·10^14^ cells mol^-1^ O_2_). The panel (H) shows estimated amount of cell dry-matter mL^-1^ for each strain as bars (left axis), the number of cells mL^-1^ (below labels). The number of genes coding for glycosyl hydrolases (**GH**) and proteases (**P**) in the genome of each strain (from Table S11 and S13) is shown (right axis, symbols explained in the pale). **P** and **GH** were correlated (r^2^=0.93), and the cell dry-weight was correlated to both (r^2^= 0.97 for both).

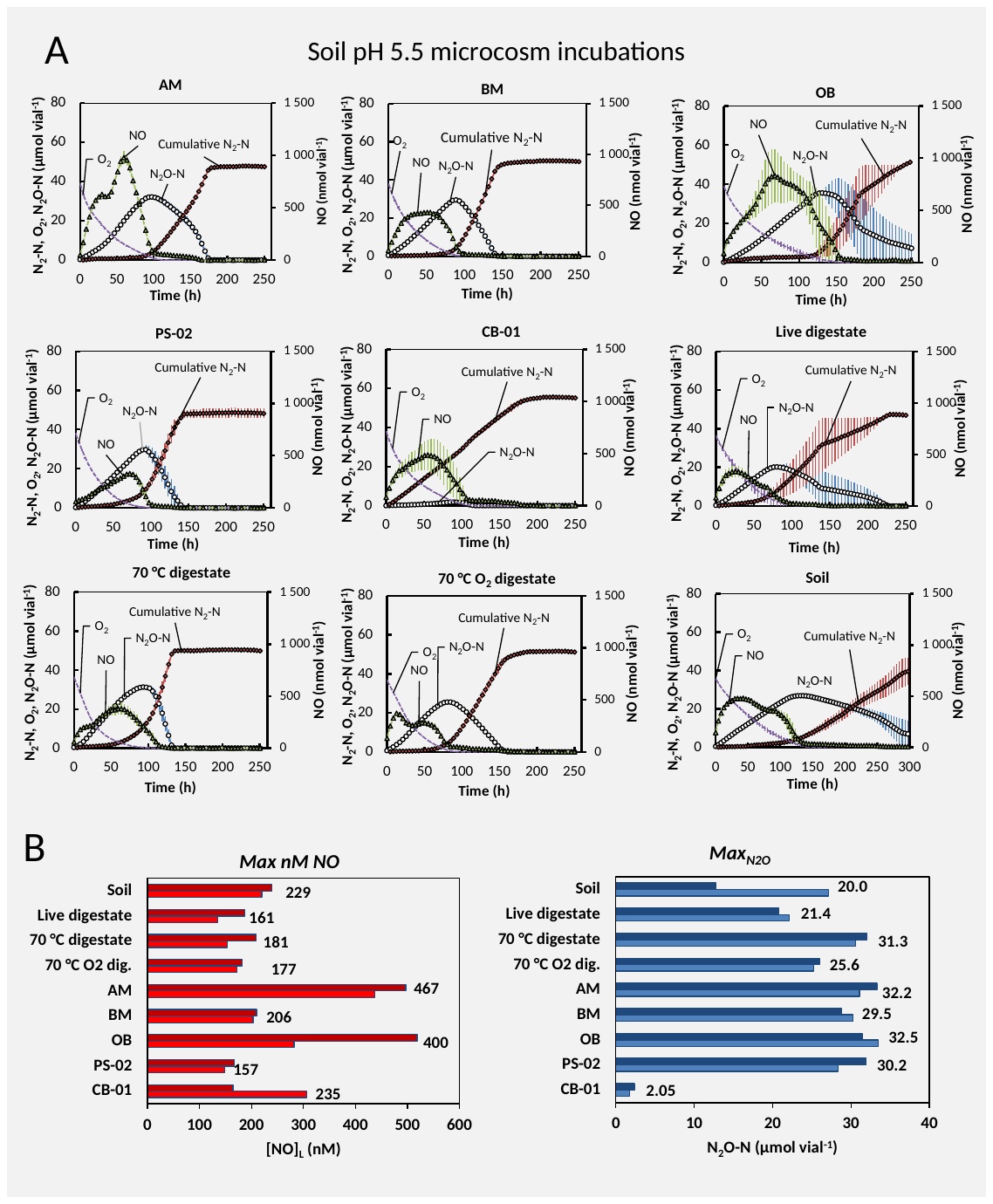

**Figure S33**: Incubation of digestate enriched with isolates (**Fig. S32**) (0.6 mL), live digestate (0.6 mL) and heat treated digestate (0.6 mL) in soil with pH=5.5 (10 g) at 20 °C**.** Panel A: kinetics of O_2_, NO, N_2_O and N_2_ throughout the incubation of soils amended with the various materials (one panel for each amendment). Average values shown (n=2). Initial oxygen (~40 µmol vial^-1^) corresponds to ~1.0 vol% in the headspace. The amounts of O_2_, NO and N_2_O are as measured, while “Cumulative N_2_-N” denotes the measured N_2_ that is corrected for leakage and losses by sampling (see Molstad et al 2007). The N_2_ and N_2_O kinetics were used to calculate the N_2_O index (***I_N2O_***), which is the area under the N_2_O- curve divided by the area under the N_2_O+N_2_ -curve for a specific time span. ***I_N2O_*** values are shown in **Fig 5** (main paper) and is a proxy for the propensity of denitrification to emit N_2_O. Panel B: peak (maximum) amounts of NO and N_2_O (results for single vials). NO is shown as nM in the liquid phase (equilibrium concentrations with measured NO in headspace), while N_2_O is shown as µmol N_2_O- N vial^-1^.

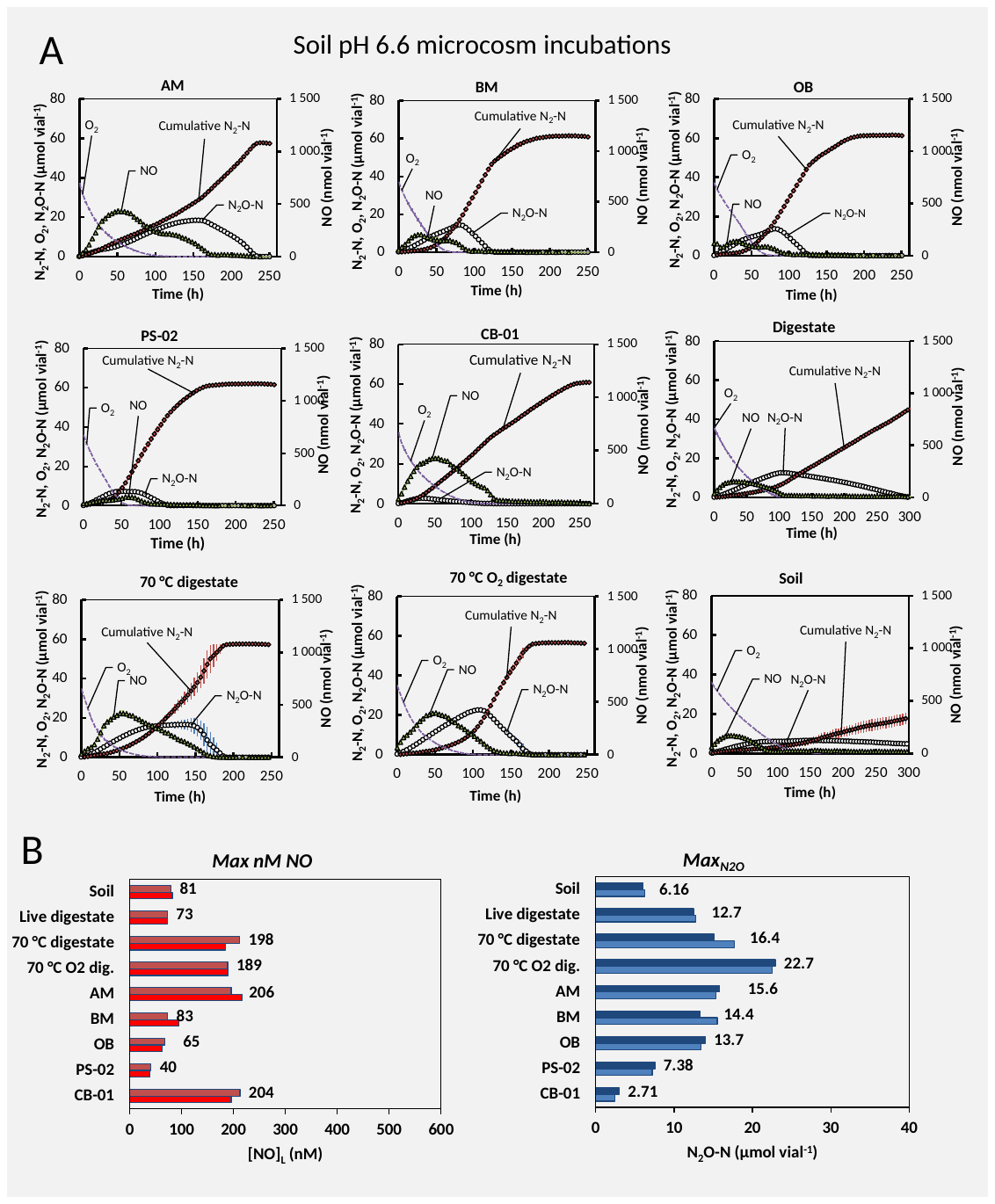

**Figure S34**: Incubation of digestate enriched with isolates (**Fig. S32**) (0.6 mL), live digestate (0.6 mL) and heat treated digestate (0.6 mL) in soil with pH=6.6 (10 g) at 20 °C**.** Panel A: kinetics of O_2_, NO, N_2_O and N_2_ throughout the incubation of soils amended with the various materials (one panel for each amendment). Average values shown (n=2). Initial oxygen (~40 µmol vial^-1^) corresponds to ~1.0 vol% in the headspace. The amounts of O_2_, NO and N_2_O are as measured, while “Cumulative N_2_-N” denotes the measured N_2_ that is corrected for leakage and losses by sampling (see Molstad et al 2007). The N_2_ and N_2_O kinetics were used to calculate the N_2_O index (***I_N2O_***), which is the area under the N_2_O- curve divided by the area under the N_2_O+N_2_ -curve for a specific time span. ***I_N2O_*** values are shown in **Fig. 5** (main paper) and is a proxy for the propensity of denitrification to emit N_2_O. Panel B: peak (maximum) amounts of NO and N_2_O (results for single vials). NO is shown as nM in the liquid phase (equilibrium concentrations with measured NO in headspace), while N_2_O is shown as µmol N_2_O- N vial^-1^.

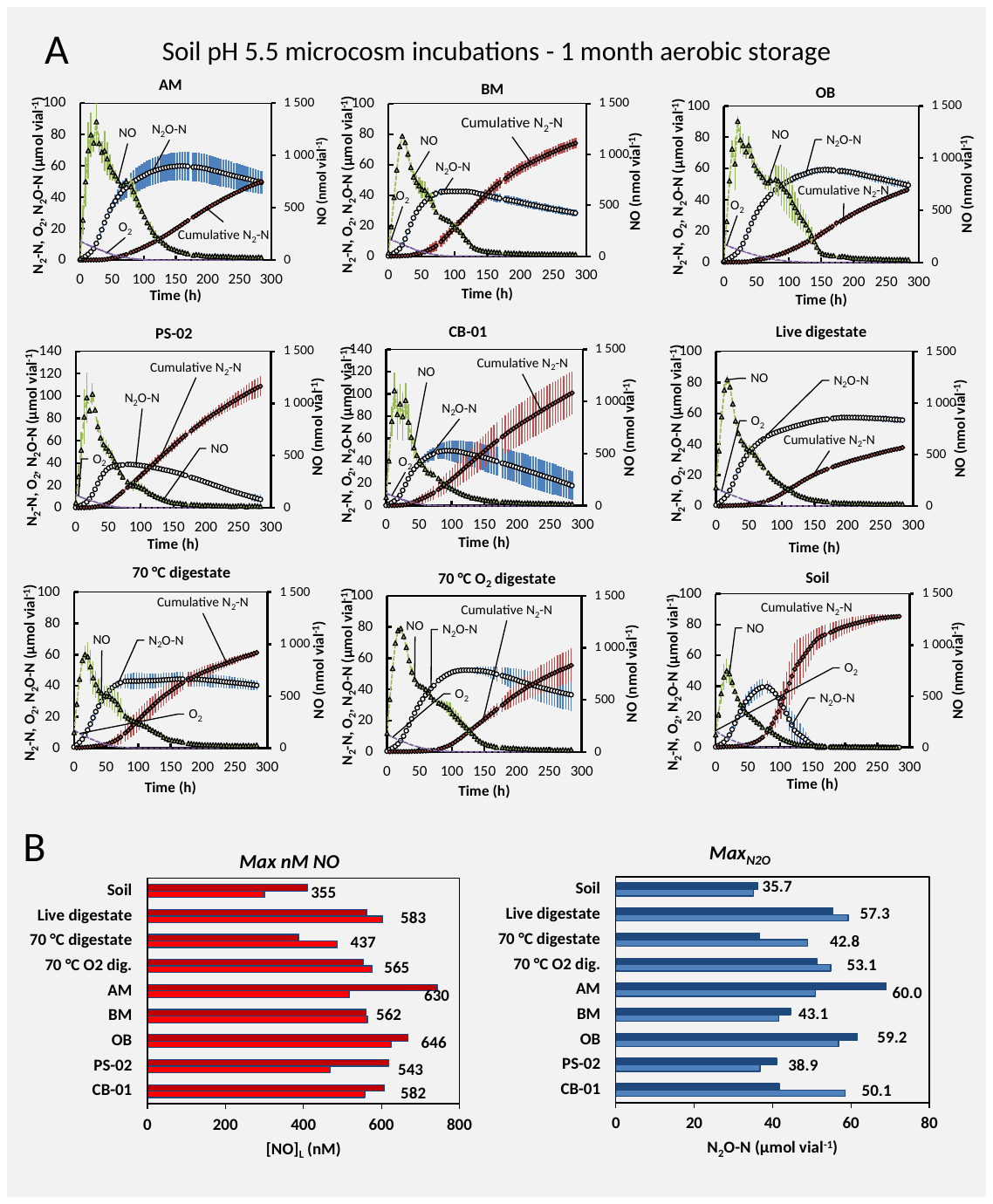

**Figure S35**: Incubation of digestate enriched with isolates (**Fig. S32**) (0.6 mL), live digestate (0.6 mL) and heat treated digestate (0.6 mL) in soil with pH=5.5 (10 g) at 20 °C after aerobic storage for 1 month (30 days) at oxic conditions (20 °C). Panel A: kinetics of O_2_, NO, N_2_O and N_2_ throughout the incubation of soils amended with the various materials (one panel for each amendment). Average values shown (n=2). Initial oxygen (~40 µmol vial^-1^) corresponds to ~1.0 vol% in the headspace. The amounts of O_2_, NO and N_2_O are as measured, while “Cumulative N_2_-N” denotes the measured N_2_ that is corrected for leakage and losses by sampling (see Molstad et al 2007). The N_2_O index (***I_N2O_***), which is the area under the N_2_O- curve divided by the area under the N_2_O+N_2_ -curve for a specific time span, was not calculable for most treatments as the experiment was not run until all available oxyanions was reduced to N_2_ or N_2_O (increasing Cumulative N_2_-N for most vials). Panel B: peak (maximum) amounts of NO and N_2_O (results for single vials). NO is shown as nM in the liquid phase (equilibrium concentrations with measured NO in headspace), while N_2_O is shown as µmol N_2_O- N vial^-1^.

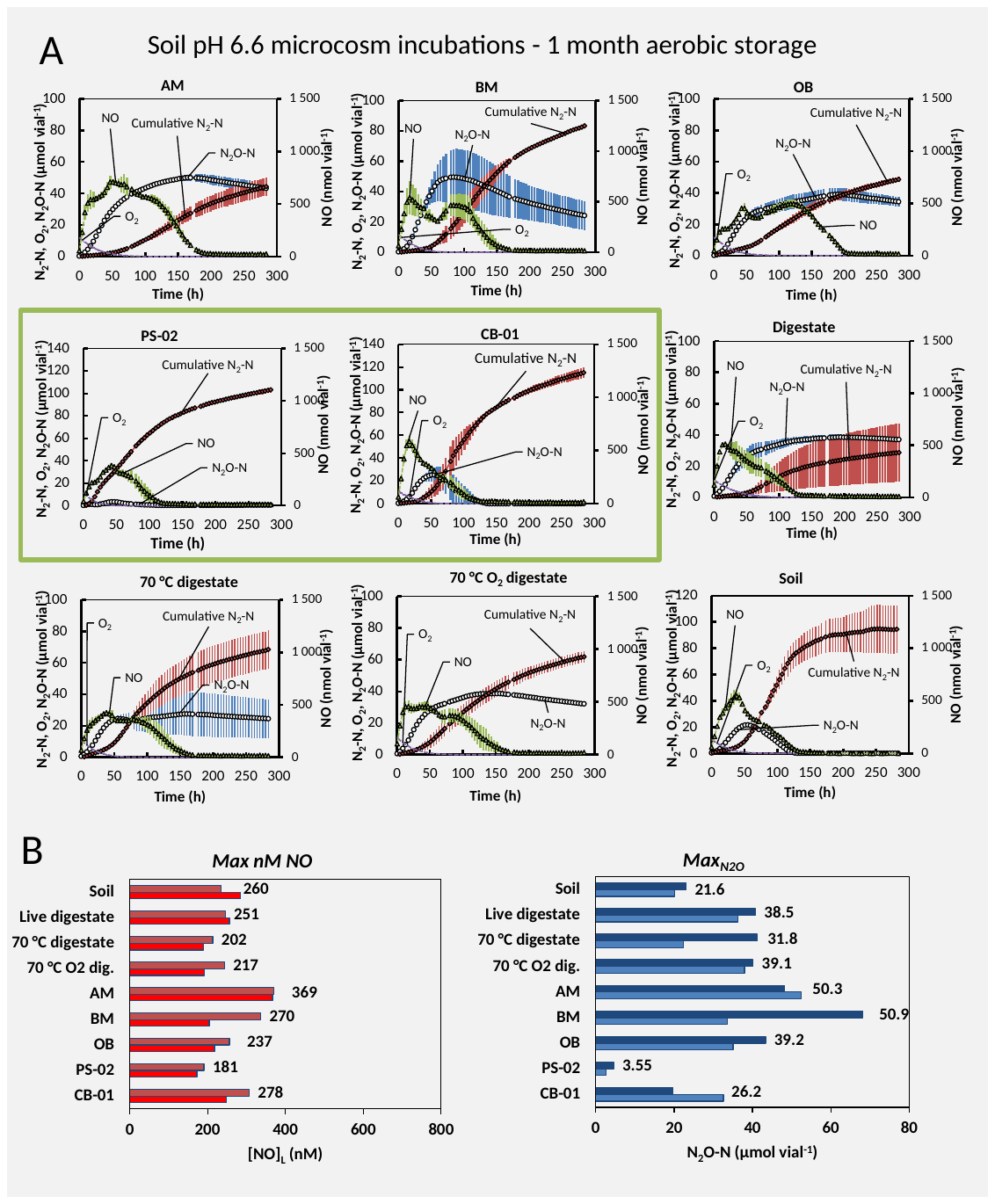

**Figure S36**: Incubation of digestate enriched with isolates (**Fig. S32**) (0.6 mL), live digestate (0.6 mL) and heat treated digestate (0.6 mL) in soil with pH=6.6 (10 g) at 20 °C were done after aerobic storage for 1 month (30 days) at oxic conditions (20°C) . Panel A: kinetics of O_2_, NO, N_2_O and N_2_ throughout the incubation of soils amended with the various materials (one panel for each amendment). Average values shown (n=2). Initial oxygen (~40 µmol vial^-1^) corresponds to ~1.0 vol% in the headspace. The amounts of O_2_, NO and N_2_O are as measured, while “Cumulative N_2_-N” denotes the measured N_2_ that is corrected for leakage and losses by sampling (see Molstad et al 2007). The N_2_O index (***I_N2O_***), which is the area under the N_2_O- curve divided by the area under the N_2_O+N_2_ -curve for a specific time span, was not calculable for most treatments as the experiment was not run until all available oxyanions was reduced to N_2_ or N_2_O (increasing Cumulative N_2_-N for most vials). The green box indicates isolates PS-02 and CB-01. Panel B: peak (maximum) amounts of NO and N_2_O (results for single vials). NO is shown as nM in the liquid phase (equilibrium concentrations with measured NO in headspace), while N_2_O is shown as µmol N_2_O- N vial^-1^. While PS-02 had a statistically significant effect on maximum N_2_O, the apparent effect of CB-01 was not statistically significant.

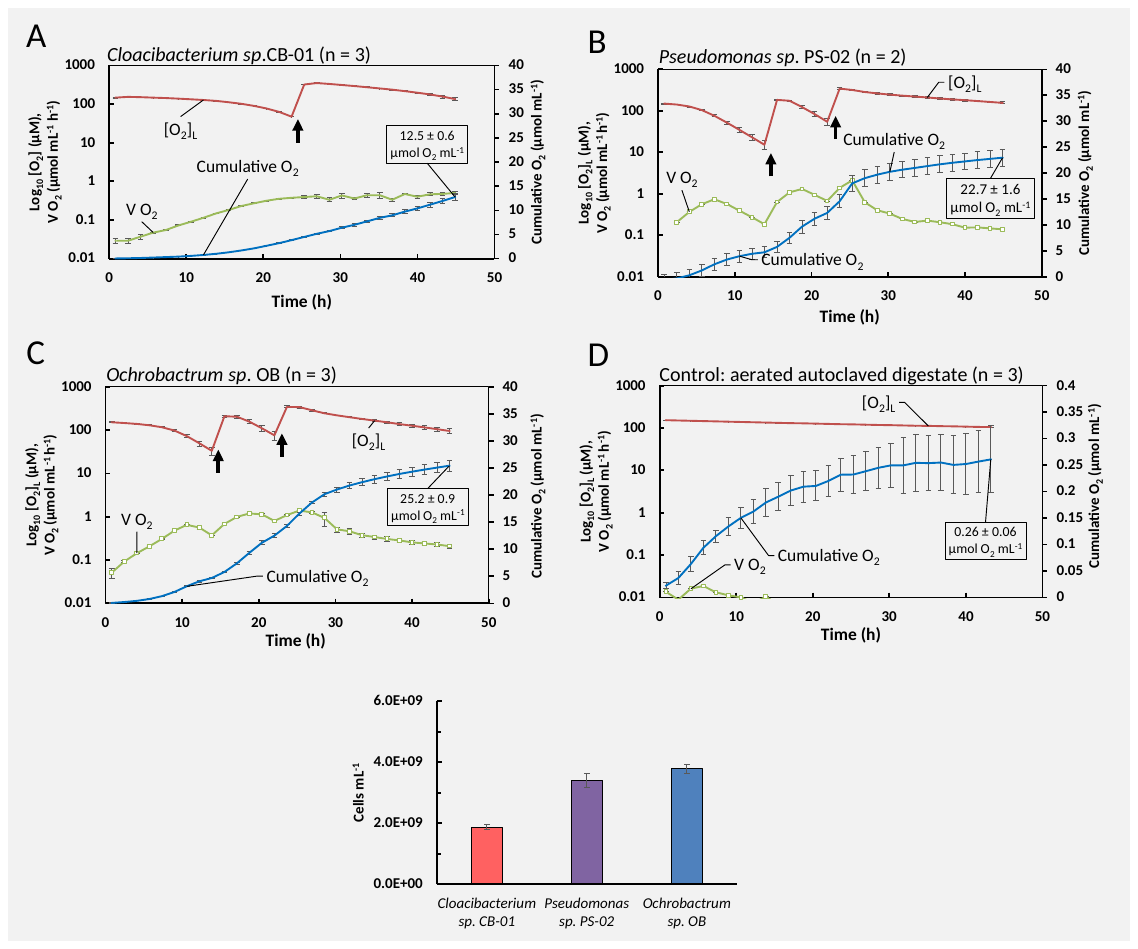

**Figure S37:** **Aerobic growth of isolated organisms in autoclaved digestate for dose response experiment.** The autoclaved digestate to be used for cultivation was pH- adjusted to 7.6 and vigorously aerated (sparging for 36 hours) before being used. The aeration was necessary because previous experiments had demonstrated substantial abiotic O_2_-consumption by oxidation of Fe^2+^ in autoclaved digestate, which would obscure the measurement of aerobic respiration by the bacteria (see **Fig. S32**). Pre-cultures of CB-01, PS-02 and OB were grown aerobically in NB medium (CB-01) and SS medium (PS-02 and OB) to OD_660nm_ 0.798, 0.379 and 0.786, respectively, and used to inoculate 120 mL vials (1 mL per vial) containing 50 mL digestate (and a magnetic bar), which were capped (butyl rubber septa), incubated at 20^o^C with vigorous stirring (600 rpm), and monitored for O_2_ concentration in the headspace. When needed, to secure oxic conditions, more O_2_ was injected. **Panels A – D**: Oxygen concentration (red), arrows = O_2_ injection. Cumulative O_2_ reduction = blue. Rate of oxygen consumption = green. **A:** *Cloacibacterium sp*. CB-01 (n=3). **B:** *Pseudomonas* sp. PS-02 (n=3). **C:** *Ochrobactrum sp*. OB (n=3). **D:** Control: no bacteria (n = 3). The cumulated oxygen consumption by each strain was used to estimate the amount of cells produced, assuming that the growth yield for all strains is the same as for *Paracocus denitrificans*, which is 30 g cell dry-weight mol^-1^ O_2_ (based on Bergaust et al (2011): 2·10^-13^ g dry-weight per cell, growth yield= 1.5·10^14^ cells mol^-1^ O_2_). The estimated amount of cell dry-weight for the three strains were 0.36 (±0.01), 0.67 (±0.04) and 0.74 (± 0.02) mg cell dry-weight mL^-1^ for CB-01, PS-02 and OB, respectively. Assuming that the three strains has the same amount of dry-weight per cell as *Paracoccus* (2·10^-13^ g cell^-1^) the estimated number of “Paracoccus equivalents” are 1.8, 3.4 and 1.9 ·10^9^ cells mL^-1^ for CB-01, PS-02 and OB, respectively.

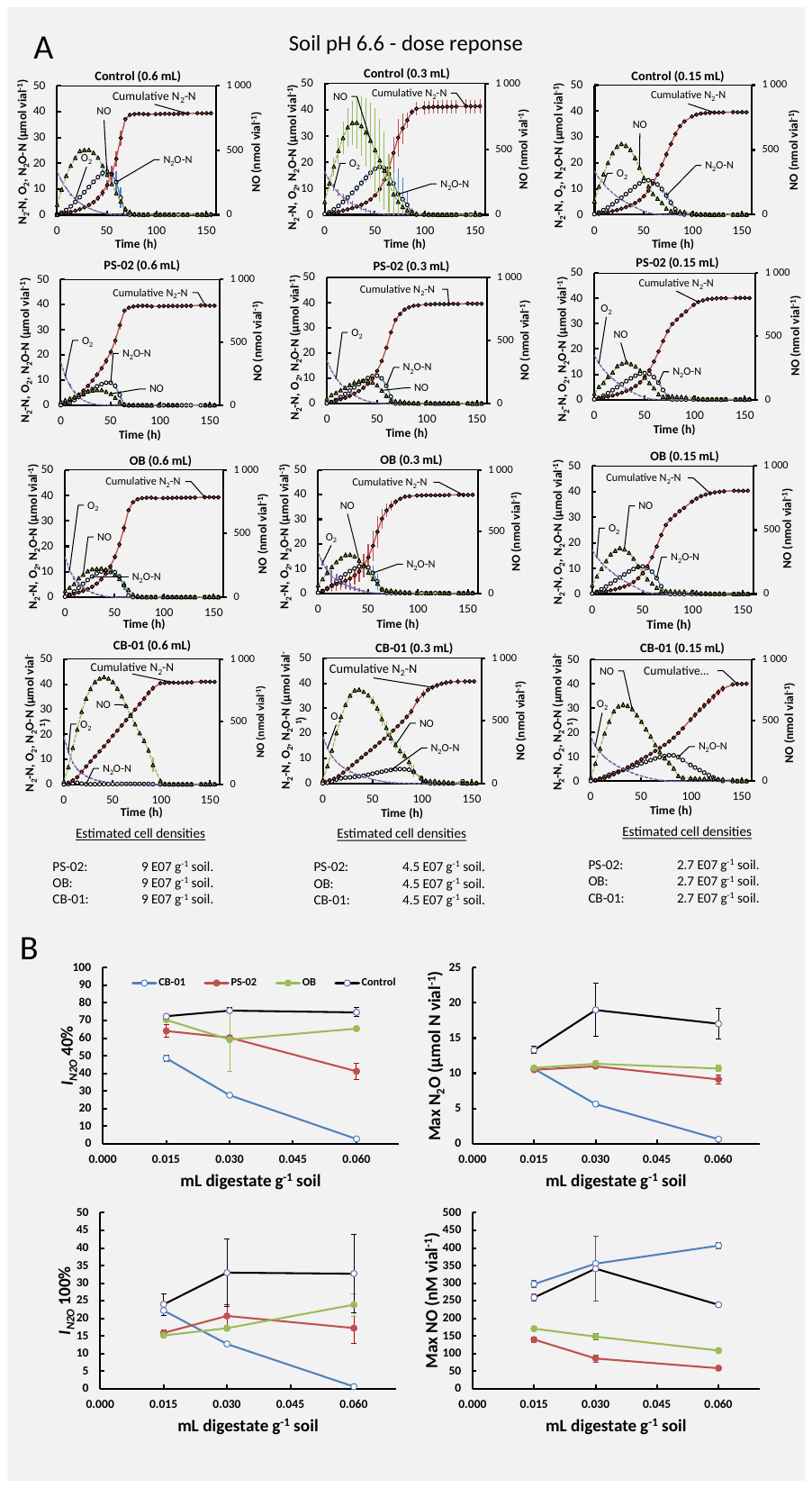
**Figure S38**: Incubation of digestate enriched with isolates OB, PS-02 and CB-01 and aerated pH adjusted autoclaved digestate (Control) in 10 g pH 6.6 soil supplemented with 25 µmol NO_3_^-^ and 0.5 mL O_2_. Panel A: kinetics of O_2_, NO, N_2_O and N_2_ throughout the incubation of soils amended with the various materials (one panel for each amendment). Average values shown, with standard deviation (n=3). Initial oxygen (~20 µmol vial^-1^) corresponds to ~0.5 vol% in the headspace. The amounts of O_2_, NO and N_2_O are as measured, while “Cumulative N_2_-N” denotes the measured N_2_ that is corrected for leakage and losses by sampling (see Molstad et al 2007). The digestate enriched with the isolates (**Fig. S37**) was diluted with sterile aerated digestate (as used in the controls) to give ~the same cell concentration per mL digestate (~2 · 10^8^ N_2_O reducing cells mL^-1^ digestate). Error bars displayed as standard deviation (n = 3 for all treatments, besides PS-02 0.15 mL with n = 2). Panel B: average peak (maximum) amounts of NO and N_2_O. NO is shown as nM in the liquid phase (equilibrium concentrations with measured NO in headspace), while N_2_O is shown as µmol N_2_O- N vial^-1^. Two ***I_N2O_*** values are shown: one for the timespan until 40% of the NO_3_^-^ -N is recovered as N_2_+N_2_O+NO-N (**I_N2O 40%_**), and one for 100% recovery (**I_N2O 100%_**).

**Table S13: Summary data for dose response experiment.** The table shows the N_2_O index values calculated for the period until 40 and 100% of NO_3_ is converted to NO+N_2_O+N_2_ (*I_N2O_* 40% and *I_N2O_* 100%, respectively), and the maximum N_2_O reached (Max N_2_O) for the dose experiment (**Fig. S38**). Average values with standard deviation are given for each treatment (n=3 replicate vials). Treatments are digestate with bacteria (CB-01, PS-02 and OB), and digestate without bacteria (Control), and 3 levels of digestate: 0.6, 0.3 and 0.15 mL digestate vial^-1^ (containing 10 g soil). The digestates with bacteria contained 0.3 mg bacterial cell dry-weight mL^-1^, hence the inoculation intensities the three levels were 18, 9 and 4.5 µg cell dry-weight g^-1^ soil. The third column for each variable shows the value expressed as % of the control value at the same inoculum intensity; significantly lower value for the bacterial treatment versus control is marked by * (p>0.05, t-test)

| **Dose** (mL vial-1) | **Strain** | ***I_N2O_* 40%** | | | ***I_N2O_* 100%** | | | **Max N_2_O (µmol N vial^-1^)** | | |
| --- | --- | --- | --- | --- | --- | --- | --- | --- | --- | --- |
|  |  | **Avg** | **St.dev** | **% of contr** | **Avg** | **St.dev** | **% of contr** | **Average** | **Stdev** | **% of contr** |
| **0.6** | CB-01 | 0.027 | 0.005 | **4 *** | 0.006 | 0.001 | **2 *** | 0.64 | 0.16 | **4 *** |
| **0.6** | PS-02 | 0.41 | 0.044 | **55 *** | 0.172 | 0.04 | **53 *** | 9.13 | 0.80 | **54 *** |
| **0.6** | OB | 0.65 | 0.012 | **88 *** | 0.239 | 0.03 | **73 -** | 10.68 | 0.53 | **63 *** |
| **0.6** | Control | 0.75 | 0.026 |  | 0.327 | 0.11 |  | 17.03 | 2.66 |  |
| **0.3** | CB-01 | 0.28 | 0.001 | **36 *** | 0.127 | 0.01 | **39 *** | 5.62 | 0.42 | **30 *** |
| **0.3** | PS-02 | 0.60 | 0.006 | **80 *** | 0.207 | 0.03 | **63 -** | 11.00 | 0.21 | **58 *** |
| **0.3** | OB | 0.59 | 0.181 | **78 -** | 0.172 | 0.01 | **52 *** | 11.34 | 0.58 | **60 *** |
| **0.3** | Control | 0.76 | 0.015 |  | 0.330 | 0.10 |  | 18.99 | 4.61 |  |
| **0.15** | CB-01 | 0.49 | 0.019 | **67 *** | 0.222 | 0.01 | **93 -** | 10.62 | 0.36 | **80 *** |
| **0.15** | PS-02 | 0.64 | 0.034 | **88 -** | 0.159 | 0.01 | **66 *** | 10.50 | 0.30 | **79 *** |
| **0.15** | OB | 0.70 | 0.010 | **97 -** | 0.152 | 0.00 | **63 *** | 10.75 | 0.35 | **81 *** |
| **0.15** | Control | 0.72 | 0.011 |  | 0.240 | 0.03 |  | 13.30 | 0.66 |  |
